# Multi-omics Integration of Microbiota Transplant Therapy in Children with Autism Spectrum Disorders

**DOI:** 10.1101/2025.10.10.681677

**Authors:** Himel Mallick, Khemlal Nirmalkar, James B. Adams, Rosa Krajmalnik-Brown

## Abstract

**Background:** Microbiota transplant therapy (MTT) is a promising avenue for the substantial improvement of gastrointestinal and behavioral symptoms in children with autism spectrum disorder (ASD). Previous work has demonstrated that microbiome and metabolite profiles of children with ASD become more similar to those of their typically developing (TD) peers following MTT.

**Methods:** To enhance a systems-level understanding of MTT in ASD children that extends beyond previously reported findings, we present a multi-omics analysis of an ASD cohort spanning 10 weeks and 2 years of follow-up after completion of MTT. We applied cutting-edge multi-omics approaches, including metagenomics, fecal and plasma metabolomics, and advanced statistical methods, including multimodal machine learning, differential network analysis, and causal mediation analysis, to extensively characterize molecular and biochemical responses before and after MTT, to identify key taxonomic, functional, and metabolite signatures associated with MTT treatment and ASD symptoms.

**Results:** Using a combination of cross-sectional and longitudinal statistical analyses and integrative machine learning techniques, we identified key meta-omic features associated with MTT. Integrated multi-omics analysis revealed that children with ASD transition to distinct biological states following MTT, clearly separated from their pre-treatment states and from TD children, as demonstrated by robust group separation and strong classification performance. Several biological signals associated with the modulation of the gut microbiome after MTT were identified, including an increase of butyrate producers such as *Faecalibacterium prausnitzii* and *Butyricimonas faecalis*; decreased fecal sulfated primary bile acid, chenodeoxycholic acid sulfate; decreased secondary bile acid, glycolithocholate sulfate; and increased sarcosine and iminodiacetate in plasma after 10 weeks of MTT compared to baseline. Differential network analysis revealed hub species, including *Prevotella copri*, *Ruminococcus callidus*, and *GGB9633 SGB15091*, as differentially connected 2 years after completion of MTT compared to baseline. Mediation analysis uncovered several key players as mediators of symptoms, including *Alistipes ihumii*, *Ruminococceae*, amino acid biosynthesis, bile acids, long-chain fatty acids, and cysteine-glutathione disulfide, among others.

**Conclusions:** This study provides one of the first comprehensive analyses of multi-omic features underlying host–microbiome interactions associated with MTT in children with ASD. It offers further evidence that fusing data across diverse molecular modalities at pre- and post- treatment time points can illuminate the potential of MTT in neurodevelopmental disorders. These findings could advance microbiome-based immunomodulatory therapies and multi-omics strategies to restore gut microbiota in children with ASD, while aiding in the discovery of novel biomarkers predictive of treatment response.

## Introduction

According to the Centers for Disease Control and Prevention (CDC), USA, 1 in 31 children is diagnosed with autism spectrum disorder (ASD) each year, and the rate is steadily rising^1^. ASD is a multifactorial neurodevelopmental disorder that affects normal brain development and involves deficits in social communication, restricted interests, and repetitive behaviors^2–5^. The etiology of ASD is believed to be highly complex and heterogeneous, but the gut microbiome, the gut metabolome, and their interactions with the host have been implicated in ASD, as about 40% of children with ASD suffer from chronic gastrointestinal (GI) symptoms, which correlate with ASD severity^6–9^.

Microbiota tranplant therapy (MTT) is a promising treatment for improving GI and ASD-related symptoms in children with ASD. Our previous work demonstrated that the microbial and metabolite profiles of children with ASD became more similar to those of their typically developing (TD) peers following MTT^10–14^. Specifically, we conducted a successful Phase I open-label clinical trial of MTT at Arizona State University, where MTT reduced GI symptom severity by ∼80% and ASD symptoms by ∼24% by the end of treatment, with nearly a 50% reduction in ASD-related symptoms two years post-treatment^10,15^. MTT involved 10 weeks of treatment, including pre-treatment with vancomycin for two weeks, a bowel cleanse, a stomach acid suppressant, and daily fecal microbiota transplant (FMT) for seven to eight weeks (**Fig. 1**). Previous analyses of this cohort, using 16S rRNA gene amplicon sequencing, shotgun metagenomic sequencing, and metabolomics evaluations, found treatment-associated changes in microbial and metabolomic profiles, along with improvements in ASD and GI symptoms.

**Figure 1.**
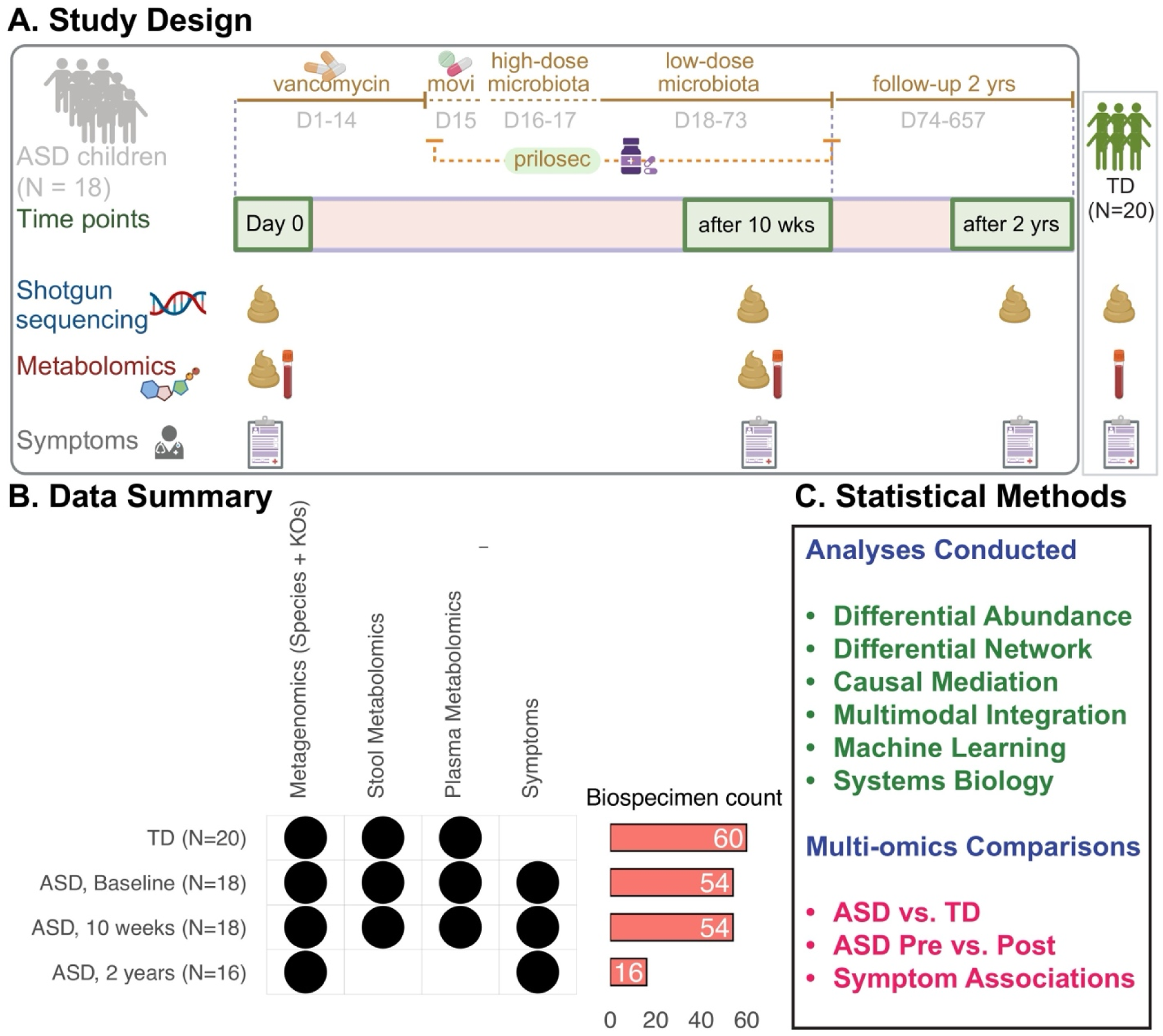
Study design and clinical trial timeline describing the collection of stool and plasma samples for multi-omics measurements and assessments of GI and ASD symptoms. A. The timeline includes the baseline period, a 10 week period following microbiota transplant therapy (MTT), and a 2-year follow-up observation. The modalities include microbiome shotgun sequencing, stool and plasma metabolomics, and symptom assessments. B. At baseline, multi-omics measurements are available for N = 20 TD children and N = 18 ASD children. Post-baseline, multi-omics measurements are available for N = 18 ASD children at 10 weeks and N = 16 ASD children at the 2-year follow-up. Symptom data are available for both baseline and post-baseline time points for ASD children. Bar plots show the total number of biospecimens available for analysis. C. Statistical methods include multimodal machine learning, differential network analysis, differential abundance analysis, similarity network fusion, and causal mediation analysis across time points and between ASD and TD groups (Methods).

Although substantial progress has been made in identifying individual microbes and metabolites associated with ASD^9^, their complex interactions remain poorly understood. While MTT has shown promise in improving GI and behavioral symptoms in children with ASD, the mechanisms through which individual microbes, functions, and metabolites exert their effects remain largely unknown^7,9,11^. Moreover, previous analyses either did not explore functional profiles in sufficient detail or did not perform integrative analyses. Due to the complex nature of the gut-brain axis— the communication channel between the gut microbiome and the brain—integrative approaches are needed to identify new preventive and therapeutic targets for ASD. This is driven by the hypothesis that integrated analysis can lead to more powerful discoveries of multi-omics biomarkers associated with MTT response and ASD outcomes.

Accumulating evidence suggests that traditional single-layer analyses of multi-omics data are less powerful than integrative approaches, as they do not account for specific inter-layer interactions between vastly different biological data types^16,17^. Only in recent years—due to advances in computational and statistical methods—has it become feasible to perform systems- level assessments of multi-omics functional architecture^9,18^. However, microbiome analysis in ASD remains limited by the lack of appropriate methodologies for jointly analyzing longitudinal multi-omics data. Most published ASD microbiome studies are either cross-sectional or treat different omics types independently, without integration^10,12–15^. Consequently, multi-omics analysis of MTT provides a unique opportunity to uncover deeper insights into the microbiome’s role in ASD in a more holistic and cost-effective manner.

Previous studies, including Morton et al.^9^, have explored associations between the gut microbiota and behavioral outcomes in ASD, revealing global microbiome–phenotype patterns across cohorts. However, these efforts primarily relied on univariate analyses, lacked mechanistic interpretation, and did not incorporate multi-omics integration or network-level modeling. To address these gaps, we applied a high-dimensional mediation framework that jointly integrates microbial taxa, functions, and metabolites to identify molecular mediators of treatment response. Coupled with network-based analysis, our approach enables a systems- level view of how coordinated shifts in the microbiome and metabolome contribute to symptom improvement in ASD.

This study thus addresses a longstanding gap in ASD microbiome research by jointly analyzing multiple omics profiles - including taxonomic and functional profiles from the microbiome, along with stool and plasma metabolites—in ASD and TD populations at baseline, and in ASD at post- treatment time points. Leveraging the longitudinal study design of the Phase I open-label ASU- MTT clinical trial, we employed multimodal integration, network analysis, and causal mediation analysis to link novel microbial data types to ASD outcomes before and after treatment. We identified multi-omics features positively and negatively associated with ASD and determined the most confident associations for experimental validation, with the long-term goal of harnessing these relationships to improve MTT response and ASD outcomes. This framework for inferring mechanisms from population-scale multi-omics studies is pivotal for advancing future mechanistic research and enhancing our understanding of the microbiome’s role in ASD.

## Results

This multi-omics study analyzes a Phase I open-label trial of MTT in children with ASD, previously characterized using 16S rRNA gene amplicon sequencing, shotgun sequencing, plasma metabolomics, and stool metabolomics. The original study (ASU-MTT) included 20 age- and gender-matched typically developing (TD) children and 18 children with ASD, aged 7–16 years (**Fig. 1A-B)**. It involved three data collection points: baseline, 10 weeks after MTT, and 2 years after MTT (with fecal samples provided by 16 of the 18 participants). Participant characteristics, along with medical and dietary history, were recorded as detailed elsewhere^10,15^. No dietary changes were made during the MTT treatment, and dietary records were updated two years post-treatment. We assessed microbial species, functional profiles (Kyoto Encyclopedia of Genes and Genomes (KEGG) Orthologs [KOs]), and plasma and stool metabolites across pre- and post-treatment time points in children with ASD, in addition to baseline multi-omics measurements in TD children. These features were used to conduct a comprehensive multimodal analysis, including differential network analysis, causal mediation analysis, and integrative machine learning. Published state-of-the-art multimodal tools were used throughout these analyses (**Fig. 1B–C**). The following sections expand on these analyses, with additional technical details provided in **Methods**.

### Multi-omics signatures differentiate pre-post ASD children across time points

We first applied unsupervised principal coordinates analysis (PCoA) to assess whether multi- omics profiles naturally separate by group, without using outcome labels to guide the analysis. Microbiome data were centered log-ratio (CLR) transformed, and metabolomics data were log- transformed, enabling appropriate compositional handling and variance stabilization while preserving biological gradients (**Methods**). Using Euclidean distance consistently across modalities, significant group differences were detected across all omics layers (PERMANOVA P < 0.05), though the degree of separation varied (**Fig. 2**). Species-level microbiome (**Fig. 2A**) and KO functional profiles (**Fig. 2B**) showed moderate clustering, while fecal metabolites (**Fig. 2C**) exhibited weaker separation, consistent with the low R² value (0.045). Notably, plasma metabolites (**Fig. 2D**) revealed a clear separation of ASD baseline (red) from all other groups (R² = 0.07). After MTT, ASD children (green and purple) shifted closer to the TD group (blue), suggesting that plasma metabolomic profiles became more normalized following treatment, as previously reported^19^. Similar group separation was observed in the integrated dataset (**Fig. 2E**; R² = 0.07), underscoring the discriminatory power of multi-omics integration. While we observed partial separation of baseline and 10-week samples without the 2-year data (**Fig. 2E)**, after including the 2-year data, the 2-year group became distinct from the others, likely due to changes in taxonomic composition and potential functional genes (**Fig. 2F**; R² = 0.25). However, the apparent separation of the 2-year group may also partly reflect the absence of metabolomics data for this time point, which could influence the integrated ordination. These results indicate that while individual omics layers capture distinct aspects of group differences, integration provides a holistic view of biologically meaningful shifts across ASD and TD groups over time.

**Figure 2.**
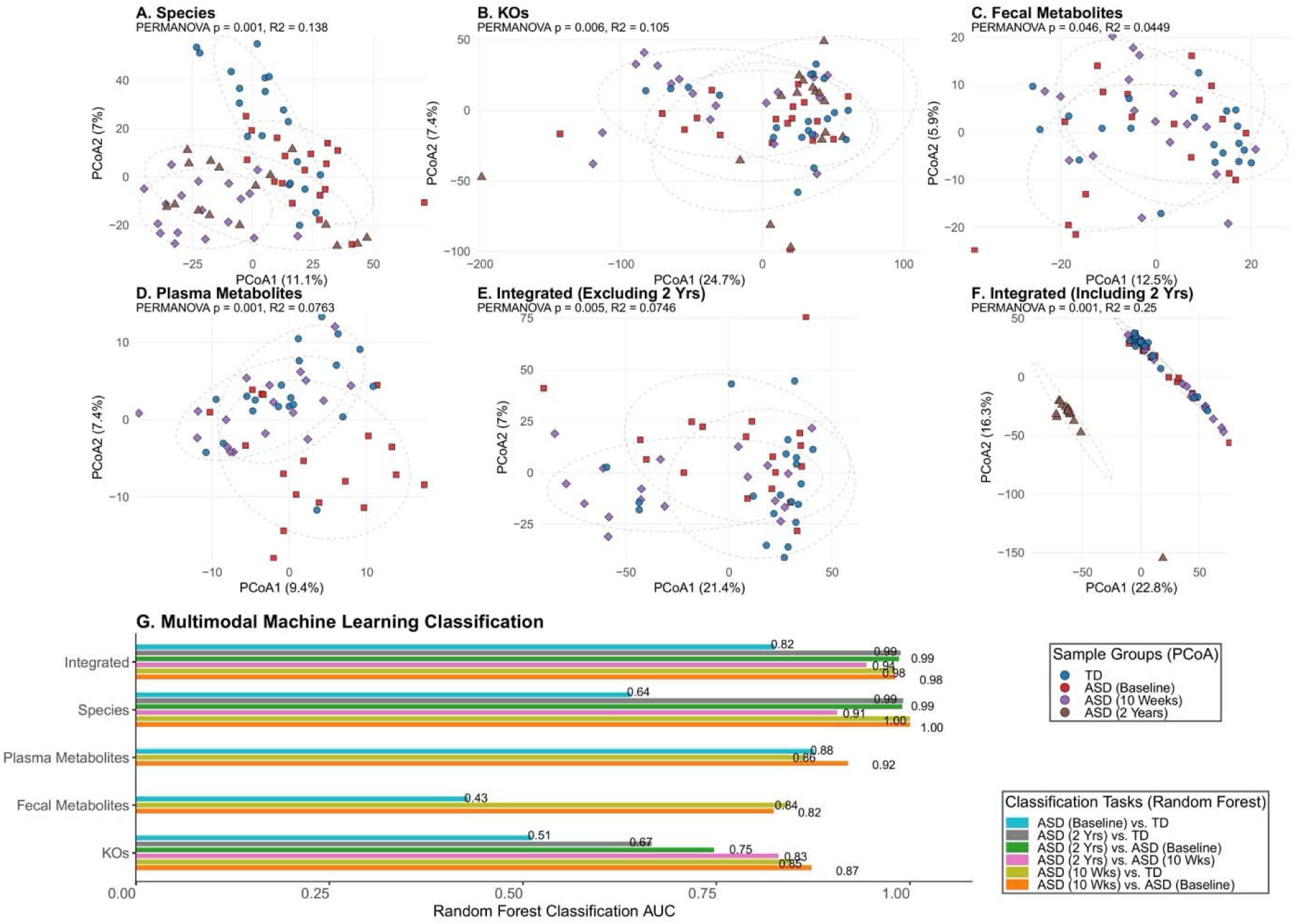
Multimodal ordination and classification performance. A–D. Principal coordinates analysis (PCoA) plots for species (**A**), KOs (**B**), fecal metabolites (**C**), and plasma metabolites (**D**), respectively. Microbiome layers were CLR-transformed, and metabolomics layers were log-transformed prior to computing Euclidean distances. **E–F** Integrated PCoA on concatenated, scaled data across all modalities, showing separation by group, excluding 2 years (**E**) and including 2 years (**F**). Variance explained and PERMANOVA p-values are shown in each panel. **G**. Random forest classification performance across modalities and integrated data, displayed as area under the receiver operating characteristic curve (AUC) for different pairwise comparisons. Random forest classifiers were trained on plasma metabolites, stool metabolites, microbial species, microbial functional profiles (KOs), and their combinations to distinguish ASD children before and after MTT at 10 weeks and 2 years. Classifiers were also trained to distinguish ASD from TD children at both baseline and post-baseline. Model training was performed using five-fold cross-validation using *IntegratedLearner*^16^. AUC refers to the area under the receiver operating characteristic (ROC) curve.

To assess whether differences in metabolite or microbial composition could classify ASD children based on their pre- and post-treatment status, we trained random forest (RF) classifiers on metabolic and microbial species profiles, both separately and in combination^16,20,21^. Classification performance was evaluated using five-fold cross-validation. All classifiers significantly outperformed random chance in distinguishing pre- and post-MTT ASD children, with Area Under the Receiver Operating Curve (AUC) values ranging from 0.75 to 1.0 (**Fig. 2F**). AUC values close to 1.0 indicate high sensitivity with a low false-positive rate, while a value of 0.5 represents random performance. The integration of metabolite and microbial data did not significantly improve classification accuracy compared to using either dataset alone, likely due to the high overlap of information between microbial and metabolomic profiles.

Notably, AUC values were substantially higher for the two-year vs. baseline comparison (except for KOs), suggesting more pronounced biological divergence over extended follow-up. To further investigate these shifts, we re-trained the integrated model using presence-absence– based features derived from microbial species and KO profiles. This approach was motivated by recent evidence highlighting the ecological importance of prevalence-based representations in characterizing microbial community structure^22,23^. The resulting models maintained similar cross-validation prediction accuracy (**Fig. S1**), reinforcing the robustness of the observed patterns.

Taken together, these findings demonstrate that multi-omics signatures are highly effective in classifying pre- vs. post-MTT ASD states, with classifiers performing better in distinguishing pre- and post-treatment samples than in separating ASD from TD at baseline (**Figs. 2F;S1**). This suggests that while baseline ASD profiles differ from TD profiles, these differences are relatively subtle compared to the marked shifts induced by MTT. Although some post-treatment changes may reflect partial restoration toward TD-like profiles, the improved separability also points to the emergence of a new distinct multi-omics profile, different from baseline ASD and TD groups. This treatment-associated biological state, rather than representing complete normalization, may reflect transitional or compensatory adaptations, microbial remodeling, or new host- microbe interactions that align with clinical improvements observed over time.

### Multivariable association delineates treatment-associated multi-omics features in ASD children

To identify robust MTT-associated features, we extracted feature importance scores from the integrated model and visualized the top predictive signals in an omics-specific manner. The analysis revealed distinct sets of discriminative features across microbial taxa, KOs, and fecal and plasma metabolites when comparing the 10 week time point to baseline (**Fig. 3)**. Importantly, the most predictive microbial and functional features at 10 weeks were largely different from those identified at the 2-year follow-up (**Fig. S2)**, suggesting that the biological signals associated with treatment response evolve over time. It’s important to note that plasma and fecal metabolite data were not available for the 2-year time point. Although plasma and fecal metabolite data were not available for the 2-year time point, this divergence in microbial and functional features highlights the dynamic nature of MTT-induced changes and supports the notion of long-term remodeling of the gut microbiome rather than a transient or uniform shift.

**Figure 3.**
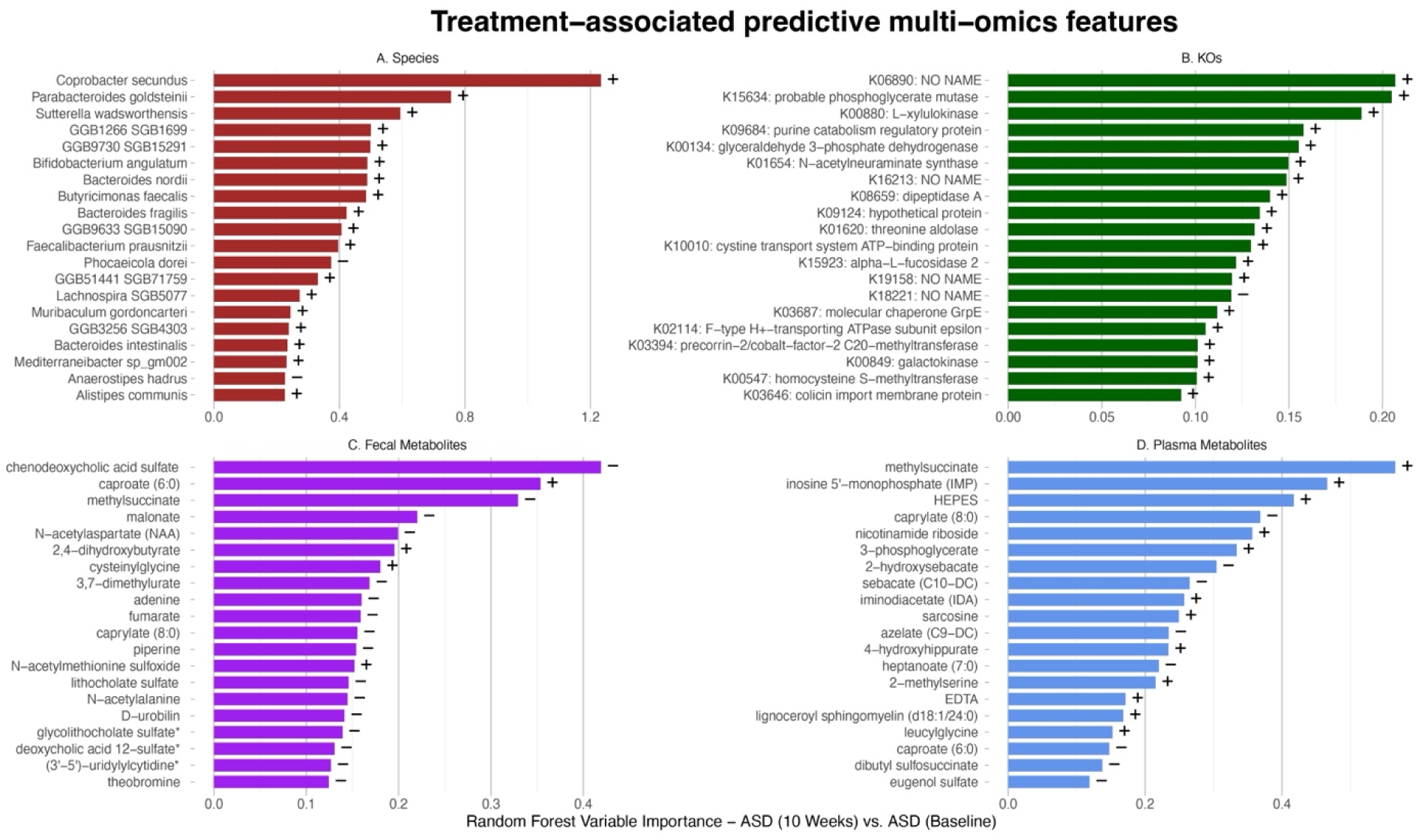
**Feature selection results from the cross-validated integrated machine learning model comparing the 10 week MTT group to baseline in ASD children**. The bar plots depict feature importance scores for the top 20 features per layer from the integrated model: microbial species (**A**), microbial KOs (**B**), fecal metabolites (**C**), and plasma metabolites (**D**). Positive and negative associations are indicated by + and – signs. Classification performance was evaluated using five-fold cross-validation (performance reported in Fig. 2F).

The key predictive features for 10 weeks vs. Baseline, includes increased sulfated bile acids from plasma and feces, as well as increase in relative abundance of beneficial, butyrate producers such as *Bifidobacterium angulatum*, *Faecalibacterium prausnitzii*, and *Butyricimonas faecalis* **(Fig. 3).** Notably, *Butyricimonas faecalis* remained elevated at the 2-year time point (**Fig. S2**). However, most bacterial taxa that increased at 10 weeks (**Fig. 3A**) significantly declined by 2 years (**Fig. S2**), while others, like *Roseburia faecis*, newly emerged—suggesting that MTT leads to a long-term, adaptive shift in gut microbial composition^10^.

Interestingly, several beneficial and probiotic bacterial species, including *Bifidobacterium*, *Bacteroides intestinalis*, *Butyricimonas faecalis*, and *Coprobacter*, significantly increased after 10 weeks of MTT compared to baseline in ASD children (**Fig. 3A**). Notably, *Bacteroides fragilis*, a bacterium whose role in ASD has been debated, also increased. Some researchers have suggested a beneficial role for *B. fragilis* in ASD^24,25^, while others have argued against it^26,27^. Additionally, sulfur-reducing *Sutterella* was dominant after 10 weeks of MTT vs. Baseline (**Fig. 3A**), and remained at increased levels after 2-years of MTT (**Fig. S1**), which contrasts with other studies involving ASD^28–30^. Interestingly, *Prevotella copri clade A* significantly decreased after 2- years of MTT vs. Baseline (**Fig. S2**), consistent with a previous report at the genus level for the same samples using shotgun metagenomics^14^, but in contrast to 16S rRNA sequencing^10^ - suggesting that these differences could arise due to differences in primer, sequencing technology or taxonomy assignment methods for shotgun and 16S rRNA gene sequencing. At 10 weeks, we did not observe any significant difference *for P. copri clade A* and any other species of *Prevotella*.

At the KO level, the tetracycline resistance gene K18221 (tetracycline 11a-monooxygenase; tetX) showed a decrease, while glucose metabolism-related genes such as K00134 (glyceraldehyde 3-phosphate dehydrogenase; GAPDH) and K15634 (phosphoglycerate mutase; PGM), along with some cell membrane-associated genes, exhibited an increase post-MTT **(Fig. 3B)**. KO profiles differed at the 2-year MTT timepoint compared to baseline, and and notably, K01130 (arylsulfatase) was significantly decreased after 2 years of MTT (**Fig. S2**). K01130 (arylsulfatase) can convert aryl sulfate into sulfate and phenol, and high concentration of phenol in blood and urine have been found in ASD children^31,32^. It has been hypothesized that early phenol exposure may be linked to an increased risk of developing ASD^33,34^. Most of these associations were also recapitulated when using microbial prevalence data, suggesting that many abundance-based associations are also reflected at the level of presence-absence (**Fig. S3**).

Regarding fecal metabolites, we observed an increase in the medium-chain fatty acid caproate (6:0), while caprylate (8:0) and several sulfated bile acids, such as chenodeoxycholic acid sulfate (a primary bile acid) and glycolithocholate sulfate (a secondary bile acid), significantly decreased after 10 weeks of MTT compared to baseline **(Fig. 3C)**. Interestingly, glycolithocholate is derived by gut bacteria from chenodeoxycholic acid and both are sulfated.

Since bile acids are crucial for nutrient digestion and absorption, and sulfation aids in detoxification and excretion^35,36^, the reduction in these sulfated bile acids post-MTT suggests a diminished excretion of these important compounds. In plasma metabolites, caproate and caprylate decreased, while inosine monophosphate, iminodiacetate (IMP), and sarcosine increased after 10 weeks of MTT compared to baseline, consistent with the findings reported by Kang et al.^10,15^ **(Fig. 3D)**.

While our primary analysis focused on multimodal integration, we also conducted a univariate analysis to isolate specific microbial changes that were differentially abundant in ASD compared to TD, or between ASD pre- and post-MTT time points (**Methods**). Several significant findings were noted (**Figs. S4-S7; Tables S1-S4**). For example, we found that *Prevotella sp P4.51|SGB1666* signficnantly increased after 10 weeks of MTT compared to baseline, which is consistent with previous findings using 16S rRNA gene sequencing^15^. Another example is *p*- cresol, a fecal metabolite, was significantly reduced after 10 weeks of MTT compared to baseline (**P = 0.03**, **Table S4**). Although marginally significant, the result is particularly intriguing, as *p*-cresol has been consistently reported to be elevated in individuals with ASD and is linked to increased autism severity^37–39^. Additionally, a study demonstred that administration of *p*-cresol in mice exhibited autism-like behavior^40^, but fecal microbiota transplant from healthy mice lead to normalization of symptoms^40^.

Notably, features identified through univariate differential abundance analysis (**Figs. S4–S7; Tables S1–S4**) showed minimal overlap with those from the integrated model. As before, many of these associations were also supported by prevalence-based representations (**Figs. S4–S7)**. As is well-documented in the literature^41–45^, univariate analysis poses challenges in distinguishing MTT-associated, biologically informative microbes from those involved in covariation relationships. These issues arise because microbial feature abundances are inherently linked through ecological interactions, which may result in spurious associations. While some microbes are truly related to MTT, others may only appear associated due to indirect correlations with the treatment. This lack of significant overlap supports the observation that multivariable analysis can effectively filter out features that may be erroneously reported as associated due to confounding relationships with truly associated features, a result often driven by compositionality and feature-feature interactions.

### Differential network analyses find strong ASD–microbiome links differentially connected across pre- and post-treatment time points

Above, we demonstrated that multimodal integration uncovers individual features from multivariable analysis that accounts for confounding feature-feature associations. Unlike univariate analysis, which examines microbial associations on a per-feature basis without considering the rest of the microbial features, multivariable analysis acknowledges that microbial features, such as species, do not function independently but interact within a complex, interdependent ecological network. These microbial networks offer insights into various interspecies and intercellular relationships, including mutualism (win-win), competition (lose- lose), parasitism or predation (win-lose), commensalism (win-zero), and amensalism (zero- lose). Understanding these relationships is crucial for unraveling the function, structure, and dynamics of microbial communities. Although analyzing a single microbial association network can provide insights into the general organizational structure of a microbial community, it is often desirable to investigate how microbial associations change across different conditions. Differential network analysis can directly assess network differences between groups and pinpoint specific microbe-microbe associations that significantly change across conditions, providing concrete hypotheses for follow-up biological perturbation experiments.

To examine whether MTT has a concrete and significant impact on the inherent connectivity of the microbial community and whether MTT alters microbe-microbe interaction networks across pre- and post-treatment periods in ASD children, we first constructed individual microbial networks independently for pre- and post-MTT time points. Next, we assessed differences between these individual networks by considering both global and local measures of differential connectivity (**Methods**). At the 10 weeks time point, a particularly well-connected representative community emerged, consisting of three species from the genera Ruminococcaceae *GGB9633 SGB15093*, and *Akkermansia* (**Fig. 4A).** Although the precise functional interactions among these species remain unclear, their strong interconnectivity suggests a shift toward a more stable and potentially beneficial gut microbiome following MTT. Another prominent hub observed at both baseline and the 2-year post-treatment time point was a three-member community comprising *Prevotella copri*, *Ruminococcus callidus*, and Ruminococcaceae *GGB9633 SGB15093*. While this subnetwork remained tightly connected over time, the surrounding microbial associations shifted markedly (**Fig. 4B**). Notably, *Prevotella copri* and *Ruminococcus callidus* also showed differential connectivity at the 2-year follow-up compared to the 10-week time point (**Fig. S8**). Additional network changes between 10 weeks and 2 years included the emergence of a newly connected subnetwork involving *Bacteroides cellulosilyticus*and Ruminococcaceae *GGB9634 SGB15093* (**Fig. S8**).

**Figure 4.**
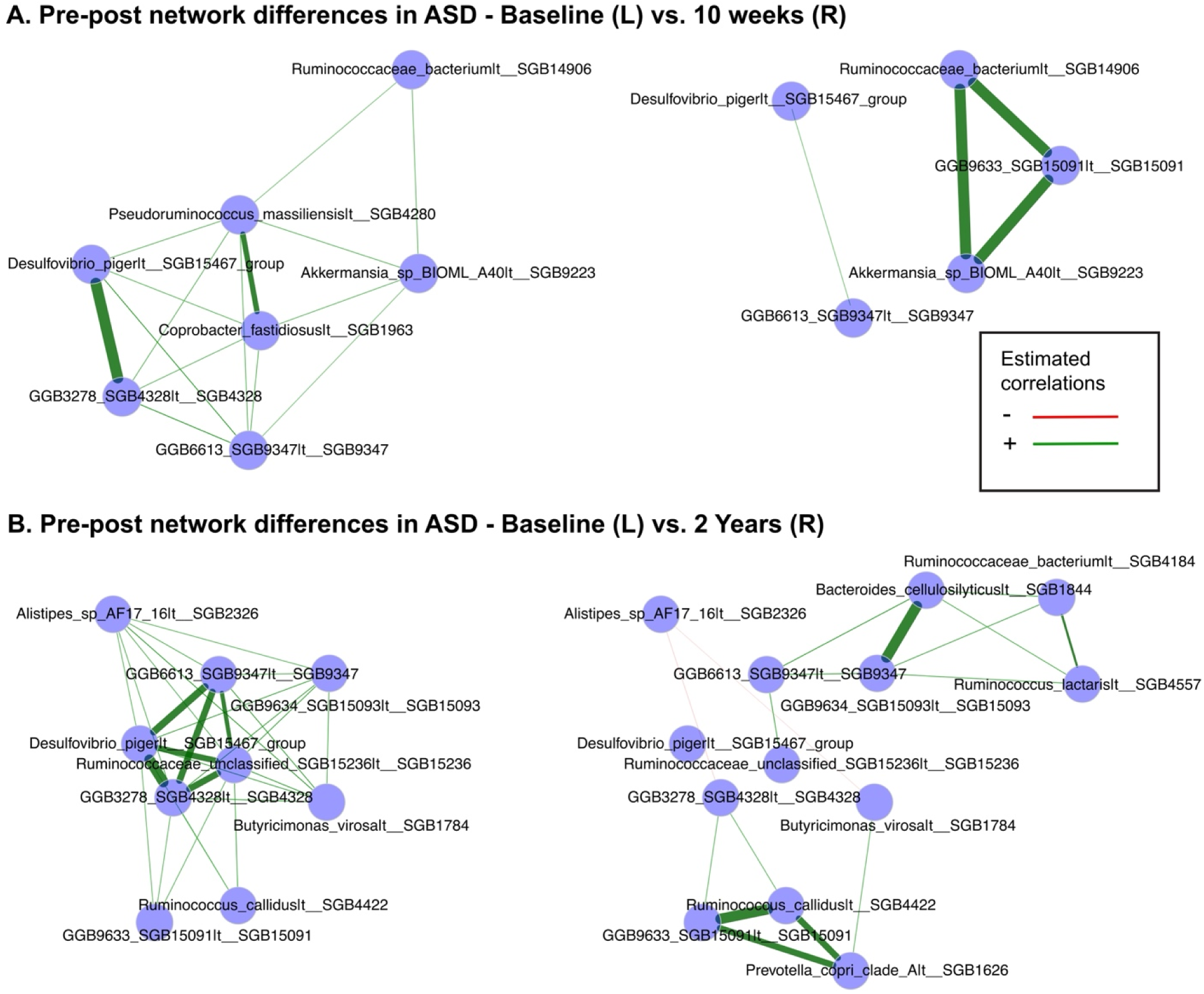
**Differential network analysis of microbiome species identifies differentially connected microbial species pre- and post-MTT**. In the differential network, several hubs are detected that are strongly associated post-baseline but weakly associated at baseline, and vice versa. For visualization purposes, only the differentially associated species are plotted. **A**. A three-member community of species from the genera *Ruminococcaceae*, GGB9633, and *Akkermansia* remained tightly connected 10 weeks after MTT treatment, but much of the rest of the network changed. **B**. A three-member community of *Prevotella copri*, *Ruminococcus callidus*, and *GGB9633 SGB15091* remained tightly connected 2 years after MTT treatment, but much of the rest of the network changed. The thickness of the edges represents the strength of association, with a thick line indicating a very strong correlation and a thin line indicating a weak correlation. L: Left side of the figure; R: Right side of the figure. Full results are provided in **Table S5**.

### MTT treatment contribution to ASD symptoms improvements waspotentially mediated by microbes and metabolites

Previous studies have demonstrated that MTT has a systematic and long-lasting effect on the gut microbiome of children with ASD, manifesting as significant improvements in both GI and autism-related symptoms^10,15^. While our earlier analyses highlighted multimodal associations between microbial features and MTT, emerging research suggests that the microbiome may also act as a *mediator* - whereby the exposure (MTT treatment) partially influences the outcome (symptom improvement) through changes in microbiome composition. In other words, changes in the microbiome due to MTT may contribute to improvements in ASD symptoms. These relationships can be investigated using statistical mediation analysis, which quantifies the proportion of the treatment effect on the outcome that is mediated through microbiome features, compared to the direct (unmediated) effect of the treatment.

However, due to the unique properties of microbiome data—such as zero-inflation, compositionality, and high-dimensionality - standard mediation methods are not directly applicable. To address these challenges, we employed advanced mediation techniques tailored for high-dimensional microbiome data with multiple mediators. These approaches estimate both the total indirect effect through all microbiome features and component-wise indirect effects of individual features.

To identify microbial multi-omics features that mediate the effects of MTT, we conducted modality-specific mediation analyses across species, KOs, fecal metabolites, and plasma metabolites. Specifically, we applied a regularized multimodal mediation method to treatment- feature-symptom triplets, which accounts for the compositional and high-dimensional nature of microbiome mediators^46^ Causal interpretation of mediation effects relies on key assumptions, including the absence of unmeasured confounding after adjusting for covariates and, for tractability in high-dimensional settings, the simplifying assumption that microbial and metabolite features act as conditionally independent parallel mediators given the exposure—despite the known biological influence of microbes on metabolite levels^47^. For symptom assessments, we used the same clinical measurements as in our previous study^10,15^ (**Methods**), including the Gastrointestinal Symptoms Rating Scale (GSRS), Parent Global Impressions (PGI), Childhood Autism Rating Scale (CARS), Aberrant Behavior Checklist (ABC), Social Responsiveness Scale (SRS), and the Vineland Adaptive Behavior Scale (Vineland).

We identified several key mediators of MTT-associated symptom improvements (**Fig. 5A**; **S9**). Among these was hycholate (plasma), a primary bile acid involved in the digestion, absorption, and metabolism of dietary fats and fat-soluble nutrients (**Fig. 5B**). In our analysis, mediation analysis showed that increase in hycholate levels after 10 weeks of MTT compared to baseline contributed to reductions in CARS scores. At the KO level, we observed a decreased abundance of K01687, a key enzyme involved in the biosynthesis of branched-chain amino acids (BCAA) such as valine, leucine, and isoleucine, after 2 years of MTT compared to 10 weeks (**Fig. 5C)**. This finding aligns with prior reports of BCAA dysregulation in ASD and suggests that reduced BCAA synthesis may support symptom improvement^48,49^. Other notable mediators included cysteine-glutathione disulfide (**Fig. 5D**), *Ruminococcaceae bacterium* (**Fig. 5E**), and the long-chain fatty acid 10-nonadecenoate (**Fig. 5F**). Additionally, increased abundance of *Alistipes ihumii* was associated with greater improvement in ABC scores (**Fig. 5G**), while decreased abundance of *Alistipes finegoldii* after 10 weeks was linked to improvement in CARS scores (**Fig. S9**). Together, these results highlight the complex interplay between microbial composition and ASD symptom modulation, offering insights into the mechanisms through which MTT may exert its therapeutic effects.

**Figure 5.**
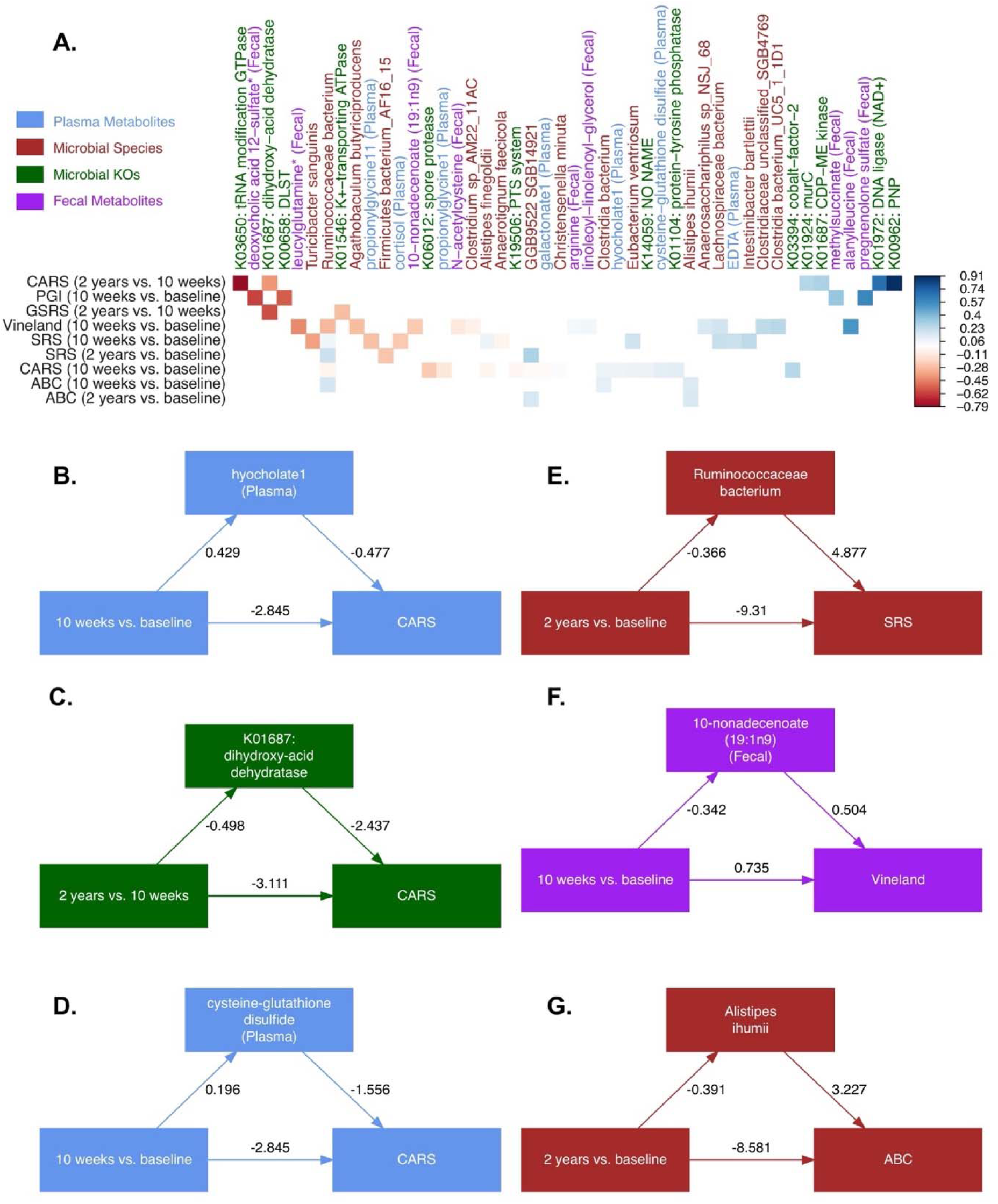
**Causal mediation analysis revealed several key mediation linkages among the gut microbiome, metabolites, and ASD symptoms**. **A**. Correlogram plot showing the multi-omics features that were associated with ASD symptoms across time points. Potentially mechanistic associations between MTT-associated microbes and metabolites and ASD symptoms included bile acids (**B**), long-chain fatty acids (**C**), cysteine-glutathione disulfide (**D**), Ruminococceae (**E**), amino acid biosynthesis (**F**), and *Alistipes ihumii* (**G**). Mediation plots in **B-G** should be interpreted as MTT causally contributed to improvements in ASD symptoms mediated through microbes and metabolites. Each mediation triplet includes: the top-left arrow (α) representing the effect of treatment on a mediator, the top-right arrow (β) representing the effect of that mediator on symptoms (adjusting for treatment), and the bottom arrow indicating the direct effect of treatment on symptoms not explained by the mediator. Arrows are labeled with standardized coefficients. The product αβ quantifies the indirect (mediated) effect. A negative mediated effect (αβ < 0) suggests the mediator contributes to symptom improvement, while a positive αβ implies the mediator may counteract treatment benefit or contribute to symptom worsening.

## Discussion

While the incidence of ASD is increasing worldwide, treatment remains largely focused on symptoms management^11,50,51^, highlighting a significant unmet need for direct characterization of multi-omics biomarkers associated with ASD. Many human studies have suggested significant differences in the gut microbiota of individuals with ASD as compared to their TD peers, characterized by reduced levels of beneficial bacteria such as *Bifidobacterium* and fiber consumers like *Prevotella*, and increased levels of *Clostridia*, *Sutterella*, and *Bacteroides*^9,14,15,27,52,53^. These microbes can influence the gut environment and gut-brain axis through the production of metabolites, which play roles in signaling, immune system modulation, and even antibiotic activity^53–56^, and can have beneficial or harmful effects. For example, butyrate is a beneficial metabolite that serves as the primary nutrient for the cells lining the intestinal tract^57,58^. In contrast, p-cresol is a harmful metabolite that acts as an antibiotic, neurotoxin, mitochondrial toxin, and nephrotoxin, and disrupts immune function and dopamine metabolism^59–63^. However, beyond a handful of well-characterized interactions, the specific microbes, the small molecules they produce, and how they interact to contribute to, sustain, mitigate, or predict ASD symptoms remain unclear.

Our study highlights treatment-associated shifts in the microbiota–gut–brain axis in ASD, uncovering key microbial players potentially influenced by MTT. Unlike prior studies focused solely on differential abundance, we identified additional microbial features that may mediate treatment effects on neurodevelopment. By integrating microbial taxa with metabolite profiles - particularly those previously underexplored - we present a more mechanistic perspective on how treatment alters host–microbiome interactions. Notably, several butyrate-producing and SCFA-associated microbes emerged as promising mediators of treatment response. Furthermore, microbial co-occurrence association networks revealed treatment-impacted interaction hubs that may modulate neurodevelopmental outcomes, offering targeted candidates for future experimental validation.

Consistent with previous studies, unsupervised multi-omics analyses revealed that children with ASD occupy distinct biological states that are quantifiable across microbiome and metabolome profiles, with these states further shifting in response to MTT. The clear separations observed in unsupervised ordination analyses, combined with high predictive performance in classifying treatment status, support the robustness of these biological distinctions. Importantly, the finding that post-treatment profiles are more readily distinguishable from baseline ASD profiles than ASD is from TD suggests that MTT induces substantial, measurable changes in the host– microbiome ecosystem. While some of these changes may represent partial restoration toward TD-like profiles, the emergence of a separable post-treatment state also implies the formation of a treatment-associated biological configuration that may underlie or contribute to observed clinical improvements. These results underscore the potential of multi-omics signatures not only for tracking treatment response but also for characterizing treatment-induced biological states that could guide precision interventions in ASD.

Integrated machine learning identified important omics features after 10 weeks of MTT compared to baseline (**Fig. 3**) including: **1)** increased beneficial and butyrate producers – *Faecalibacterium prausnitzii* and *Bifidobacterium -* which are known to support gut barrier function, reduce inflammation, provide energy to colonocytes^64–66^ and can improve brain function and prevent depression-like behavior in mice^67,68^; **2)** an increase in glucose metabolism-related genes, such as K00134 (glyceraldehyde 3-phosphate dehydrogenase; GAPDH), suggesting that short-chain fatty acid-producing microbes introduced via MTT may aid in glucose breakdown and enhance energy homeostasis^69^; **3)** an increase in plasma IMP and sarcosine, alongside a decrease in fecal sulfated bile acids such as chenodeoxycholic acid sulfate (a primary bile acid) and glycolithocholate sulfate (a secondary bile acid), suggesting improved energy metabolism, a potential shift in bile acids, and reduced excretion^35,70–72^. On the other hand, differential abundance analysis revealed a decrease in the relative abundance of the phenolic metabolite p-cresol—known to be elevated in ASD and capable of inducing autism-like behavior in mice—after 10 weeks of MTT compared to baseline^40,73^ (**Table S4**).

Furthermore, causal mediation analysis (**Fig. 5**) suggests that MTT modulates the gut ecology (microbes and metabolites) in children with ASD by reducing plasma levels of the bile acid hyocholate and cysteine-glutathione disulfide, as well as decreasing gut levels of *Ruminococcaceae bacterium* and Alistipes species. These shifts were associated with improvements in CARS, SRS, and other behavioral symptoms. Increased abundance of Alistipes has been reported in children with ASD^74,75^. Elevated levels of Alistipes have also been linked to pro-inflammatory states and impaired gut barrier function^74^, both commonly observed in ASD. Notably, Alistipes levels were significantly reduced following MTT in ASD children, coinciding with improvements in both gastrointestinal and behavioral symptoms^10,14,15^.

Additionally, elevated plasma concentrations of bile acids and cysteine-glutathione disulfide - key markers of metabolic dysregulation and oxidative stress - have been consistently reported in individuals with ASD^76–78^. These metabolites decreased significantly after MTT, aligning with clinical improvements in symptoms such as social responsiveness and overall autism severity^15^. This suggests that MTT may exert therapeutic effects by restoring microbial composition and rebalancing host metabolic pathways related to redox homeostasis and bile acid signaling, which are implicated in the gut-brain axis dysfunction in ASD.

There are several strengths of the current study. First, updated taxonomic profiling using MetaPhlAn 4 identified ASD- and treatment-associated species and pathways that were previously unrecognized. Second, while Similarity Network Fusion (SNF) hinted at a convergence of overall profiles between ASD and TD children two years after MTT, differential network analysis captured microbial and functional variation across pre- and post-treatment time points, revealing treatment-associated hub species. Third, causal mediation analysis identified several microbial and metabolomic mediators—an approach not previously applied in ASD multi-omics research.

Nevertheless, our study has limitations, including a small sample size and an open-label design. Consequently, developing realistic causal models of ASD will require addressing the inherent complexity of the disorder through larger, randomized, double-blind, placebo-controlled trials. This will necessitate multidisciplinary strategies and a concerted effort to analyze multiple novel omics endpoints in parallel. We anticipate that this study will pave the way for discovering comprehensive multi-omics signatures associated with ASD outcomes, validating prior findings while uncovering new insights. To our knowledge, no previous study has employed these methods to longitudinally investigate the ASD microbiome in the context of MTT. Ultimately, our work underscores the importance of advanced multi-omics approaches in enhancing mechanistic understanding and guiding the development of more effective treatments for ASD.

## Methods

### Metagenomics Sequencing

The details of the metagenomic sequencing have been described in the previous study^14^. Briefly, fecal DNA extracted from earlier studies was used for sequencing. The DNA samples were from three distinct timepoints: ASD at baseline (n = 18), at the end of the 10 week treatment (n = 18), and at the 2-year follow-up (n = 16). Additional samples were collected from the TD group (n = 20), major donor (n = 5), and maintenance donor (n = 2) cohorts. For shotgun metagenomics, DNA was sequenced on the Illumina NextSeq 500 platform (Illumina, CA, USA) to generate 2 × 150 bp paired end reads with a minimum depth of 10 million reads.

We obtained 28,470,588 ± 9,835,000 (mean ± standard error) reads per sample from shotgun metagenomic sequencing. The quality of raw reads was assessed using MultiQC. Adapters and low-quality reads (length < 50 bp or PHRED < 30) were removed. To avoid human genome contamination, reads were mapped against the UCSC Genome Browser’s hg38 human genome reference database using the Burrows–Wheeler aligner (bwa), and mapped reads were discarded. Reads that were unmapped to the human genome were retained for downstream analyses.

Species-level profiling was characterized using MetaPhlAn4^79^ (default settings; https://github.com/biobakery/biobakery/wiki/metaphlan4). The MetaPhlAn4 method involves mapping filtered sequencing reads to a set of species-specific genomic markers, followed by an estimation of the relative abundance of each species based on the distribution of mapped reads. MetaPhlAn4 integrates an expanded compendium of microbial genomes and metagenome- assembled genomes (MAGs) to define a broad set of species-level genome bins (SGBs), including both known species (kSGBs) and novel, uncharacterized species (uSGBs). Based on 1.01 million bacterial and archaeal MAGs and isolate genomes, the MetaPhlan4 database was mpa_vJan21_CHOCOPhlAnSGB_202103. This enhanced dataset expanded the definition to 54,596 SGBs and identified unique marker genes for 21,978 kSGBs and 4,992 uSGBs allowing for more accurate taxonomic profiling.

For functional profiling, HUMAnN3^80^ (https://github.com/biobakery/biobakery/wiki/humann3) was used to identify functional genes and pathways associated with microbiome markers. HUMAnN3 integrates with MetaPhlAn4 and its ChocoPhlAn pangenome database, and utilizes the MetaCyc, MinPath, and UniRef90 databases. Gene family abundances with at least a 90% match to UniRef90 were converted to KEGG Orthologs (KOs). Relative abundance was then calculated from the absolute abundance of KOs for each sample.

### Fecal and Plasma Metabolomics

An untargeted liquid chromatography–mass spectrometry (LC–MS) metabolomic profiling approach was previously performed by Metabolon Inc. to measure fecal and plasma metabolites using ultra-high-performance liquid chromatography tandem mass spectroscopy (UHPLC- MS/MS) (https://www.metabolon.com) (accessed on 10 January 2022). Sample preparation, metabolite measurements, and results are described in our prior work^19^.

### GI and ASD Symptoms Assessment

All GI and ASD symptoms measurements are described in detail elsewhere^10,15^. In brief, for GI symptoms, a revised version of the Gastrointestinal Symptoms Rating Scale (GSRS) with the five domains of Abdominal Pain, Reflux, Indigestion, Diarrhea, and Constipation was used. Daily stool records using the Bristol Stool Form scale were also recorded. For ASD symptoms, Parent Global Impressions–III (PGI-III), Childhood Autism Rating Scale (CARS), Aberrant Behavior Checklist (ABC), Social Responsiveness Scale (SRS), and Vineland Adaptive Behavior Scale II (VABS-II) measurements were taken for all ASD participants.

### Filtering, Transformation, and Normalization

To reduce the effect of sparsity in microbiome data in all subsequent analyses, features with no variance or with more than 90% zeros were removed. In addition, a variance filtering step was applied to remove features with very low variance. While we operated on the normalized relative abundance scale for microbial species and KOs, for the quality control and normalization of metabolomics data, we implemented a median normalization approach in combination with winsorization using the R package *metabolomicsR (*https://github.com/XikunHan/metabolomicsR). We did not transform the data, as we used a Tweedie compound Poisson model for all biotypes, which employs a log-link function to model the untransformed measurements, allowing the exploitation of the mean-variance relationship in the data without an ad hoc transformation.

### Ordination Analysis

We performed centered log-ratio (CLR) transformation for microbiome data and log transformation for metabolomics data, enabling appropriate compositional handling and variance stabilization, respectively, while using Euclidean distance-based principal coordinates analysis (PCoA) to assess separability across ASD time points and between ASD and TD groups in multi-omics profiles. Prior to integrated analyses, data from each modality were scaled to unit variance to ensure that no single modality dominated the integrated analysis. PCoA was performed using the *cmdscale* function in the R package *vegan*. The variance explained by the first two principal coordinates was calculated, and group separations were assessed using permutational multivariate analysis of variance (PERMANOVA) with 999 permutations (*adonis2*, *vegan*).

### Differential Abundance Analysis

Differential abundance (DA) analysis of all microbial measurement types in ASD was tested using a per-feature mixed effects model, in which, subjects were included as random effects to account for the correlations in the repeated measures, and the relative abundance of each feature was modeled as a function of pre-post treatment status (a categorical variable with the preceding time point as the reference group). For the DA analysis of ASD vs. TD children, the relative abundance of each feature was modeled as a function of ASD disease status (a categorical variable with TD as the reference group). Fitting was performed with the R package *Tweedieverse (*https://github.com/himelmallick/Tweedieverse), which implements a Tweedie compound Poisson generalized linear mixed-effects model (GLMM) with a degenerate distribution at the origin and a continuous distribution on the positive real line. The significance of the association was assessed using Wald’s test. We corrected for multiple hypothesis testing using a Benjamini-Hochberg discovery rate (FDR) approach with a target FDR of 0.05.

### Integrated Machine Learning

We used the R package *IntegratedLearner* (https://github.com/himelmallick/IntegratedLearner) for multimodal predictive model building. Briefly, *IntegratedLearner* provides an integrated machine learning framework to consolidate predictions by borrowing information across several longitudinal and cross-sectional omics data layers using a variety of integration paradigms and base models within a unified estimation framework. We considered three integrated classifiers for predicting the pre/post status: 1) 10 weeks from baseline vs. baseline, 2) 2 years after MTT vs. 10 weeks from baseline, and 3) 2 years after MTT vs. baseline. In each case, subject labels were randomly balanced prior to training, and default settings were used. Feature importance scores from the integrated models were retained for downstream analysis. The area under the receiver operating characteristic (ROC) curve (AUC) was used as evaluation metric for assessing prediction performance.

### Differential Network Analysis

We used the R package *NetCoMi*^81^ (https://github.com/stefpeschel/NetCoMi) for differential network analysis. NetCoMi combines Fisher’s z-test, a non-parametric resampling procedure, and the discordant method in a single workflow to build differential networks, focusing on features with significant differences in connectivity between groups. Network reconstruction was performed on CLR-normalized abundances using default parameters, with an edge detection threshold set to 0.5. We selected the top 200 most variable features based on their variance between groups. Significant edges were identified using Student’s t-test, and community structures were analyzed through greedy optimization of modularity. Hub nodes were detected with a threshold of 0.8, and the network was quantitatively assessed using a permutation approach (100,000 bootstraps) with an adaptive Benjamini–Hochberg correction for multiple testing. For network visualization, we used *layoutGroup = "union",* where a union of the two layouts is used in both groups to ensure that nodes are placed as optimal as possible equally for both networks. To achieve sparsity, we only plotted the significant associations with FDR < 0.01.

### Causal Mediation Analysis

We used the R package HIMA2^46^ (https://github.com/joyfulstones/HIMA2) for causal mediation analysis on the triplet: pre-post treatment, microbial features, and symptoms. High-dimensional mediation analysis (HIMA) is a penalized regression approach in which the outcome model is fitted with a minimax concave penalty, performing feature selection on the potential mediators. The mediator models are then fitted among the remaining mediators using ordinary linear regression. HIMA2 reduces the dimension of mediators to a manageable level based on the sure independence screening (SIS) method and employs a p-value correction procedure that maintains the FDR for detecting active mediators (FDR < 0.25).

## Funding

This work was supported by the Arizona Board of Regents, Finch Therapeutics, and the ASU Biodesign Center for Health Through Microbiomes Start-up.

### Institutional Review Board Statement

The study was conducted according to the guidelines of the Declaration of Helsinki, and approved by the Institutional Review Board (Arizona State University (ASU IRB Protocol #: 00001053 and # 00004890), as described in Kang et al., 2019^10^. The protocol for the original treatment study was approved by the FDA (Investigational New Drug number: 15886).

### Informed Consent Statement

Informed consent was obtained from all subjects involved in the study. The participant’s name and identifiers were removed and are not used in any sections of the manuscript, including the Supplementary Materials. The trial was registered at the ClinicalTrials.gov (NCT02504554) on 30 March 2015.

### Data Availability Statement

The metagenomic sequencing data presented in this study are openly available in the NCBI SRA repository under BioProject ID and can be accessed at https://www.ncbi.nlm.nih.gov/bioproject/?term=PRJNA782533. The study metadata and clinical data are available in Kang et al.^10^ Raw metabolomic data are available from Nirmalkar et al. (2024)^19^. A curated, analysis-ready dataset is available at https://github.com/himelmallick/ASDMultiomics.

### Code Availability Statement

The software packages used in this work are free and open source. Analysis scripts using these packages to generate figures and results from this manuscript (and associated usage notes) are available at https://github.com/himelmallick/ASDMultiomics.

## Acknowledgments

We gratefully thank all children with ASD and their families for participating in the study. We would like to thank Thomas Borody, Alexander Khoruts, Michael J. Sadowsky, and Alessio Fasano for their help with the treatment portion of the study and for their help with the original Investigational New Drug (IND) application and study. We also would like to acknowledge Juan Maldonado and team from the Genomics Core–Biodesign Institute ASU for metagenomic sequencing. Jason Yalim and team from the Research Computing Core Facilities (https://rcstatus.asu.edu/) at Arizona State University for his assistance and use of HPC cluster (Agave) to run the sequencing data analyses.

## Author Contributions

Conceptualization: HM, KN, JBA, RK-B; Data Curation: HM, KN, JBA; Statistical Analysis: HM, KN; Funding Acquisition: JBA, RK-B; Investigation: HM, KN, JBA, RK-B; Project Administration: JBA, RK-B; Software: HM, KN; Supervision: JBA, RK-B; Visualization: HM, KN; Writing (original draft): HM, KN, JBA, RK-B; Writing (review and editing): HM, KN, JBA, RK-B. All authors have read and agreed to the published version of the manuscript.

## Supplementary Materials

**Supplementary Figure S1**. **Integrated machine learning for predicting pre- and post- treatment ASD status from gut microbiome multi-omics features using microbial prevalences.** In this model, species and KO prevalences are used instead of abundances, while the metabolomics data remain unchanged. Random forest classifiers were trained on plasma metabolites, stool metabolites, microbial species, microbial functional profiles (KO), and their combinations to distinguish ASD children before and after MTT at both 10 weeks and 2 years. Classifiers were also trained to distinguish ASD from TD children at both baseline and post-baseline. Model training was performed using five-fold cross-validation within the IntegratedLearner framework^16^.

**Supplementary Figure S2**. **Feature selection results from the integrated machine learning model using microbial abundances**. In this model, species and KO prevalences are used instead of abundances, while the metabolomics data remain unchanged. The bar plots display feature importance scores for the top 20 features from each layer of the integrated model. Positive and negative associations are indicated by “+” and “–” signs, respectively.

**Supplementary Figure S3**. **Feature selection results from the integrated machine learning model using microbial prevalences**. In this model, species and KO prevalences are used instead of abundances, while the metabolomics data remain unchanged. The bar plots display feature importance scores for the top 20 features from each layer of the integrated model. Positive and negative associations are indicated by “+” and “–” signs, respectively.

**Supplementary Figure S4. Statistically significant differentially abundant microbial species from univariate analysis across comparisons.** ASD (Baseline) vs. TD (Baseline), ASD (10 weeks after MTT) vs. TD (Baseline), ASD (2 years after MTT) vs. TD (Baseline), ASD (10 weeks after MTT) vs. ASD (Baseline), ASD (2 years after MTT) vs. ASD (Baseline), and

ASD (2 years after MTT) vs. ASD (10 weeks after MTT). In addition to log fold changes (LFCs), the per-group prevalence of each feature is also provided. Only top 30 statistically significant results at FDR 0.25 are included. Full results are provided in **Table S1**.

**Supplementary Figure S5**. **Statistically significant differentially abundant microbial KOs from univariate analysis across comparisons.** ASD (10 weeks after MTT) vs. TD (Baseline), ASD (2 years after MTT) vs. TD (Baseline), ASD (10 weeks after MTT) vs. ASD (Baseline), ASD (2 years after MTT) vs. ASD (Baseline), and ASD (2 years after MTT) vs. ASD (10 weeks after MTT). In addition to log fold changes (LFCs), the per-group prevalence of each feature is also provided. Only top 30 statistically significant results at FDR 0.25 are included. Full results are provided in **Table S2**.

**Supplementary Figure S6**. **Statistically significant differentially abundant plasma metabolites from univariate analysis across comparisons**. Only top 30 statistically significant results at FDR 0.25 are included. LFCs are provided for ASD (Baseline) vs. TD (Baseline), ASD (10 weeks after MTT) vs. TD (Baseline), ASD (10 weeks after MTT) vs. ASD (Baseline). Full results are provided in **Table S3**.

**Supplementary Figure S7**. **Statistically significant differentially abundant fecal metabolites from univariate analysis across comparisons**. Only top 30 statistically significant results at FDR 0.25 are included. LFCs are provided for ASD (Baseline) vs. TD (Baseline), ASD (10 weeks after MTT) vs. TD (Baseline), ASD (10 weeks after MTT) vs. ASD (Baseline). Full results are provided in **Table S4**.

**Supplementary Figure S8**. **Differential network analysis of microbiome species for the 2 years after MTT vs. 10 weeks after MTT comparison in ASD children.** The thickness of the edges represents the strength of association, with a thick line indicating a very strong correlation and a thin line indicating a weak correlation. Full results are provided in **Table S5**.

**Supplementary Figure S9. Directed Acyclic Graphs (DAGs) of statistically significant multimodal mediation effects at FDR 0.25.** The numbers in the DAGs denote the strength of the corresponding directed associations.

**Supplementary Tables S1-S4. Univariate associations between multi-omics features and covariates.** The tables list statistically significant associations (FDR < 0.25) for comparisons between ASD vs. TD or ASD pre- vs. ASD post-treatment, across multiple data modalities (**S1: Species**, **S2: KOs**, **S3: Plasma Metabolites**, **S4: Fecal Metabolites**) using a multivariable mixed effects model, as described in **Methods**. Features are sorted by minimum FDR-adjusted p-values. For each feature, coefficient estimates, prevalence, and the two-tailed p-values and q- values are reported.

**Supplementary Table S5. Differential network associations microbial species and pre- post time points.** This table presents the statistically significant differentially connected associations (FDR < 0.25) for various comparisons of ASD pre- and post-treatment time points: 1) ASD (10 weeks after MTT) vs. ASD (Baseline), 2) ASD (2 years after MTT) vs. ASD (Baseline), and 3) ASD (2 years after MTT) vs. ASD (10 weeks after MTT). For each feature, the table provides pairwise correlation coefficient estimates, two-tailed p-values for differential network analysis, and corresponding FDR-adjusted p-values. Features are sorted by their minimum FDR-adjusted p-values.

