## Supplementary figures and images for "Multi-omics Integration of Microbiota Transplant Therapy in Children with Autism Spectrum Disorders"

### Figure S1

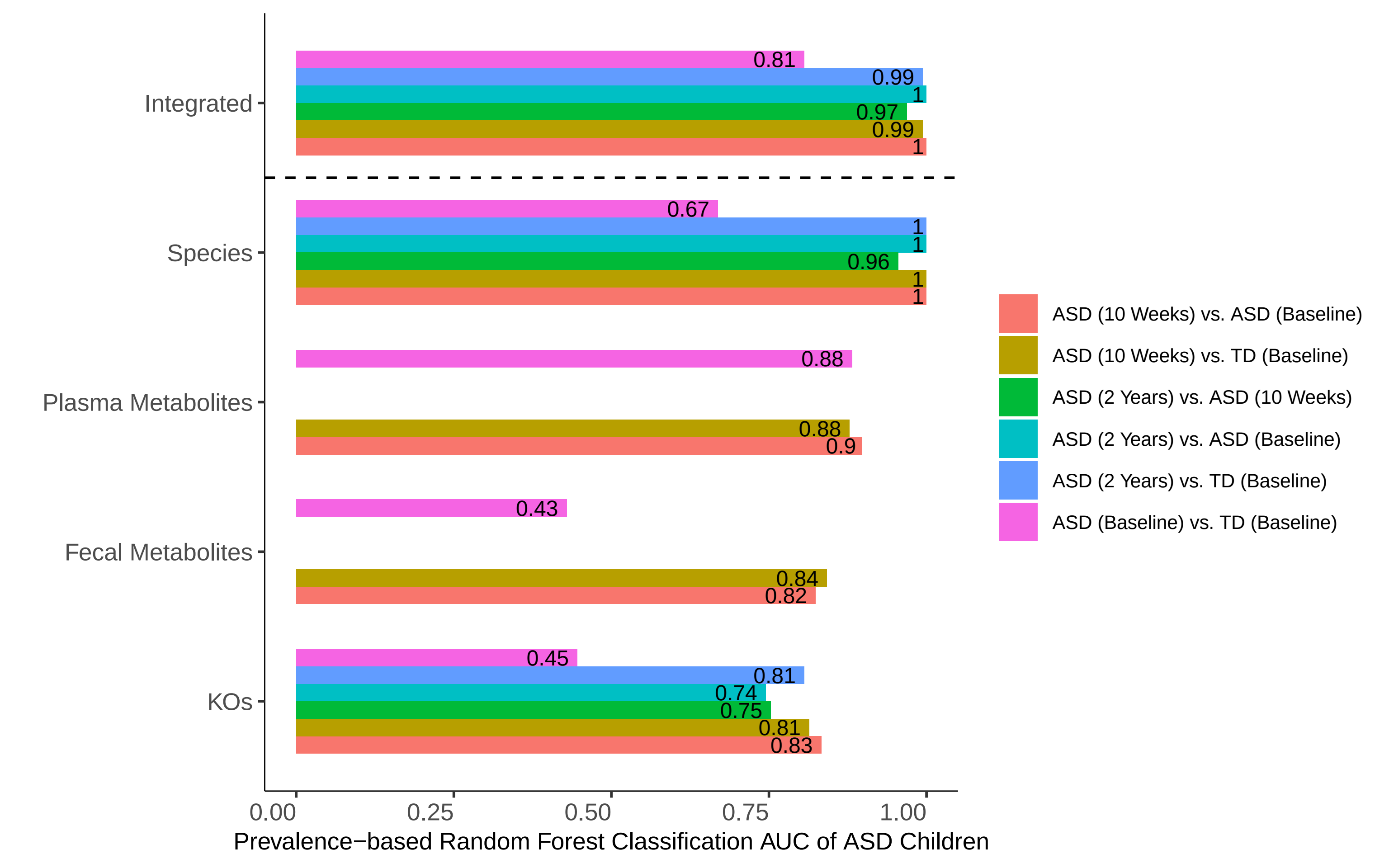

### Figure S2

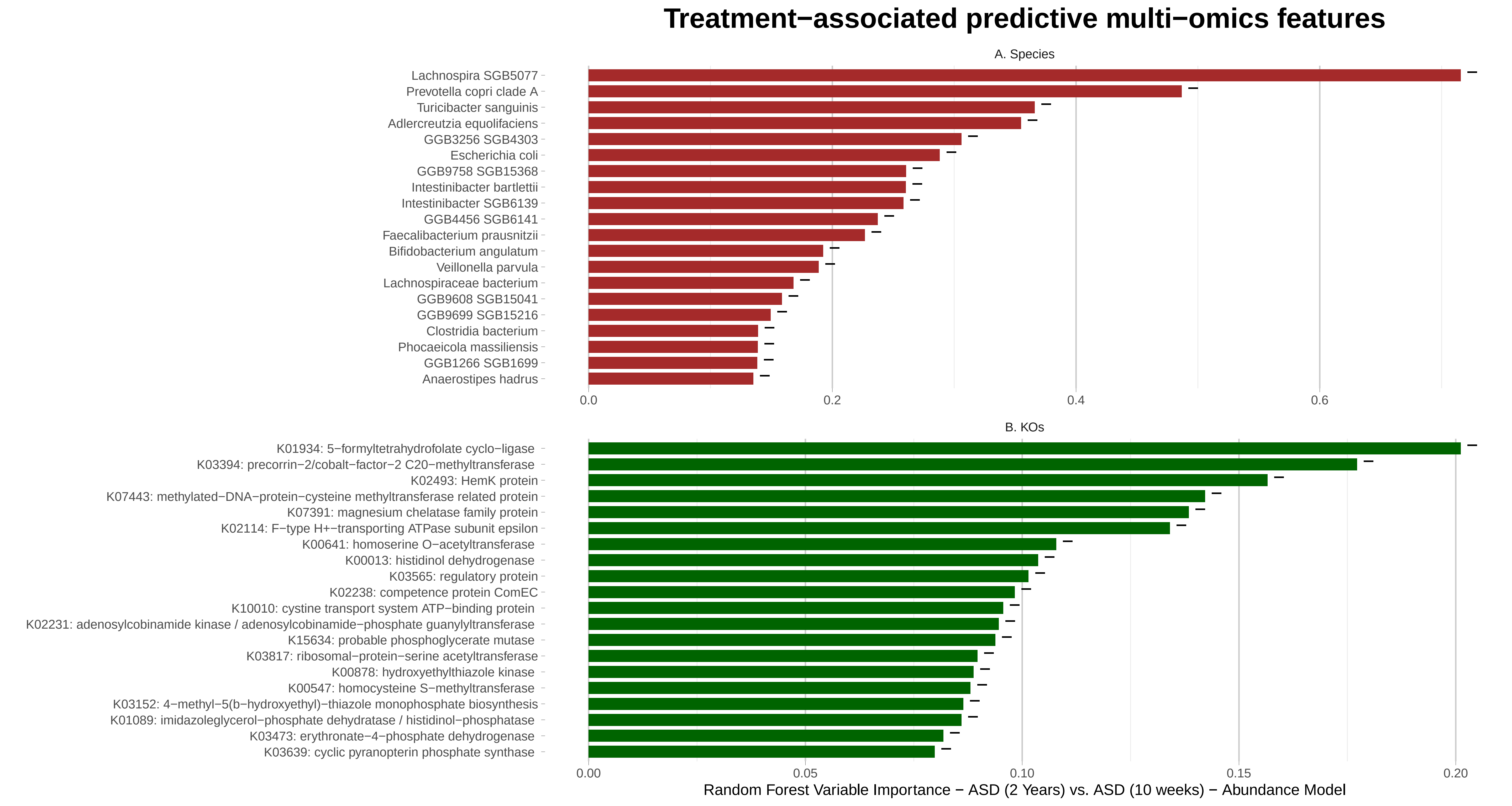

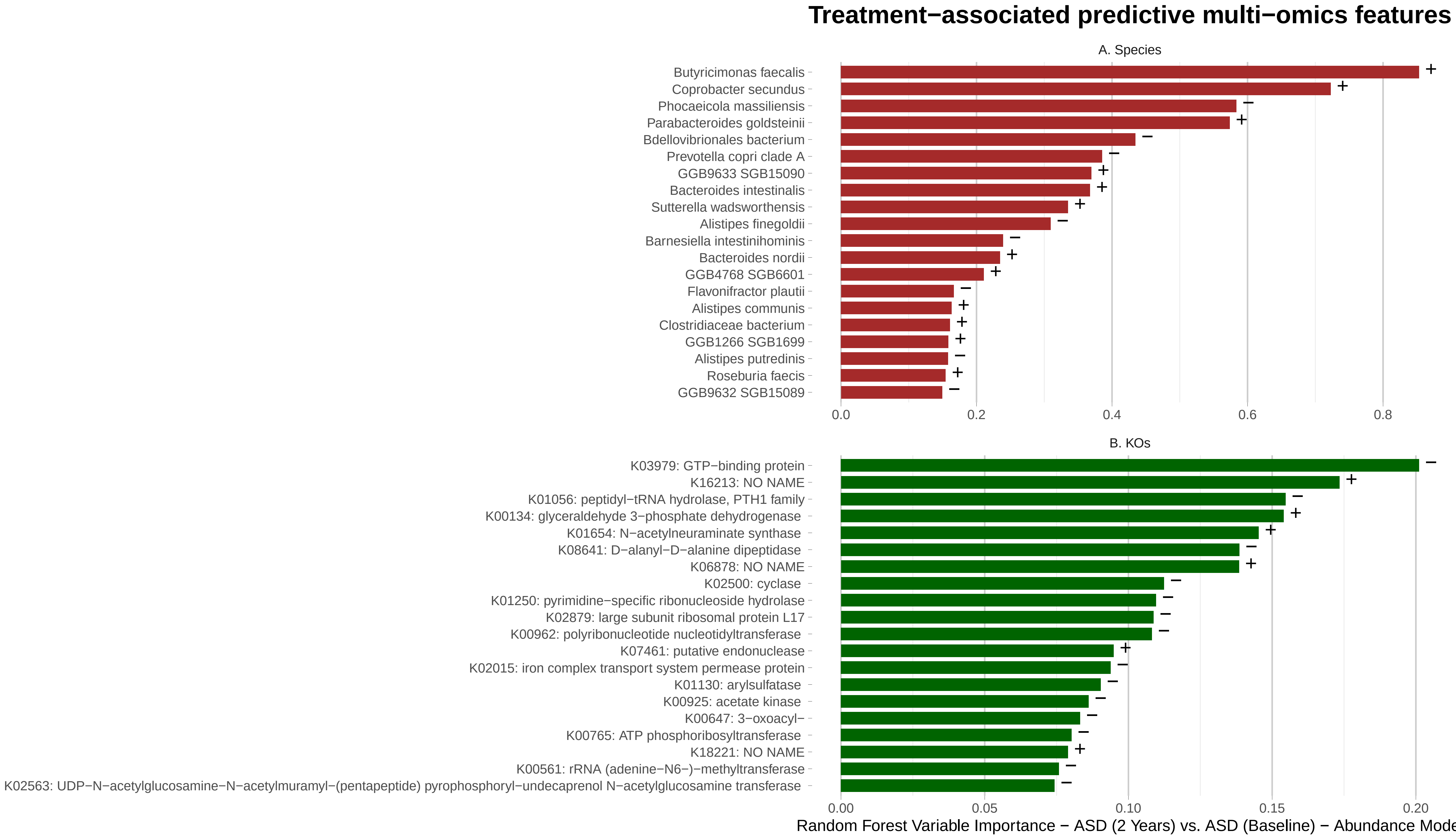

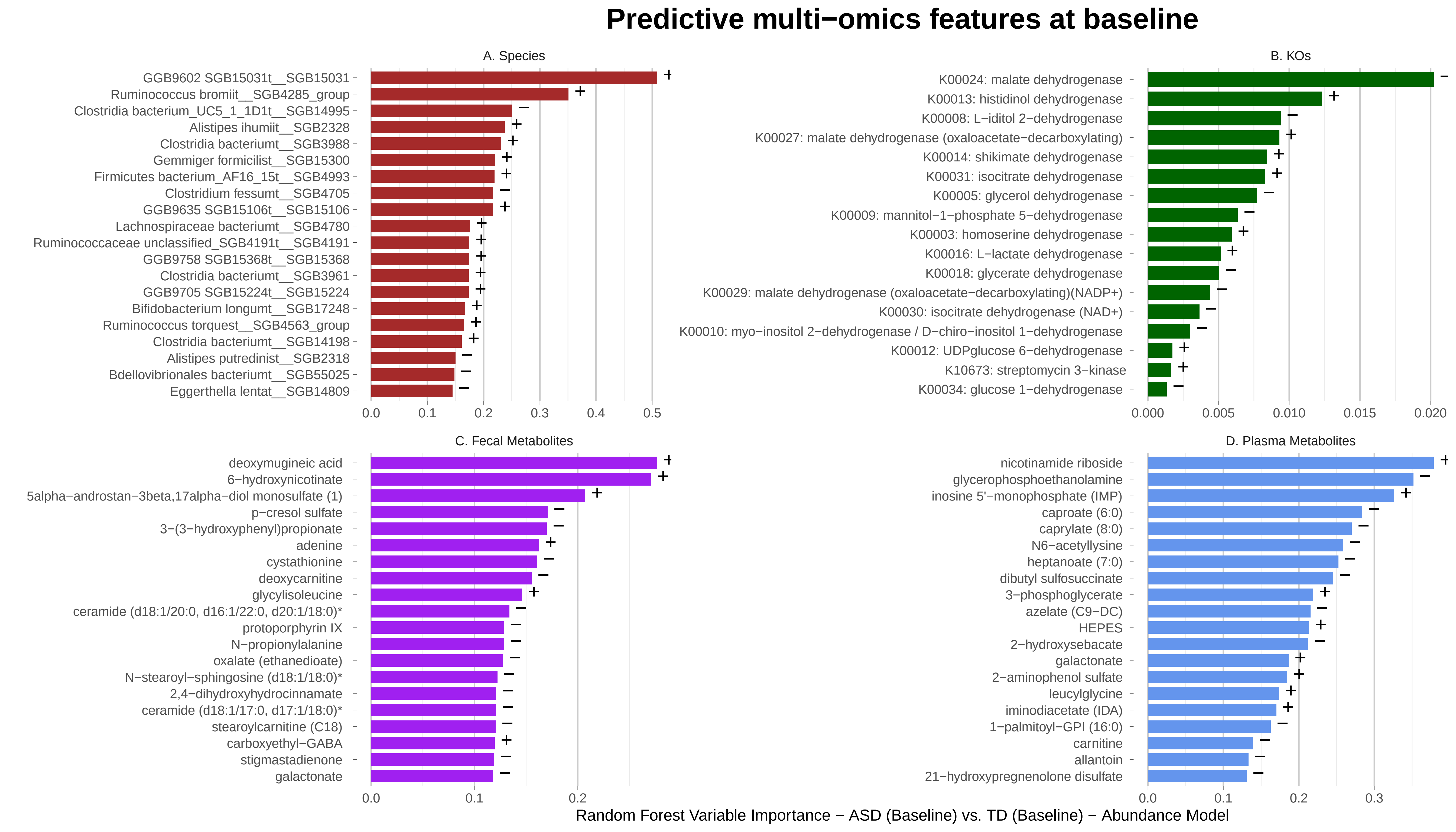

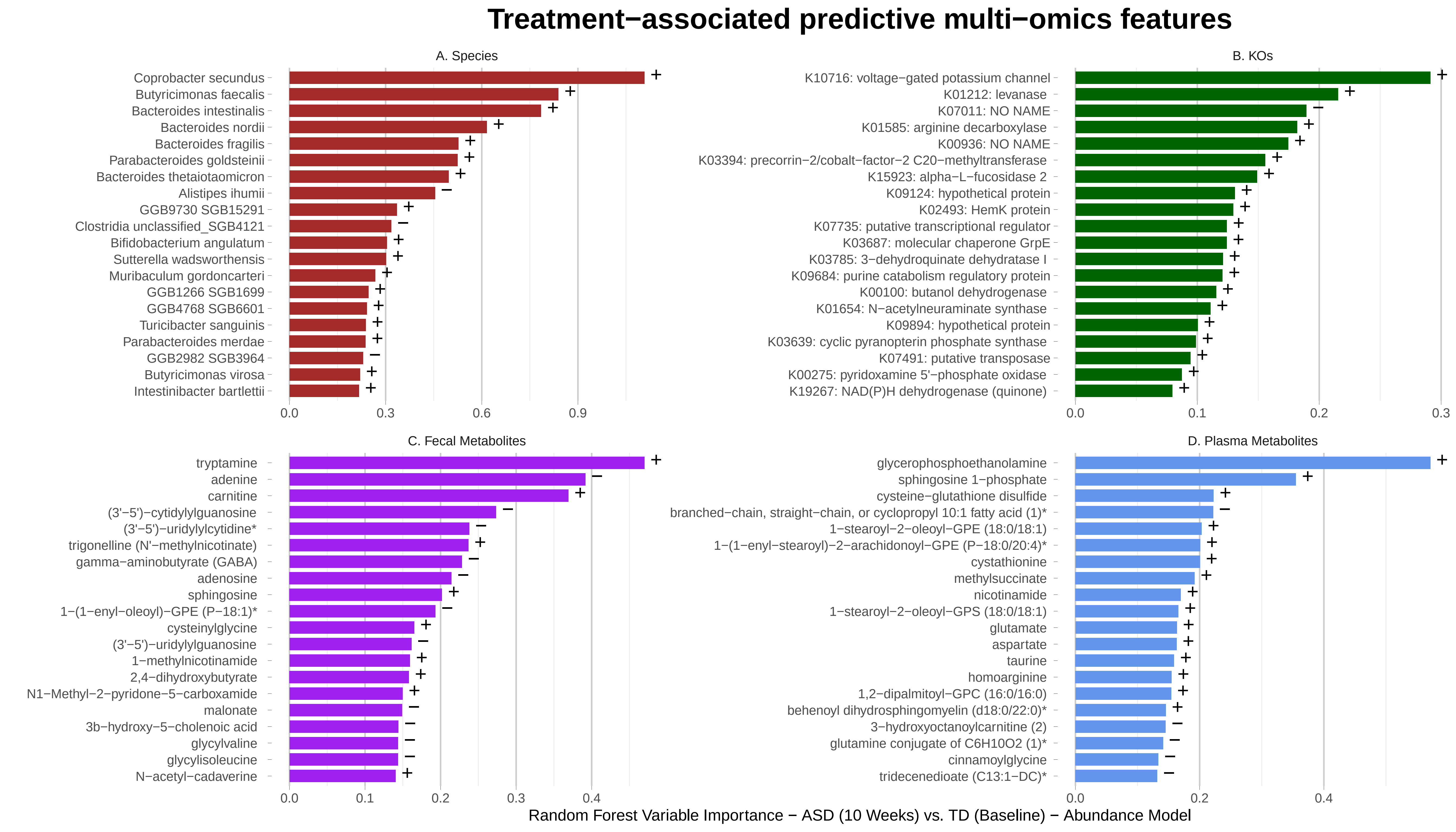

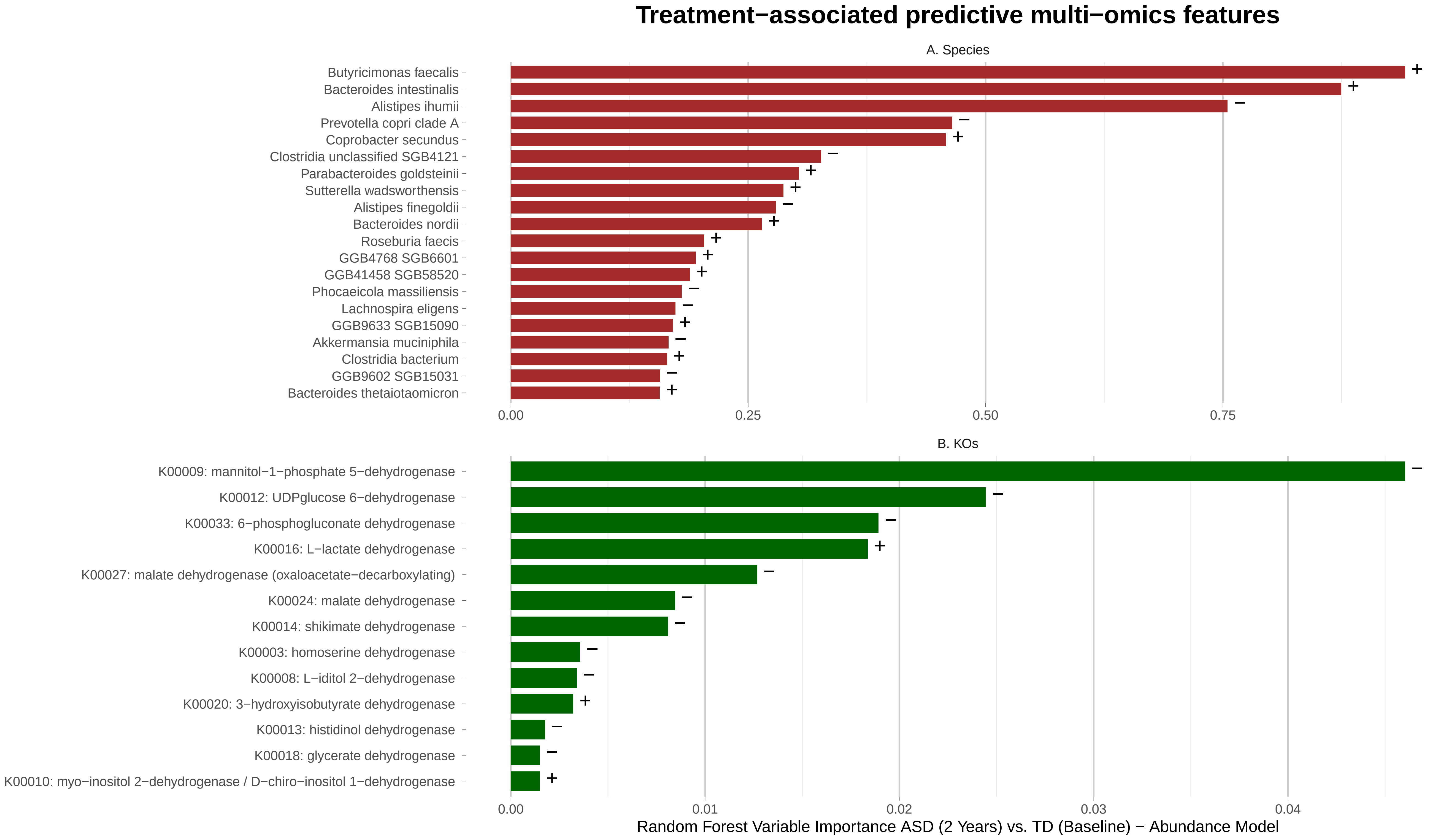

### Figure S3

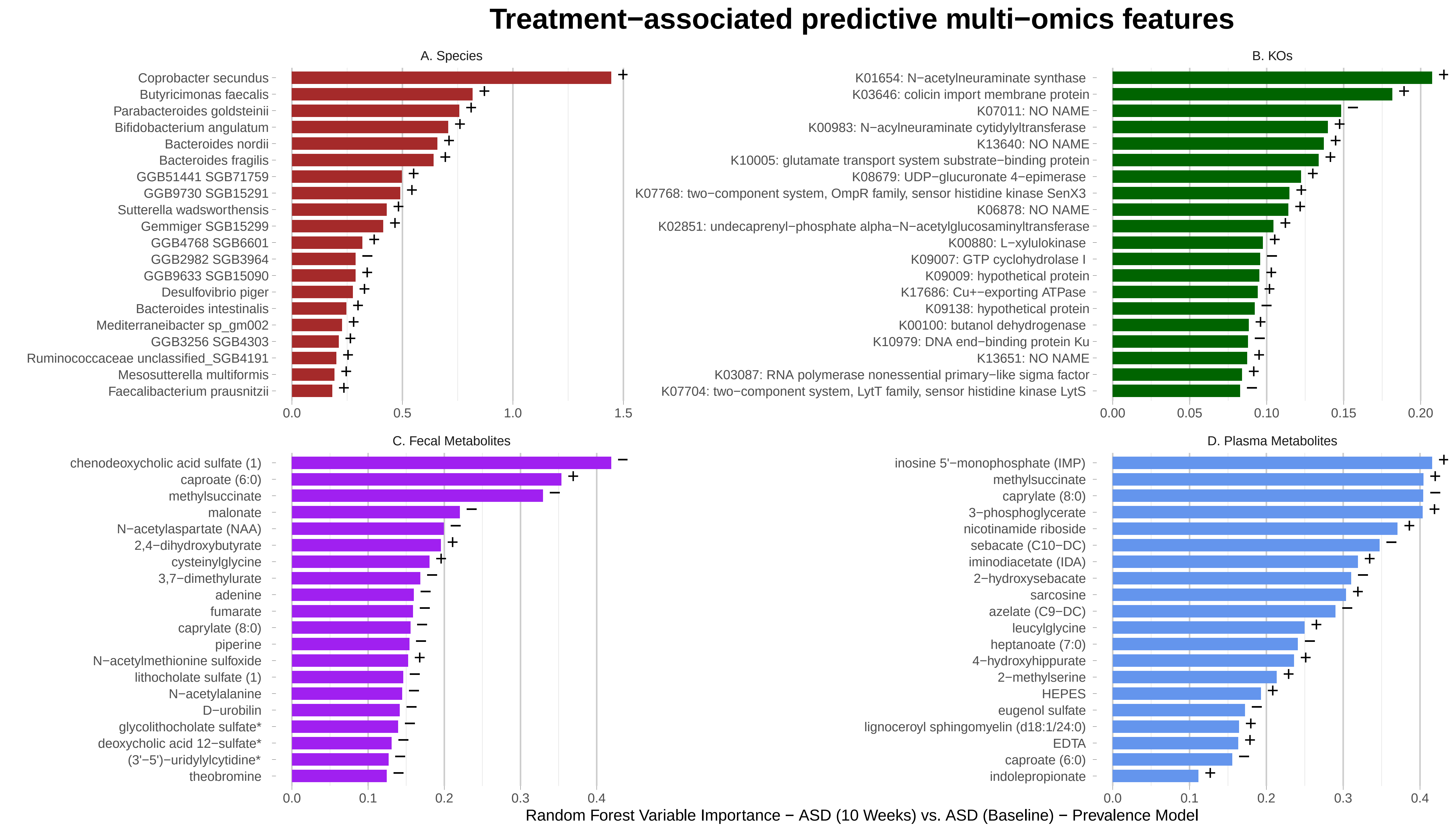

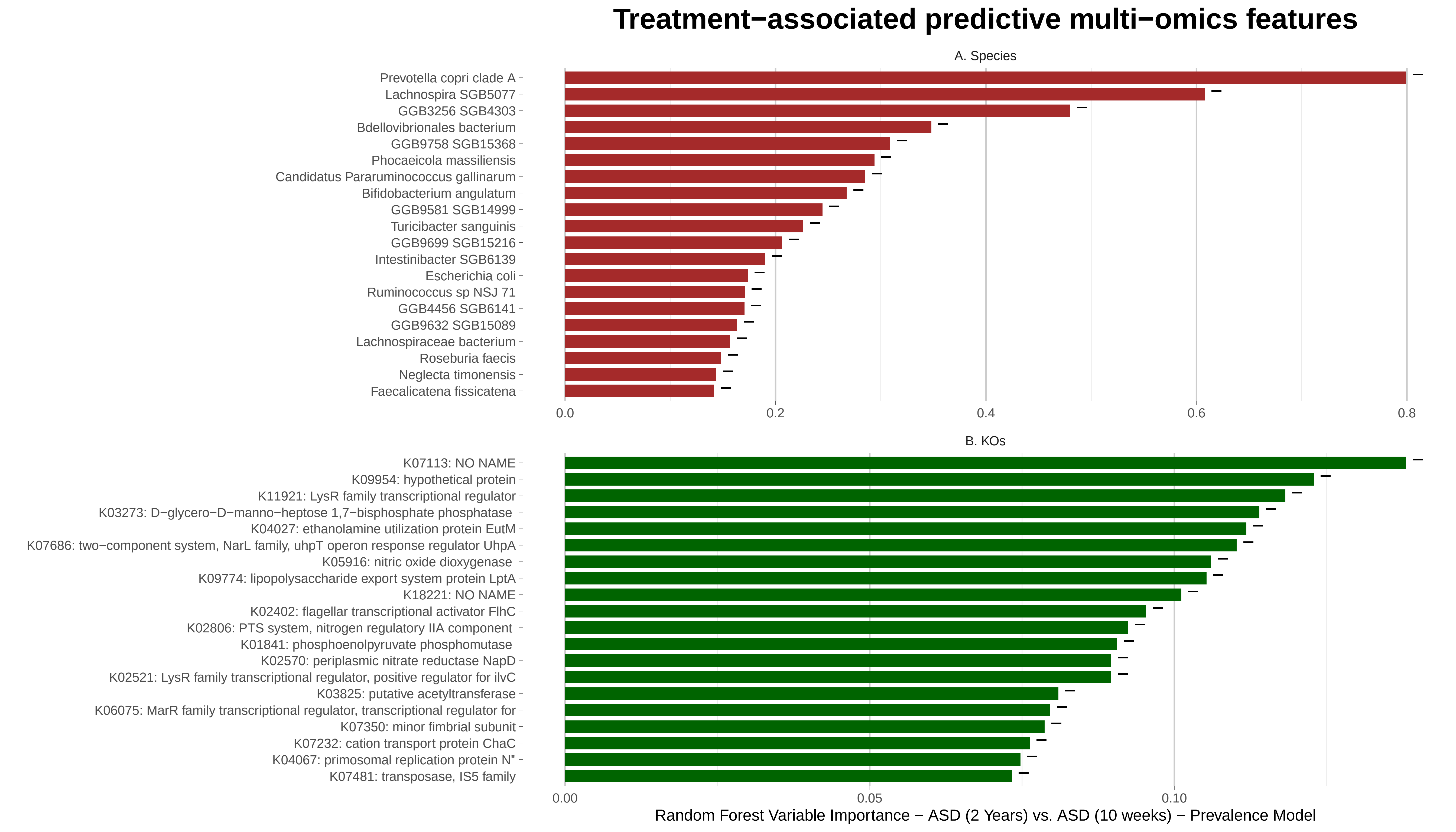

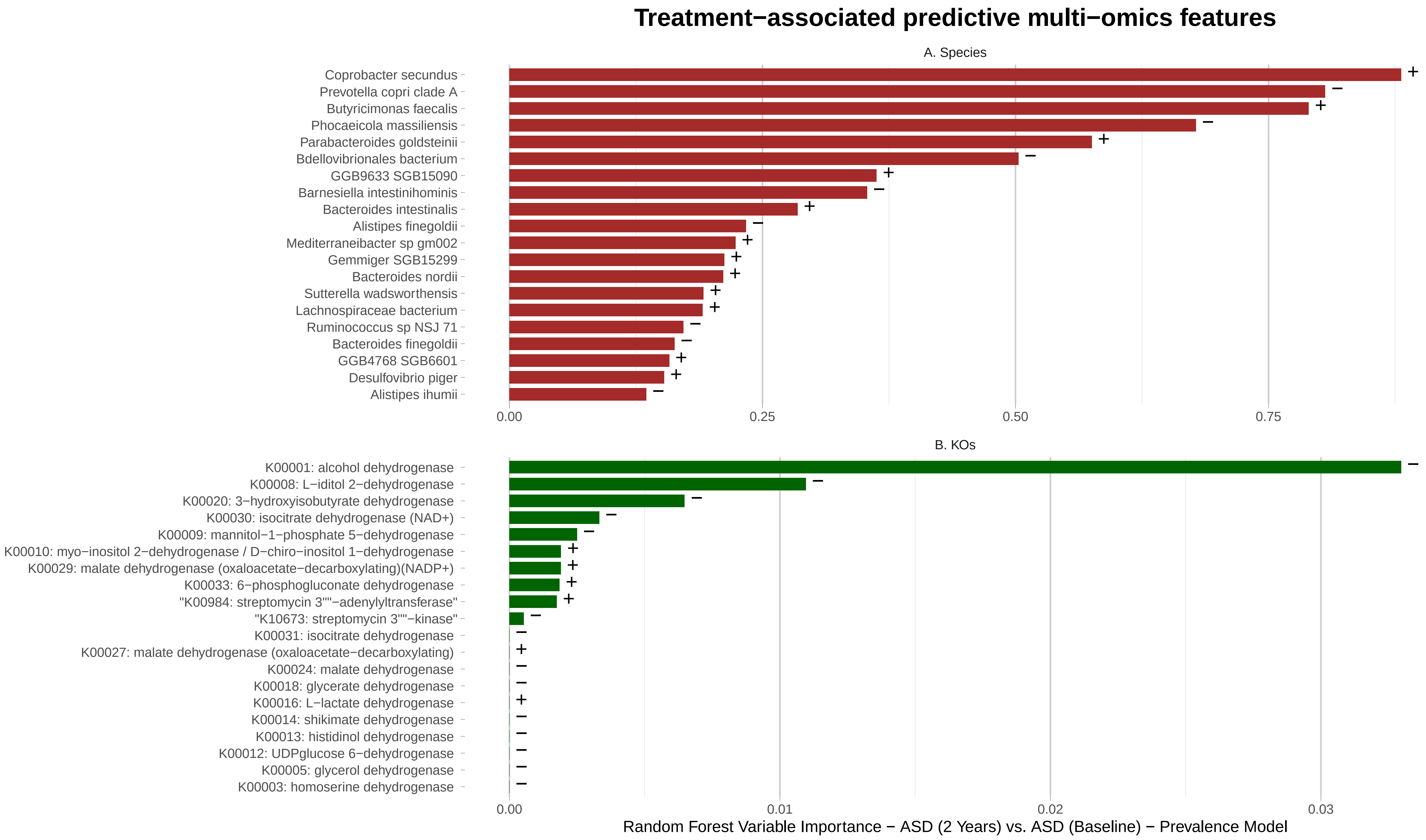

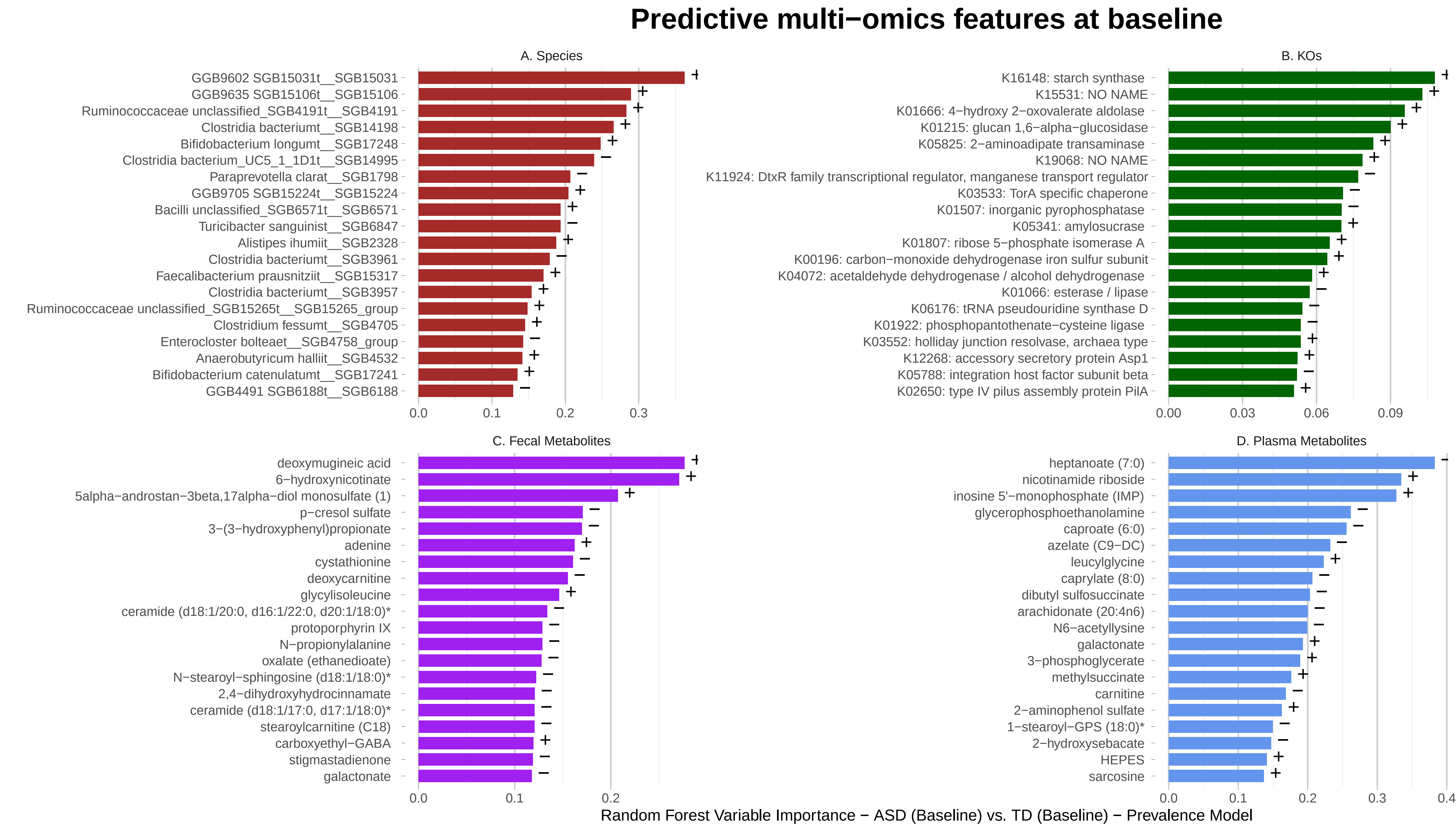

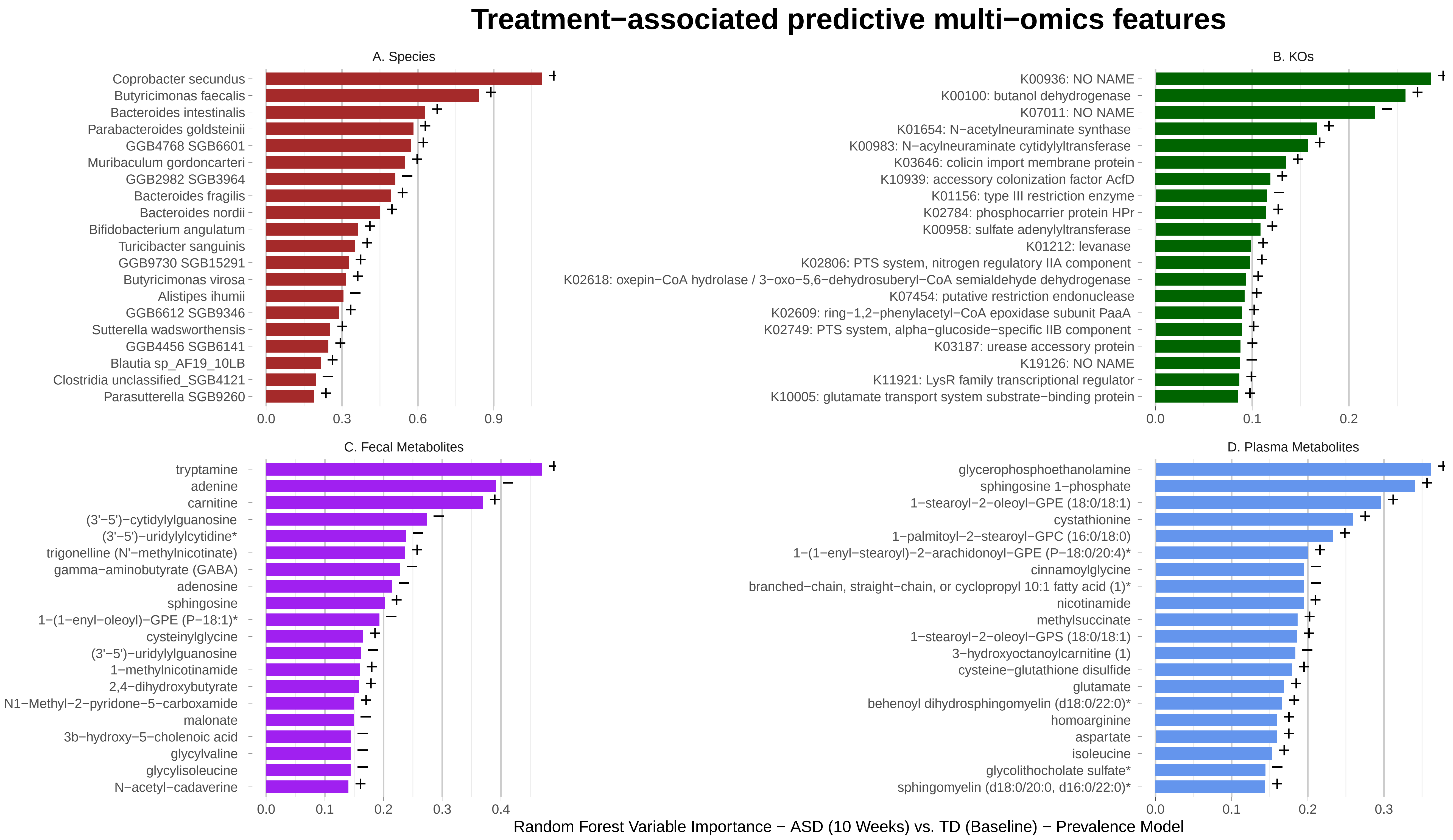

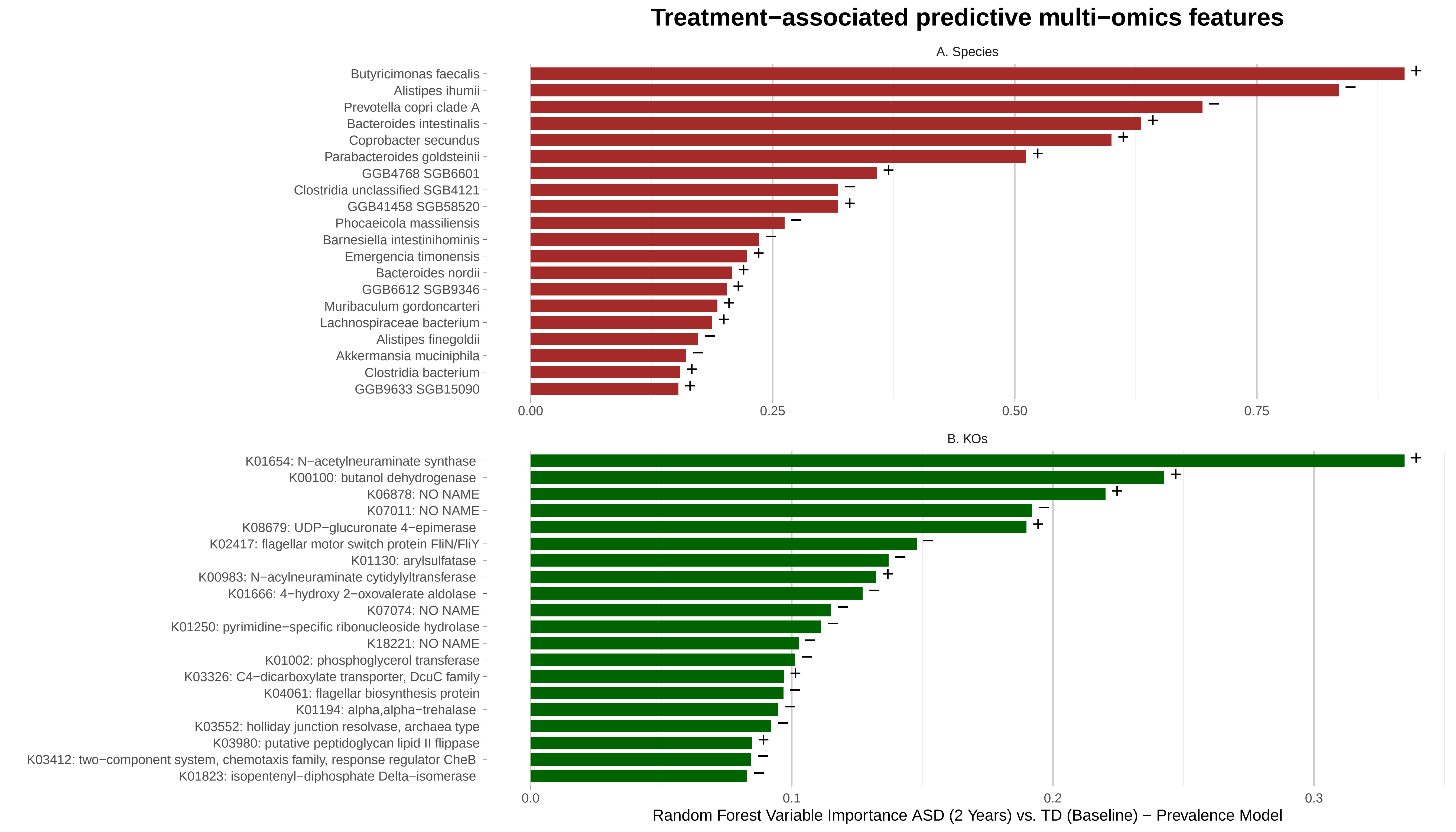

### Figure S4

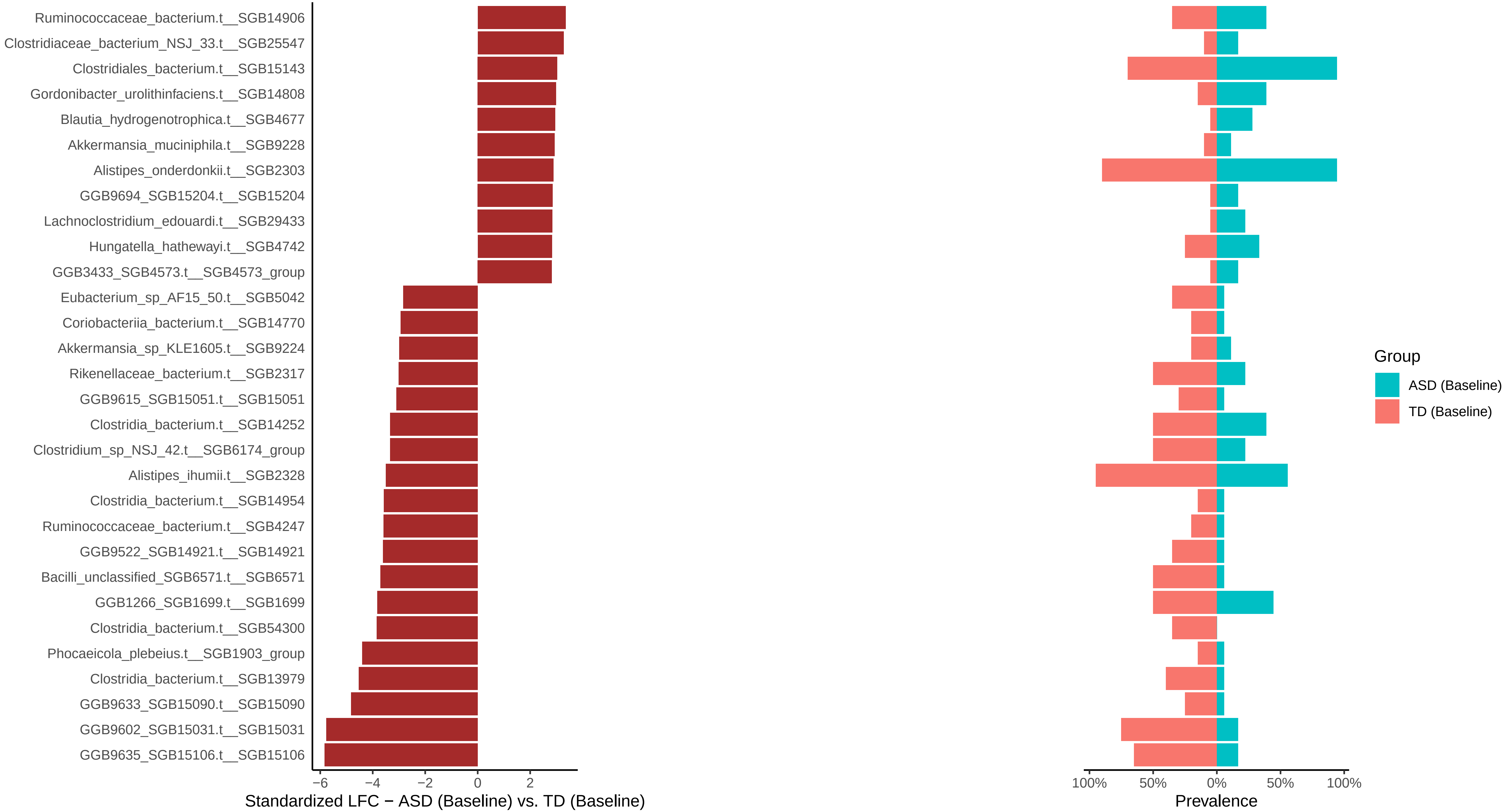

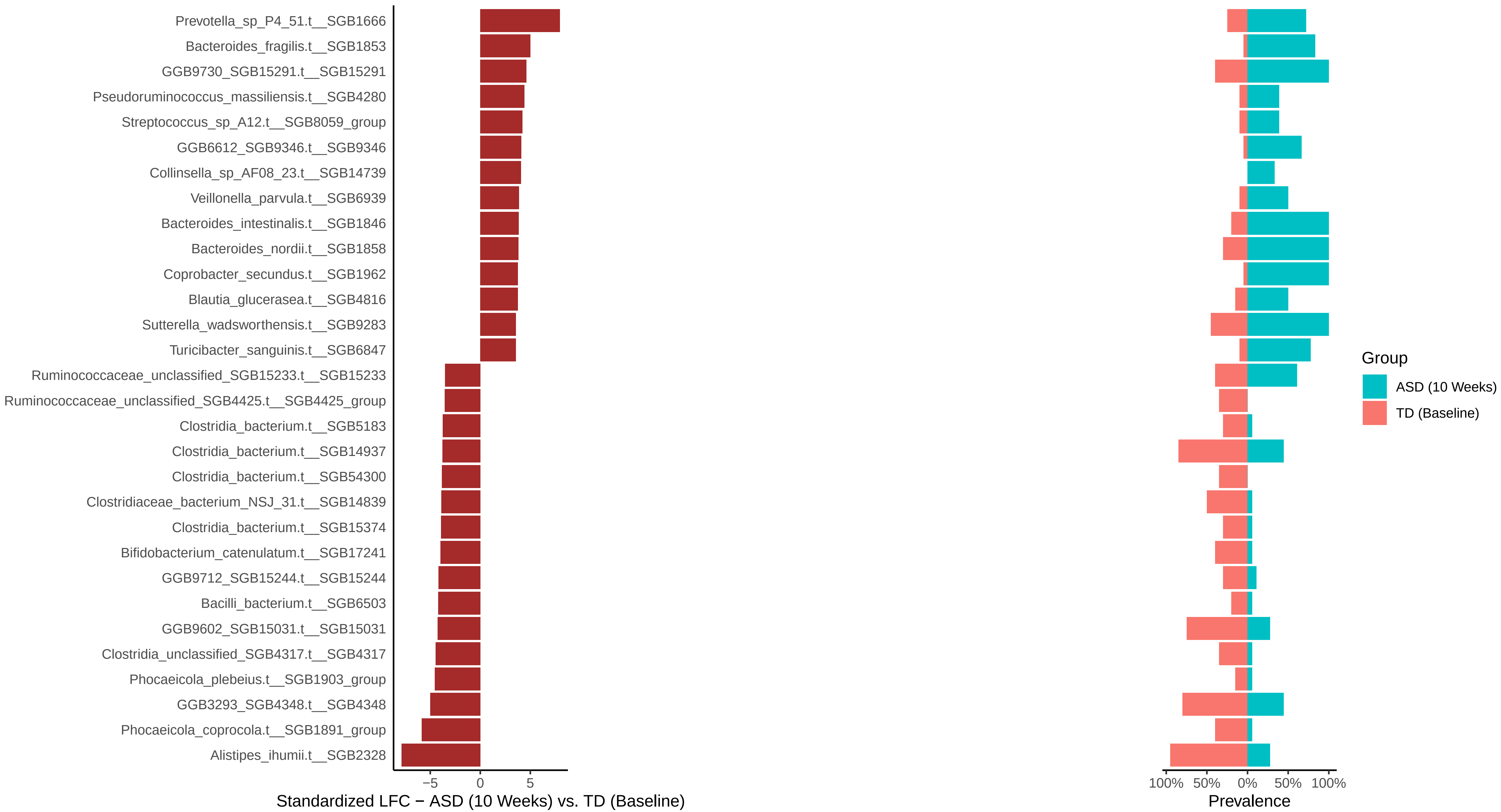

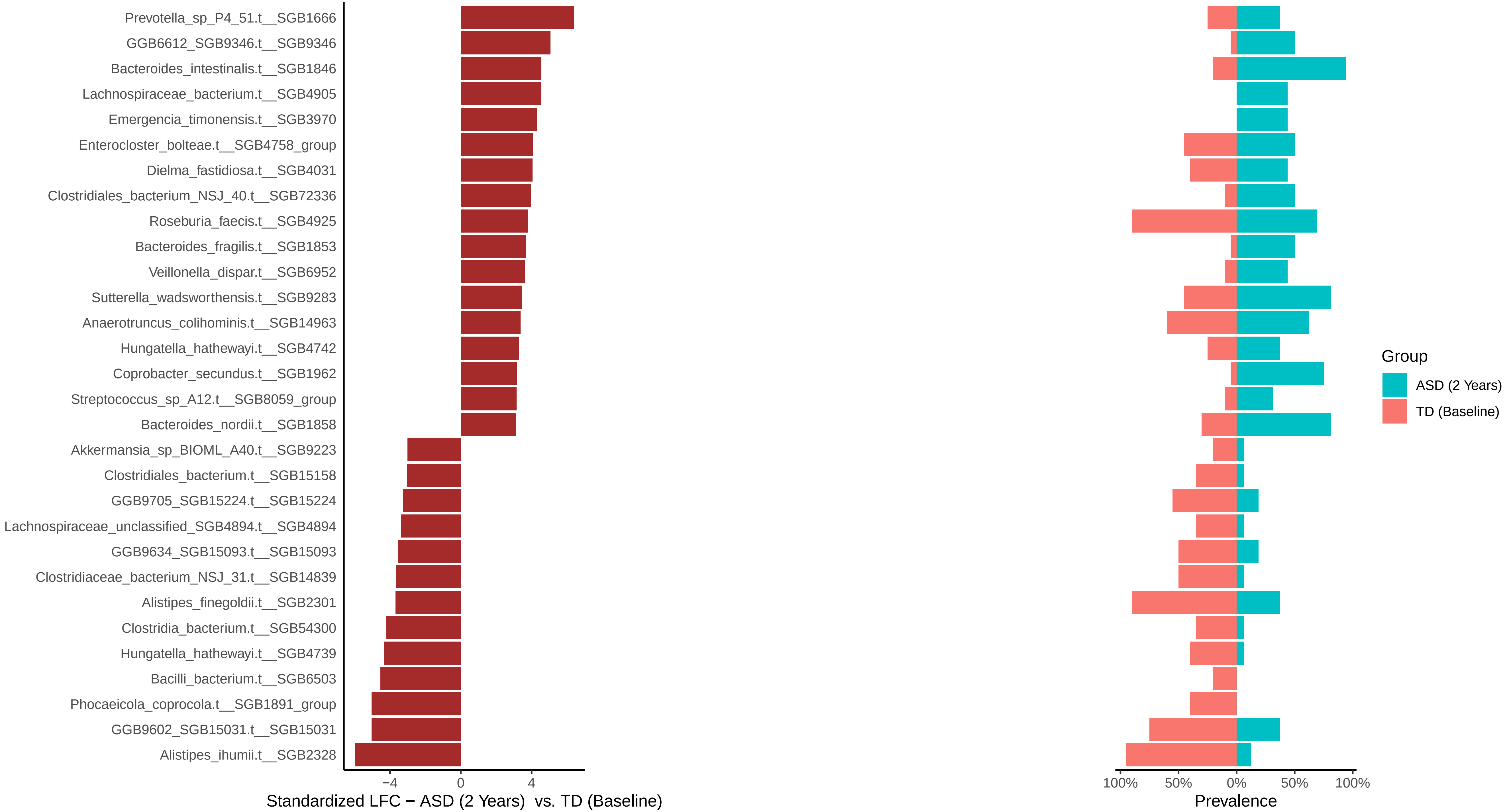

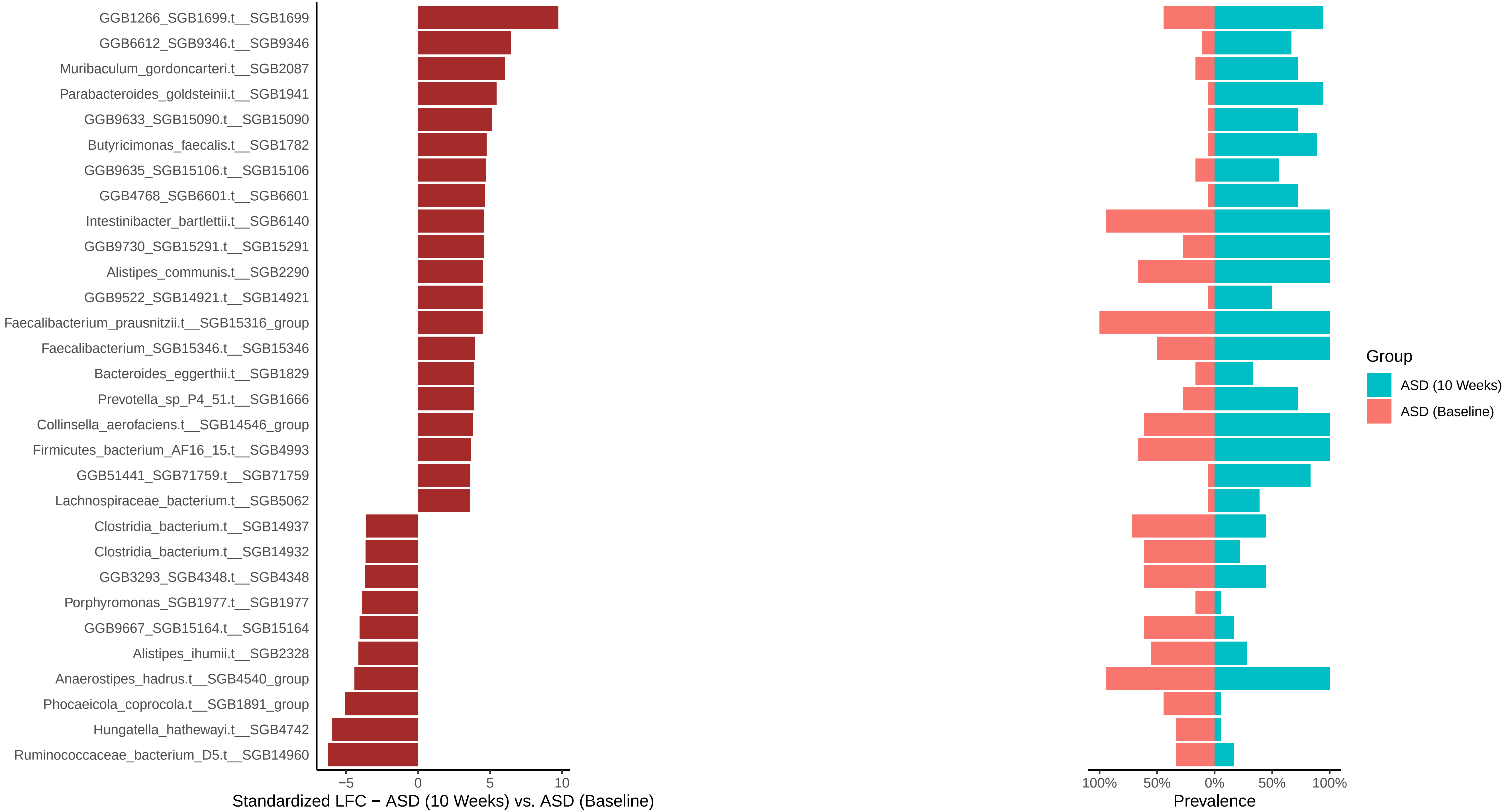

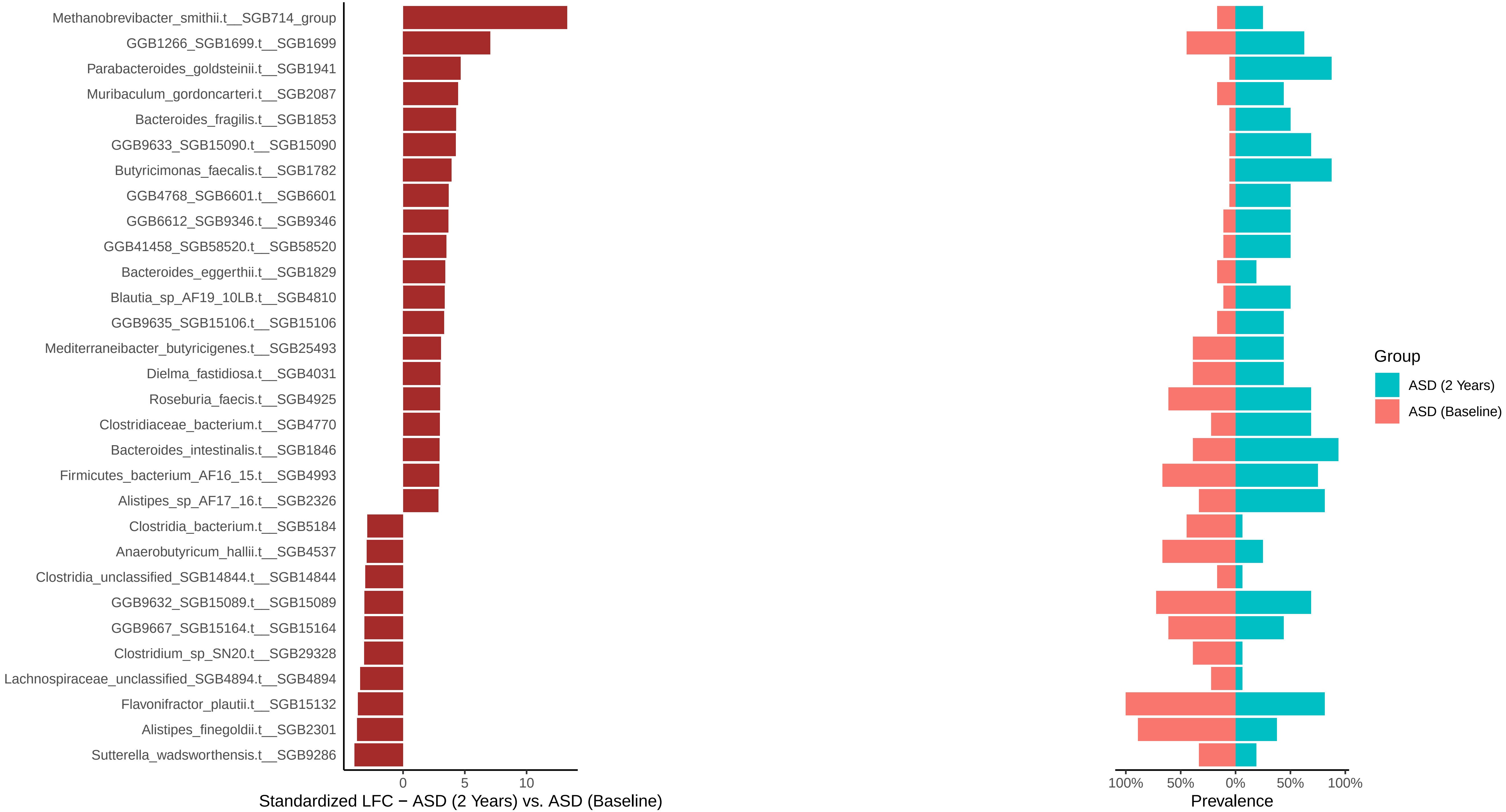

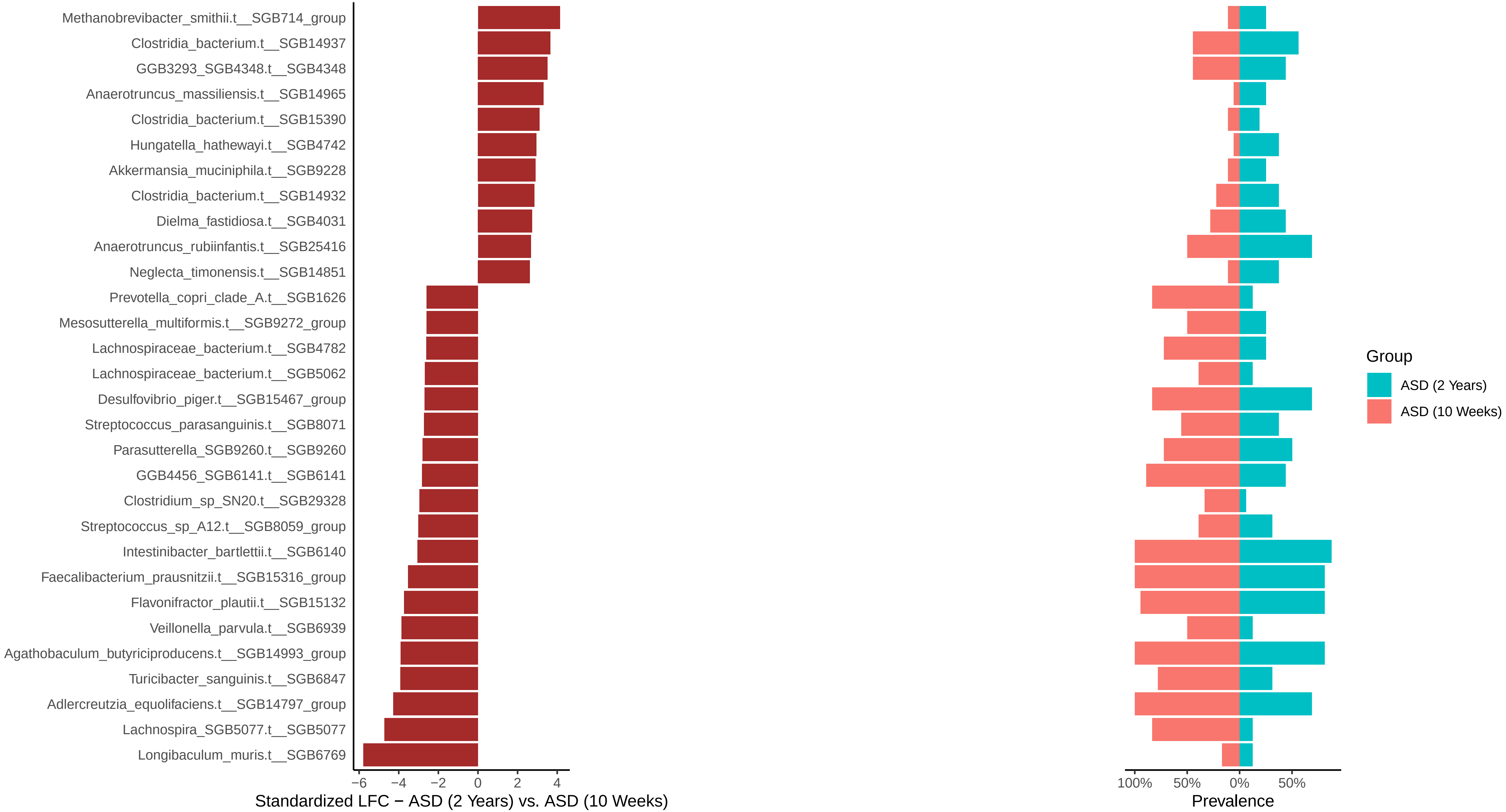

### Figure S5

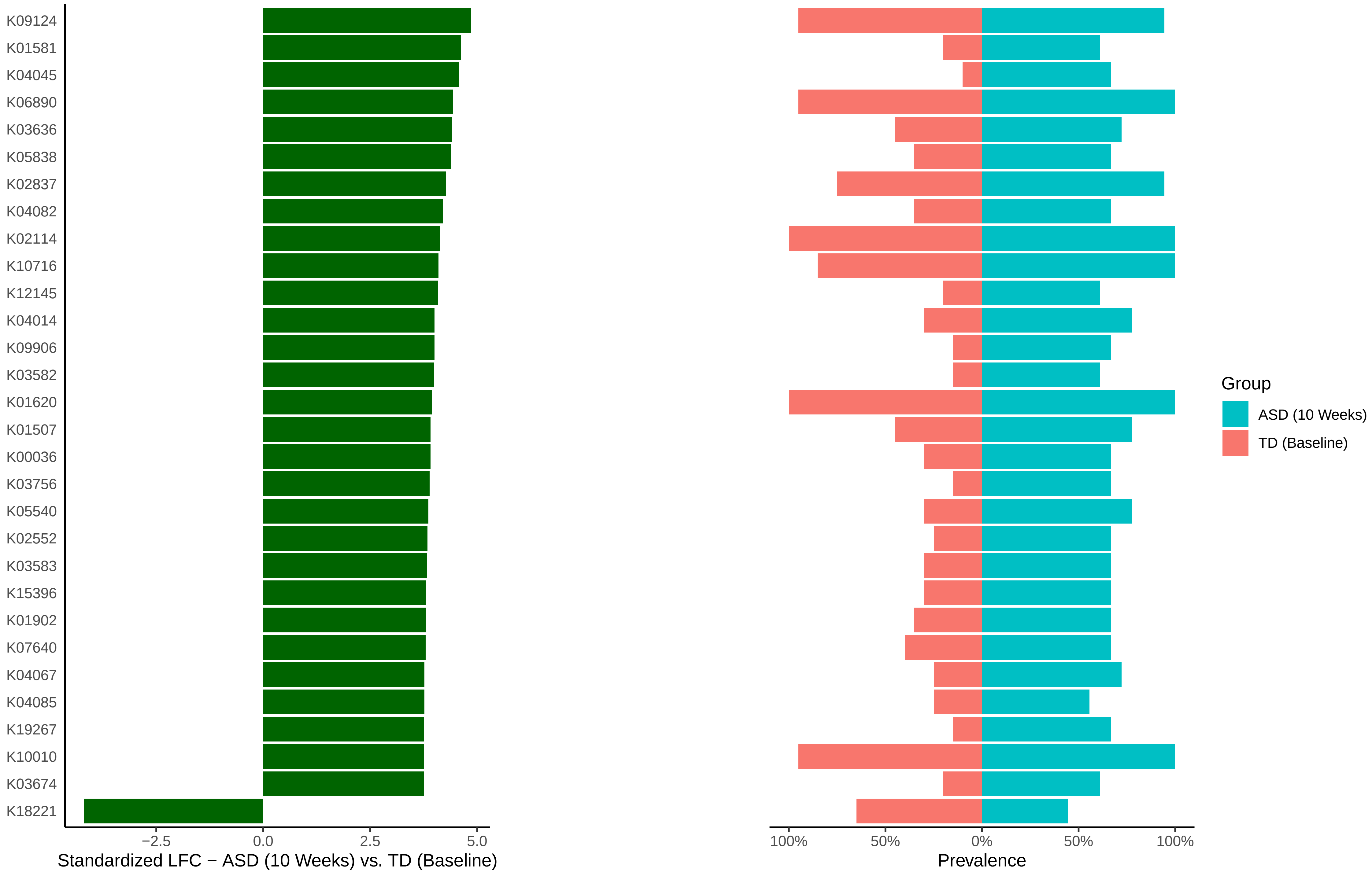

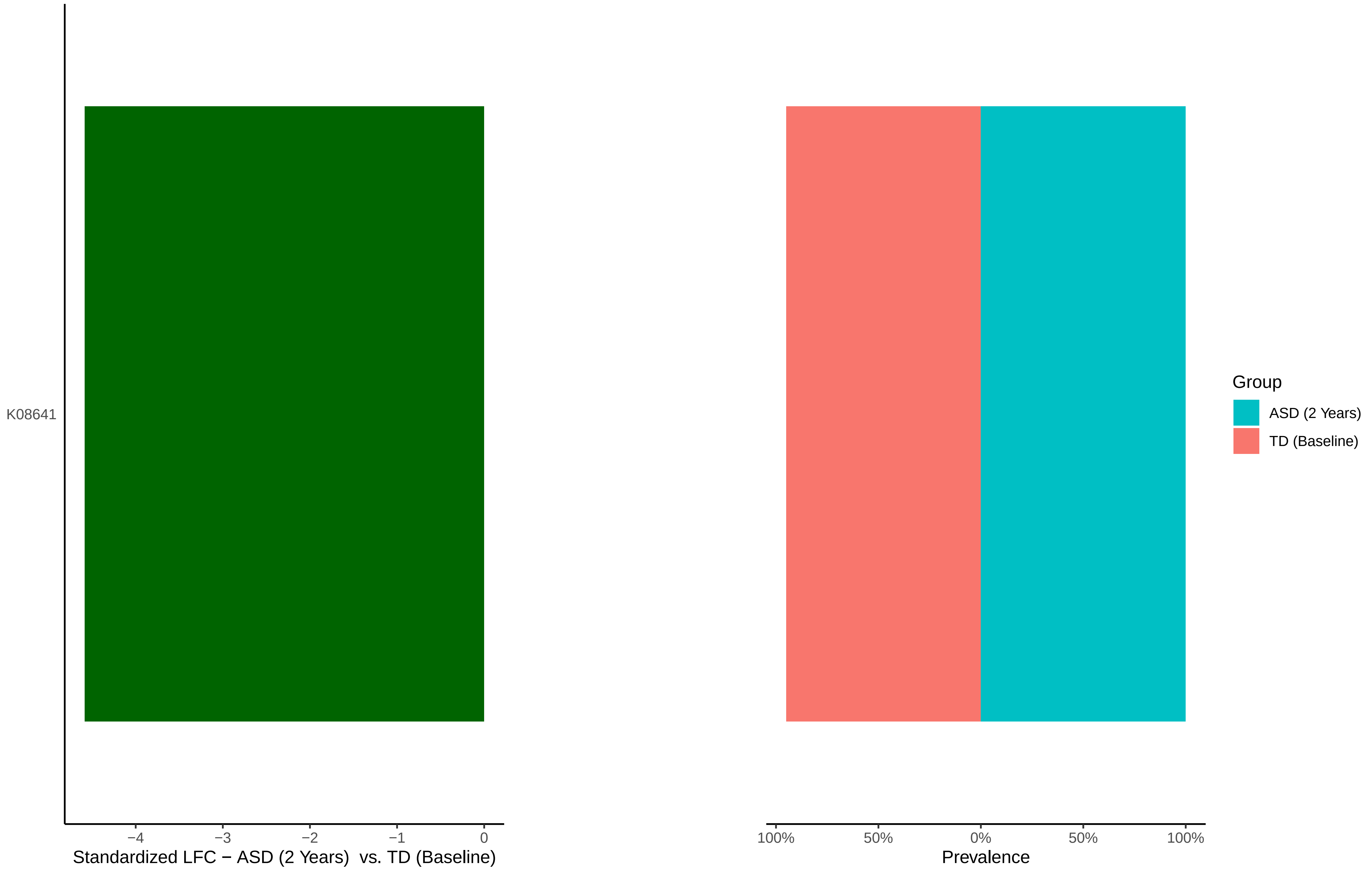

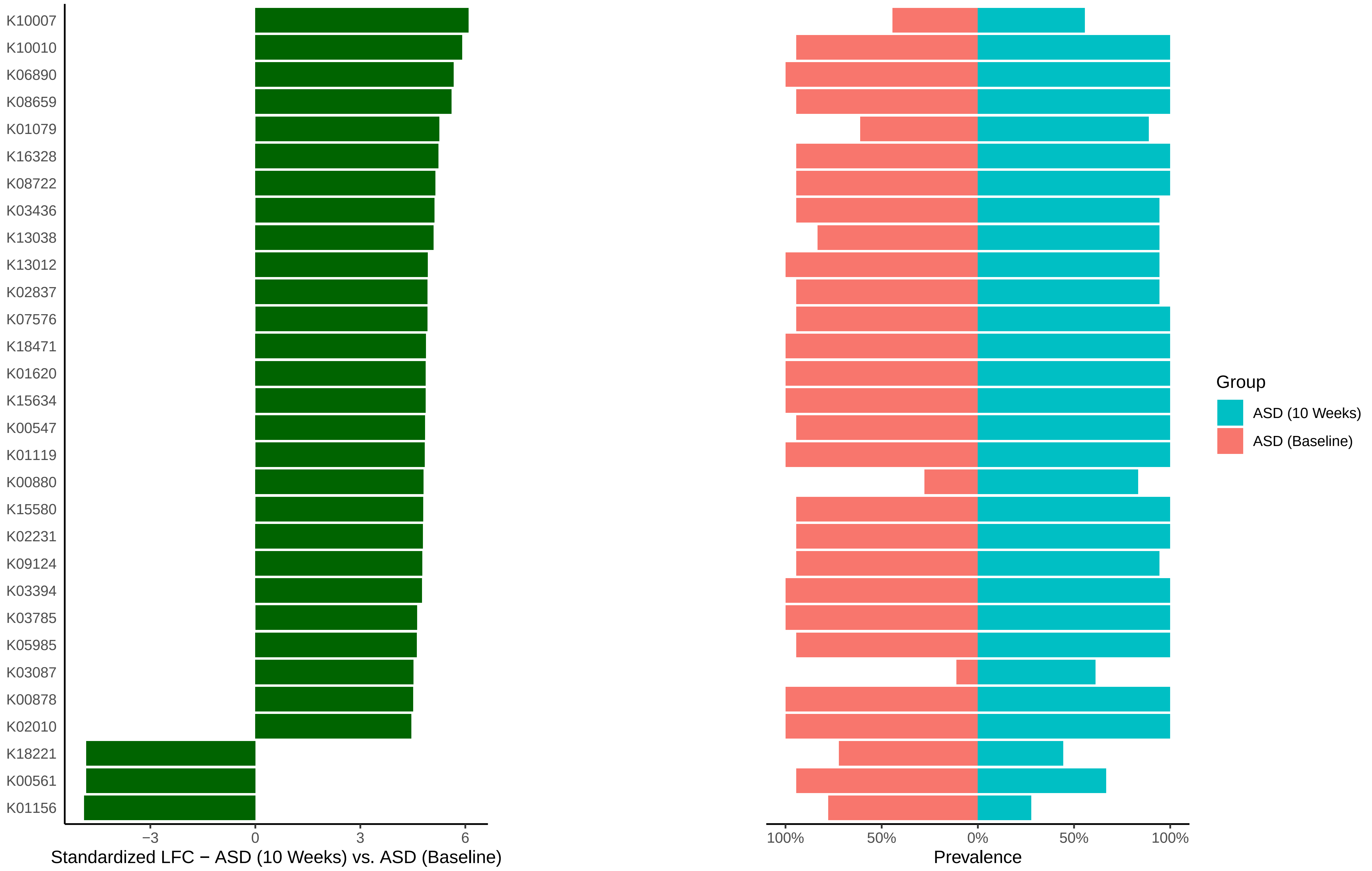

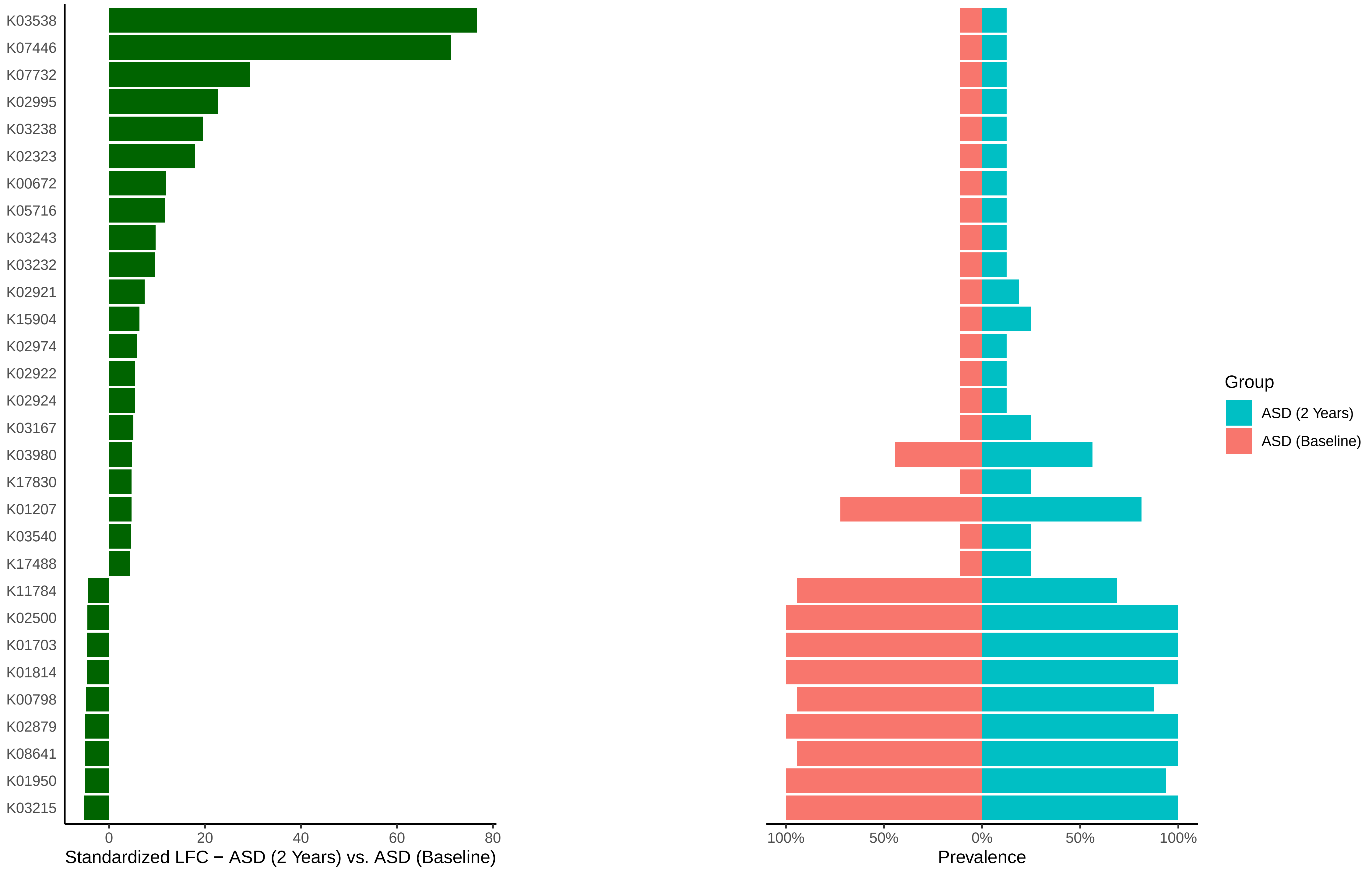

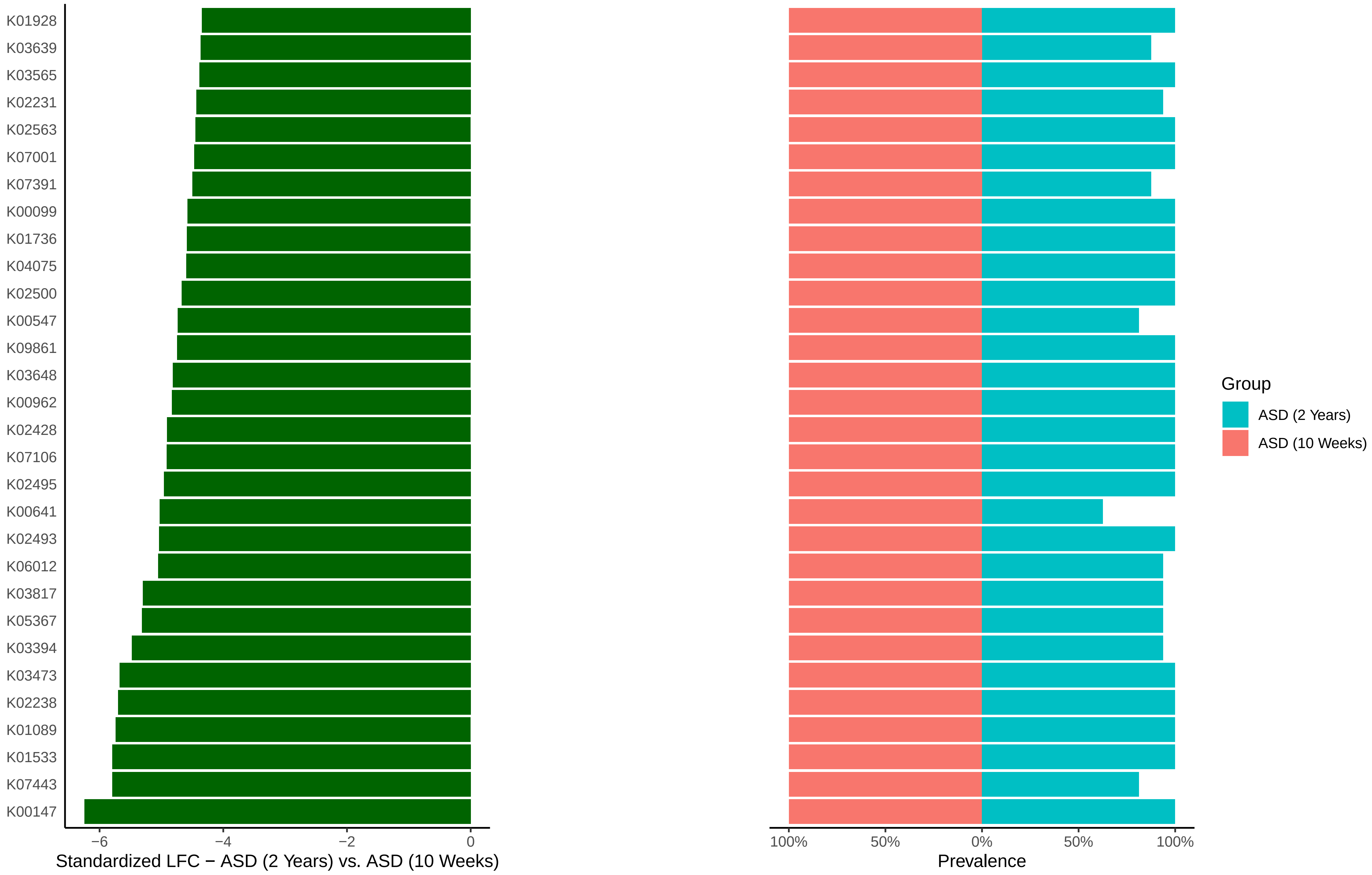

### Figure S6

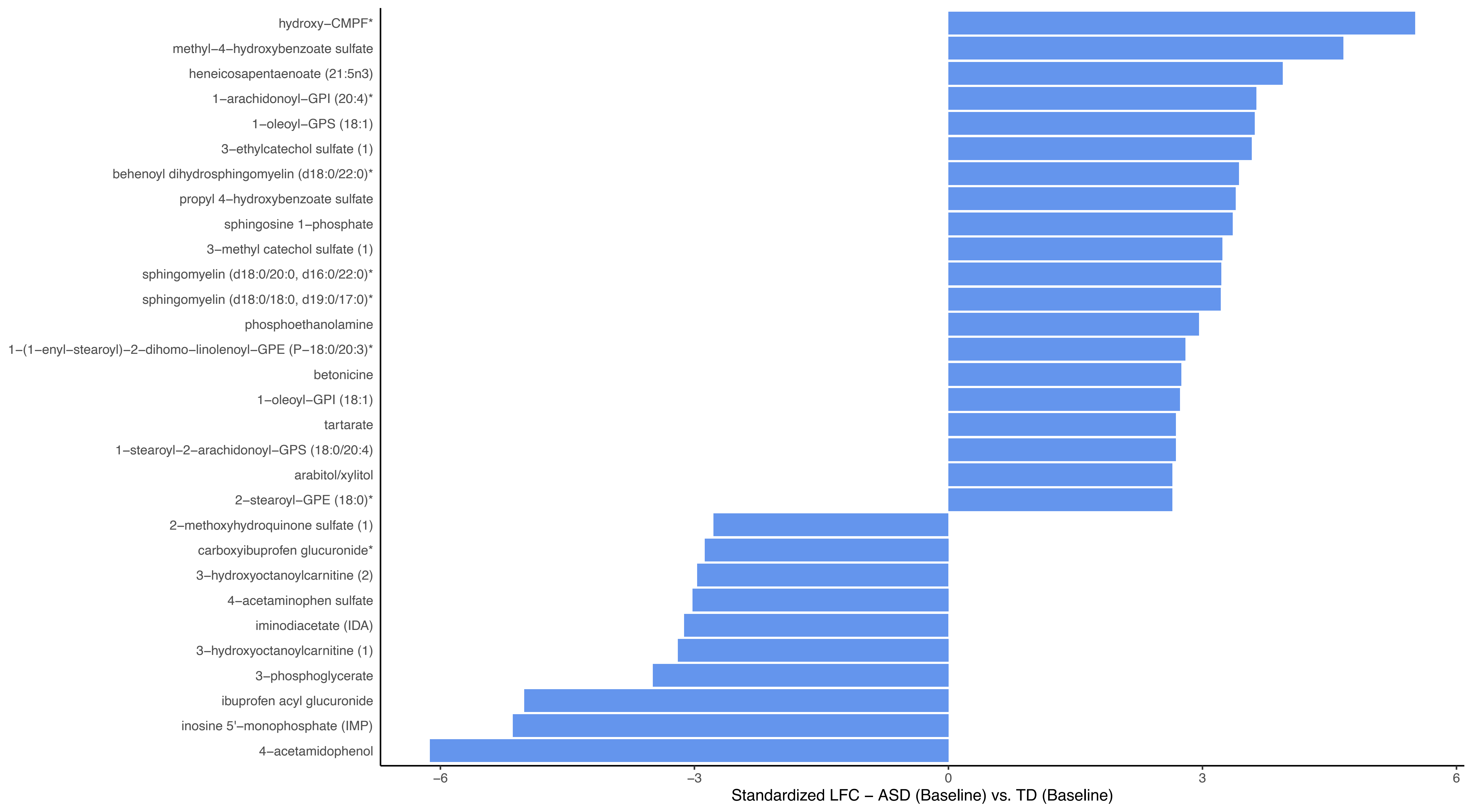

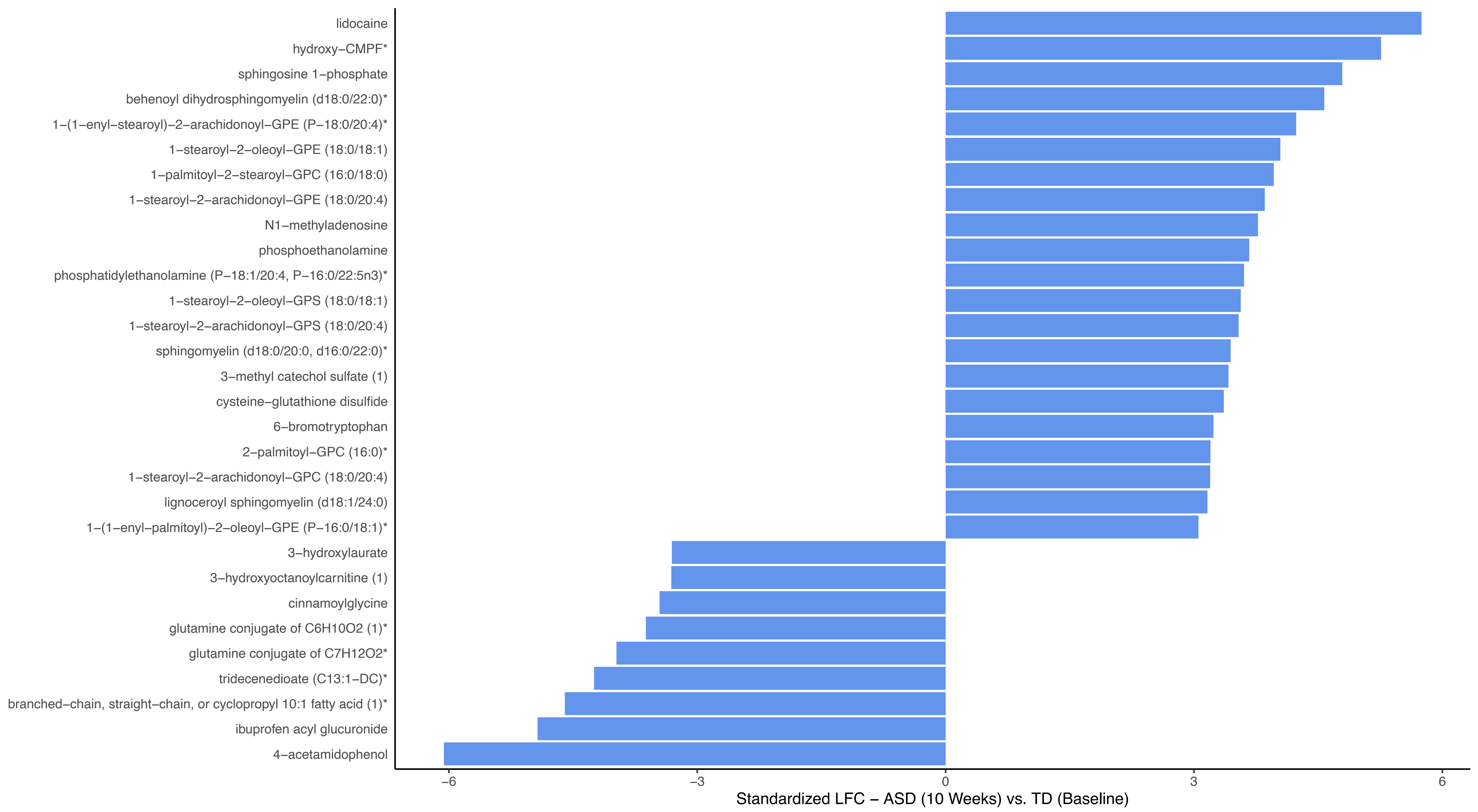

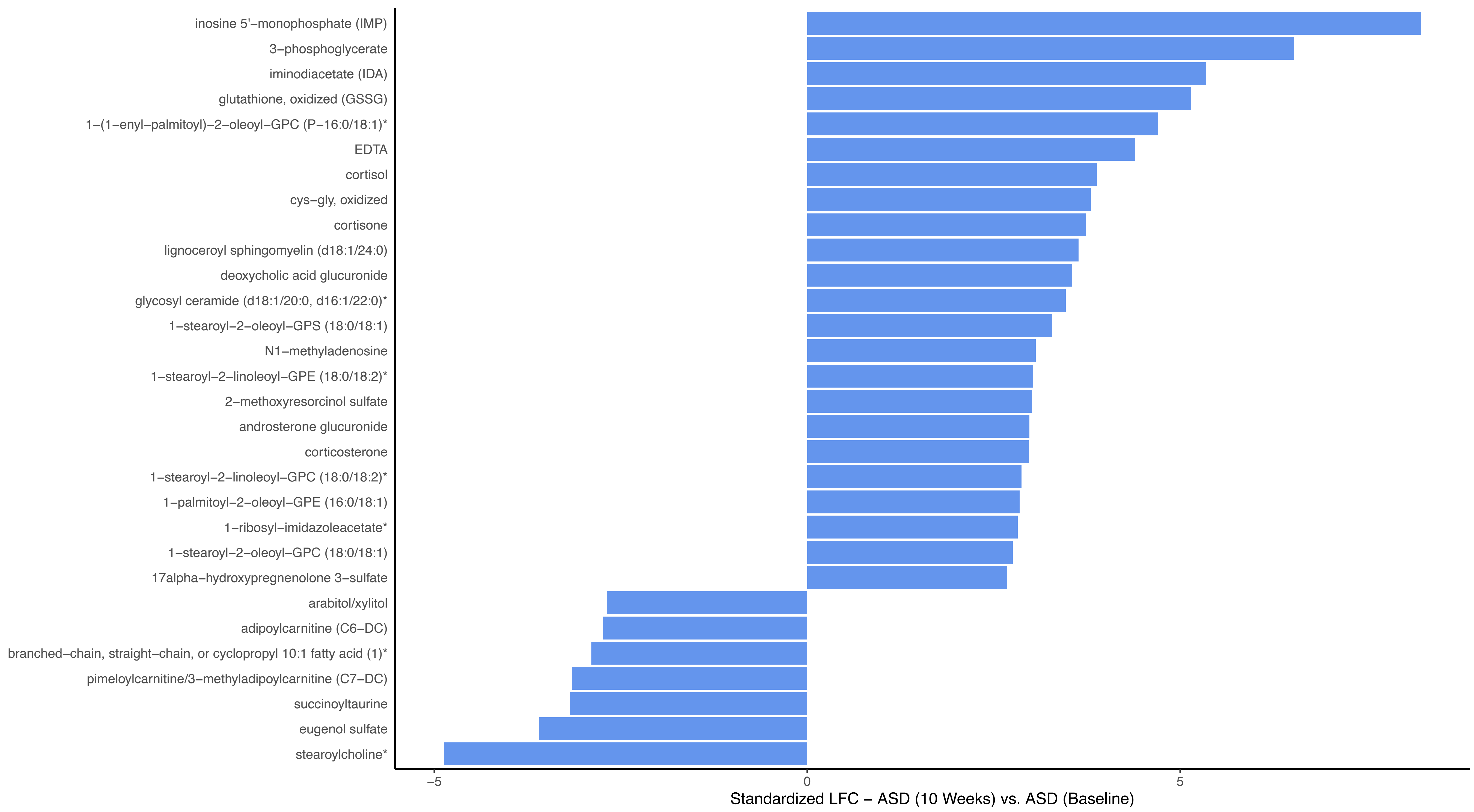

### Figure S7

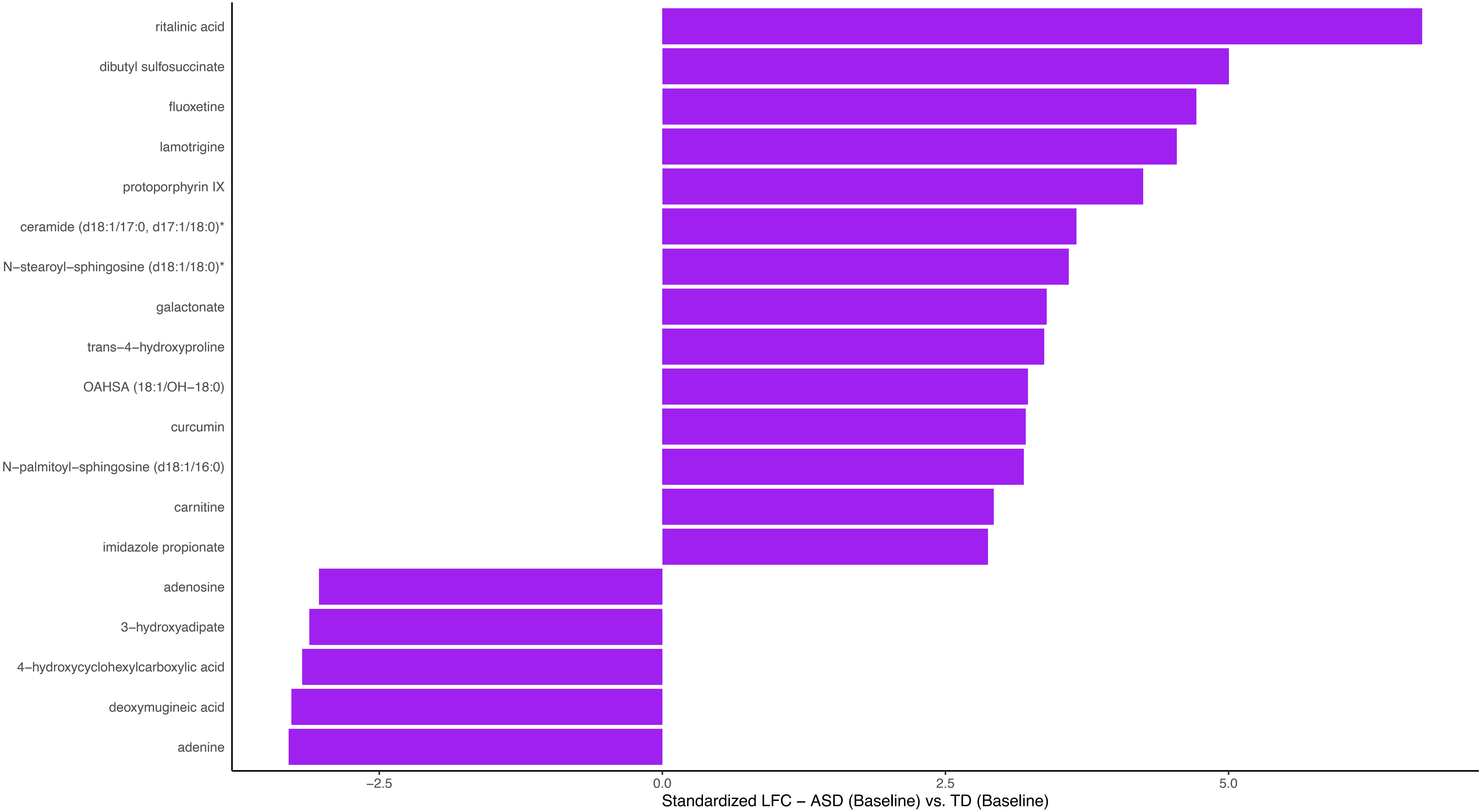

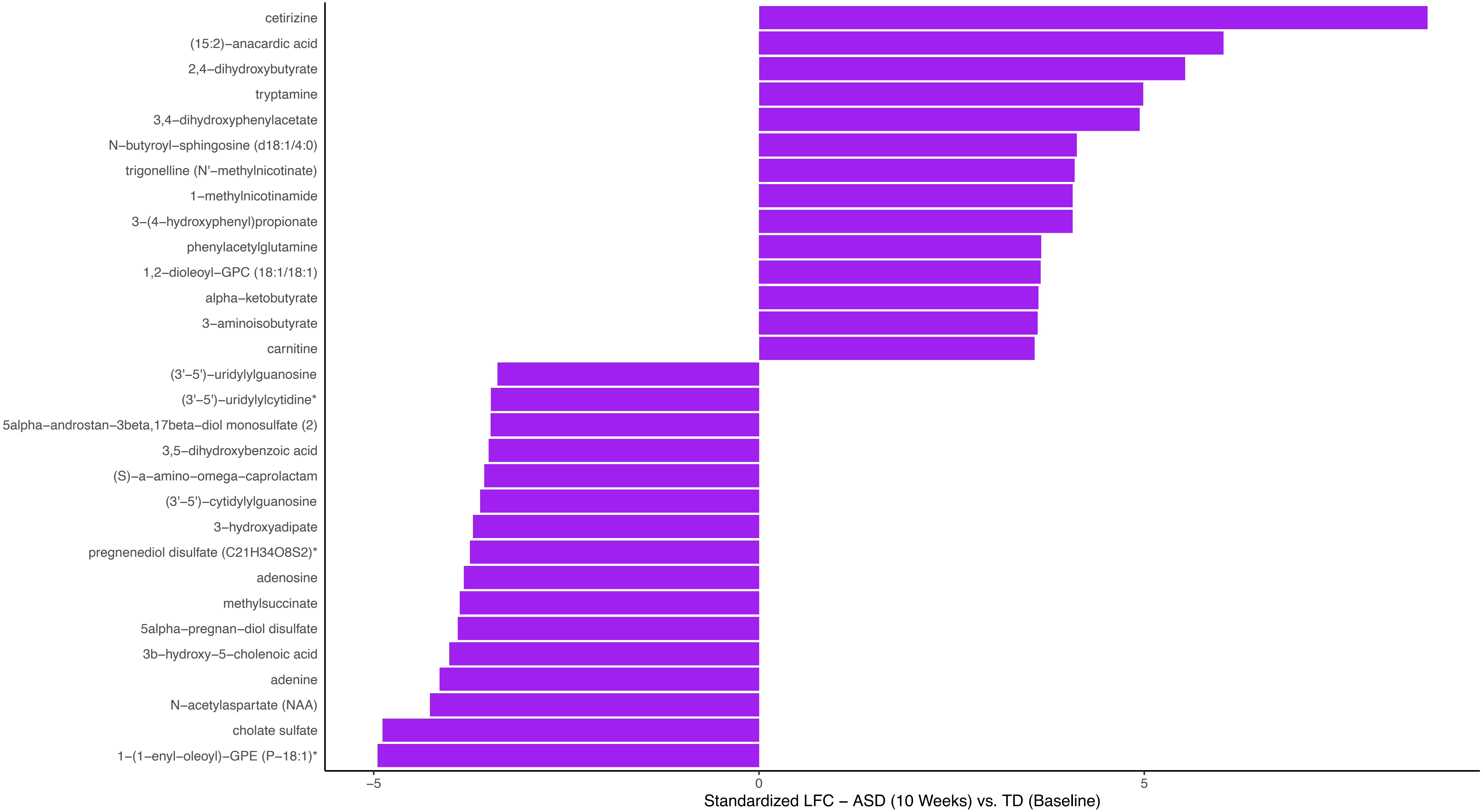

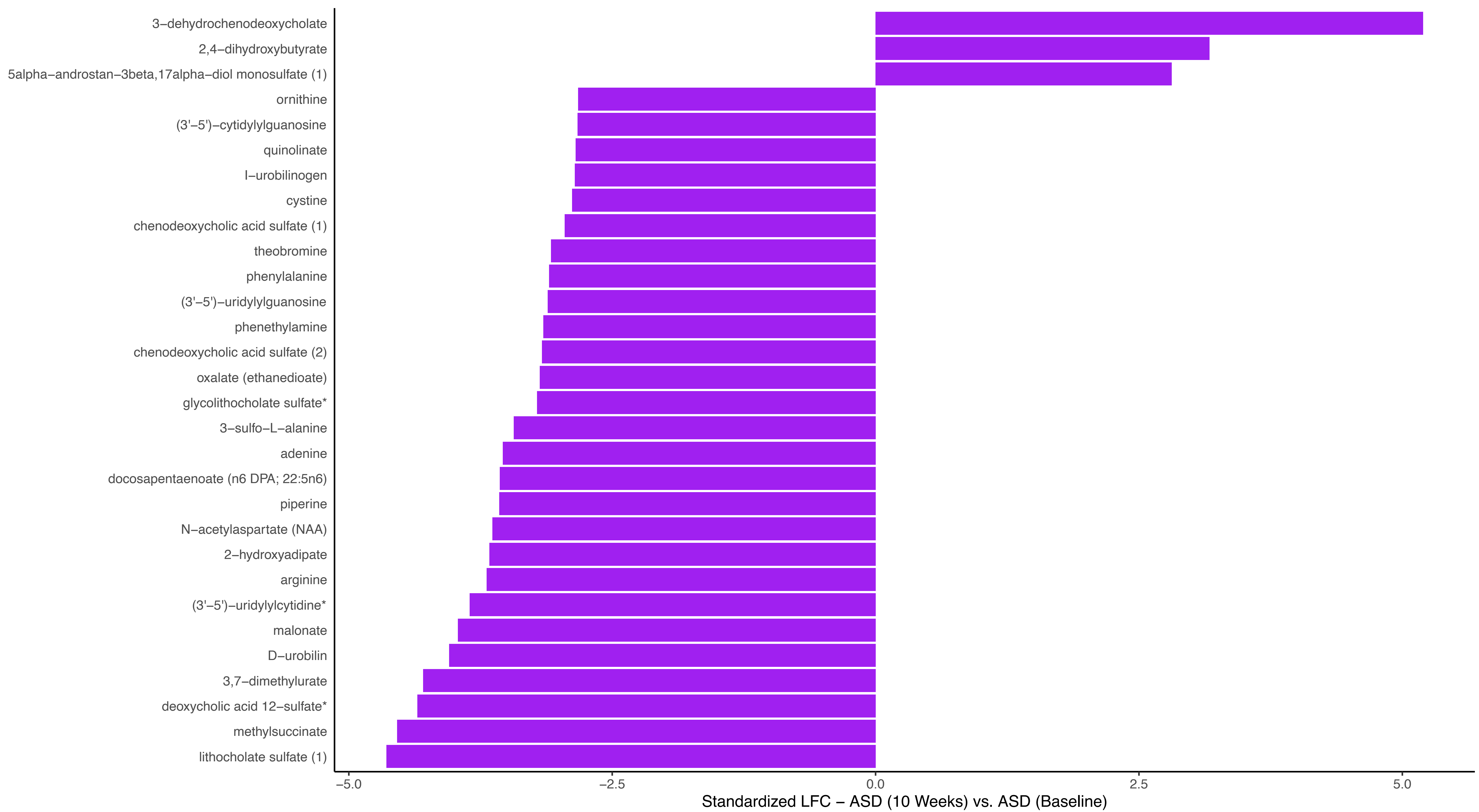

### Figure S8

#### Pre-post network differences in ASD - 10 weeks (L) vs. 2 Years (R)

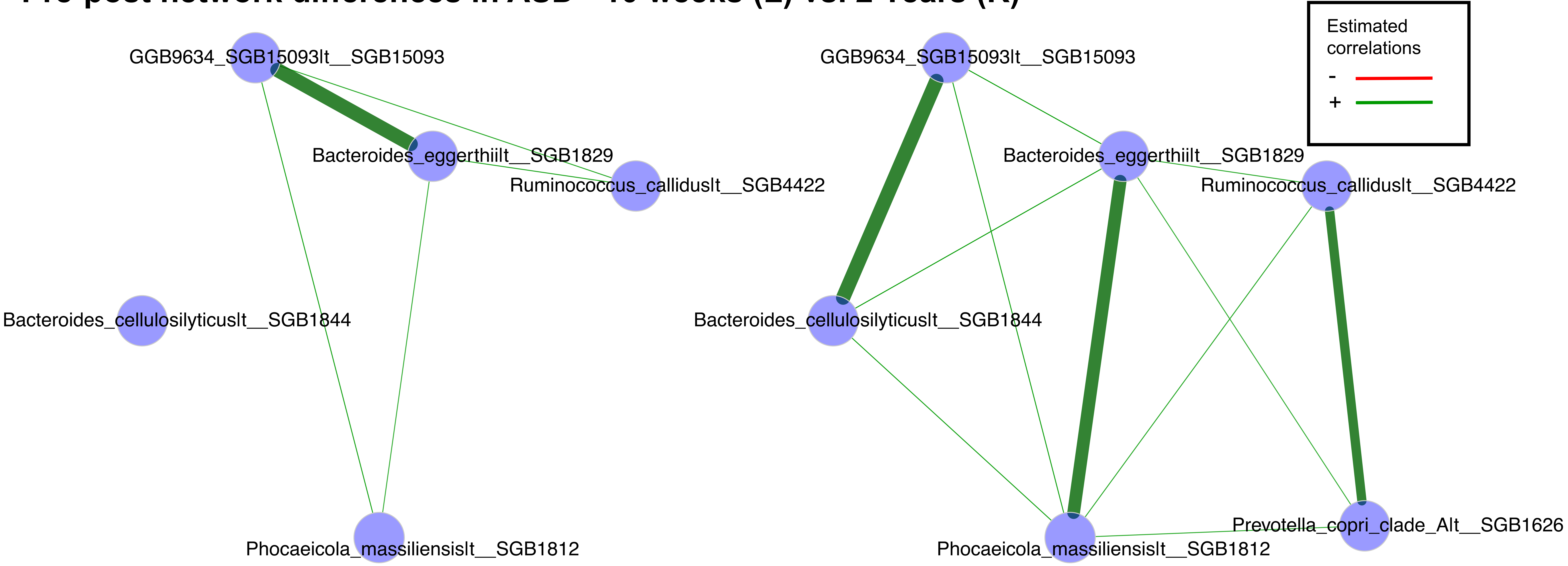
