## Supplementary material for "Multi-omics Integration of Microbiota Transplant Therapy in Children with Autism Spectrum Disorders": Figure S9

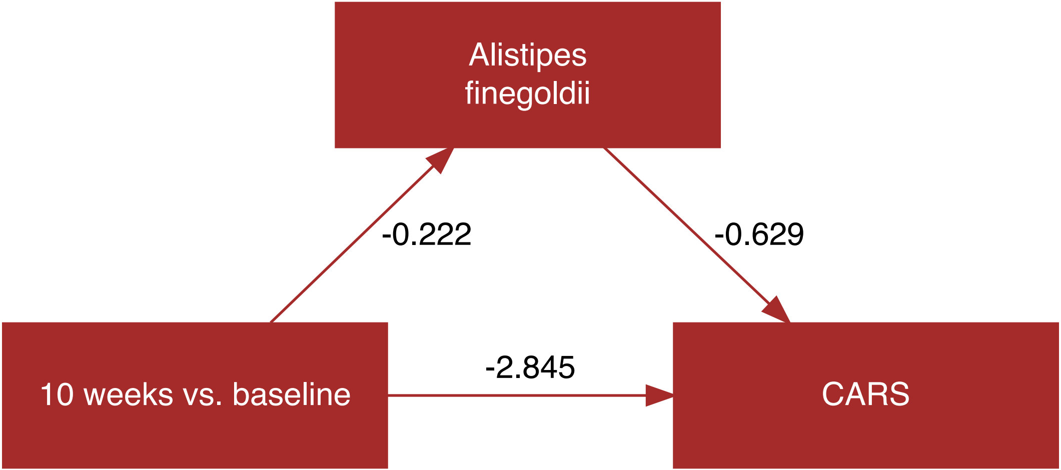
Alistipes finegoldii
-0.222
-0.629
10 weeks vs. baseline
-2.845
CARS

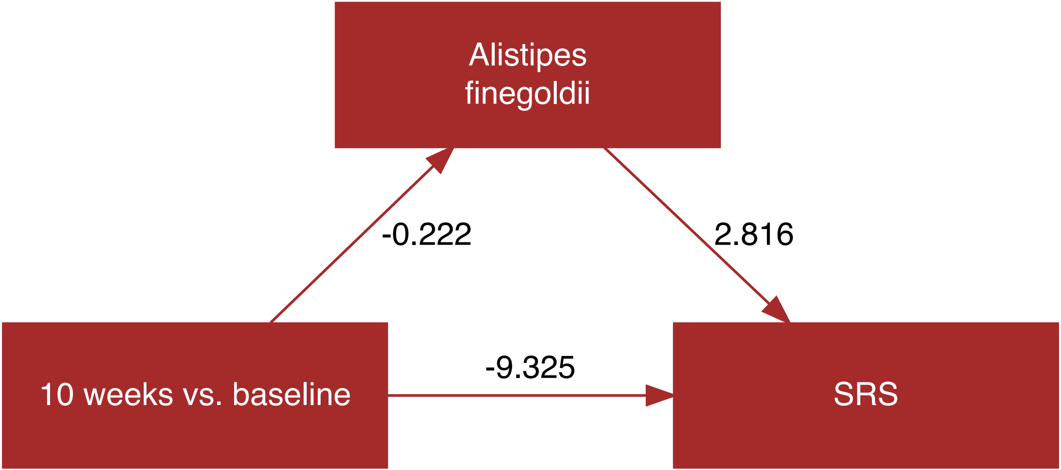
Alistipes finegoldii
-0.222
2.816
10 weeks vs. baseline
-9.325
SRS

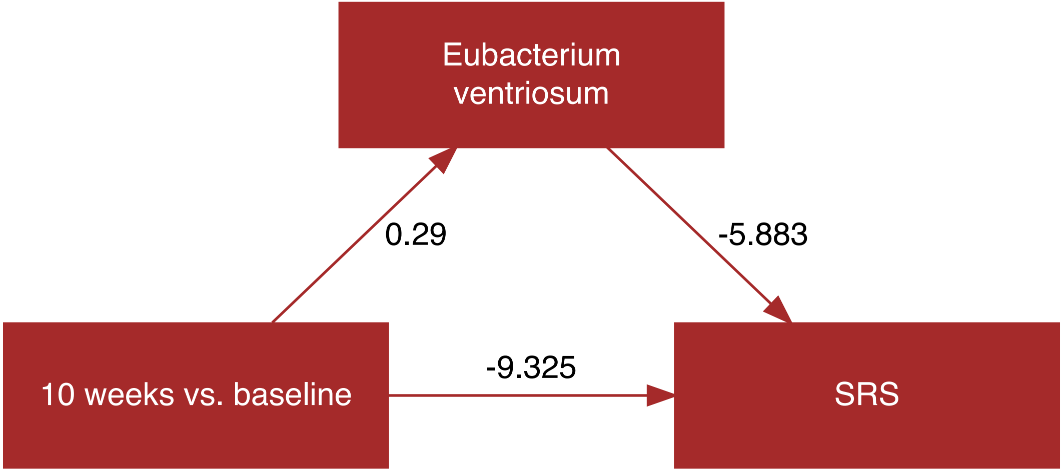
Eubacterium
ventriosum
0.29
-5.883
10 weeks vs. baseline
-9.325
SRS

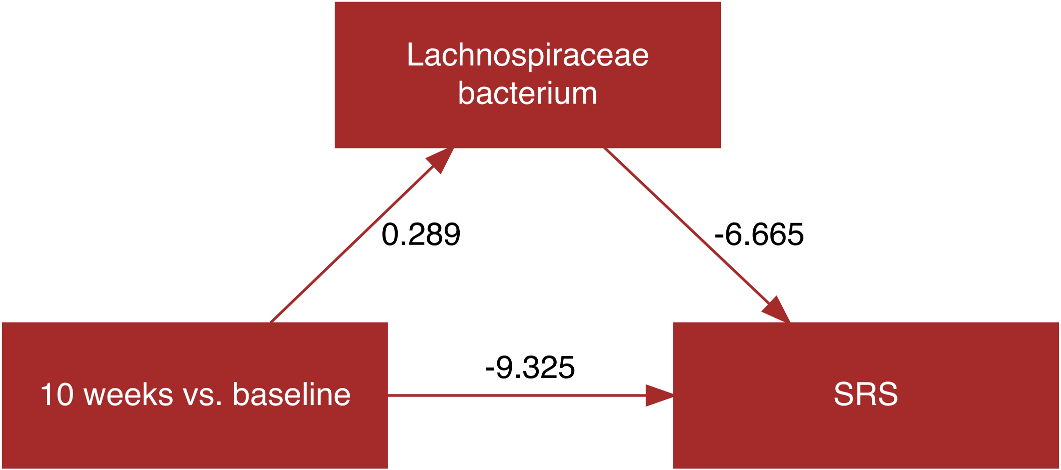
Lachnospiraceae bacterium
0.289
-6.665
-9.325
10 weeks vs. baseline
SRS

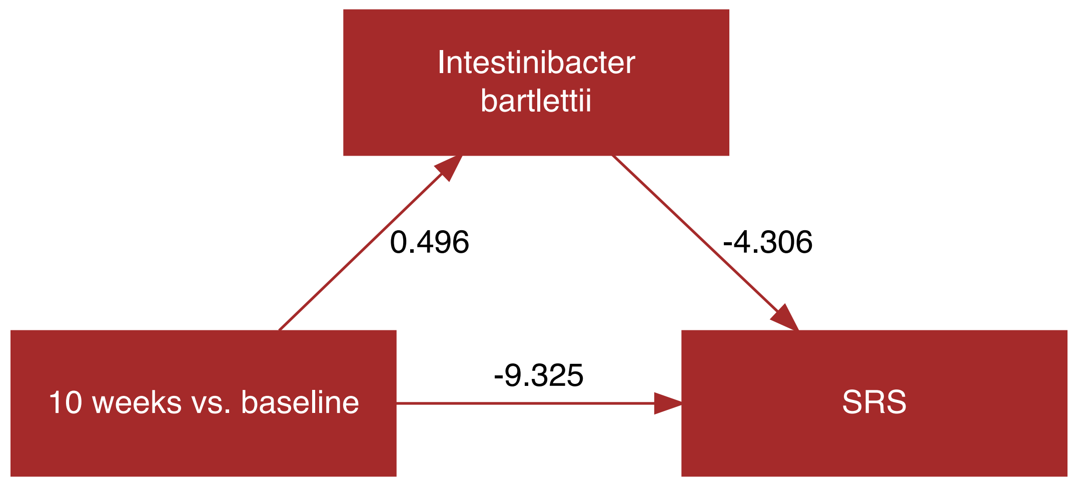
Intestinibacter bartlettii
0.496
-4.306
-9.325
10 weeks vs. baseline
SRS

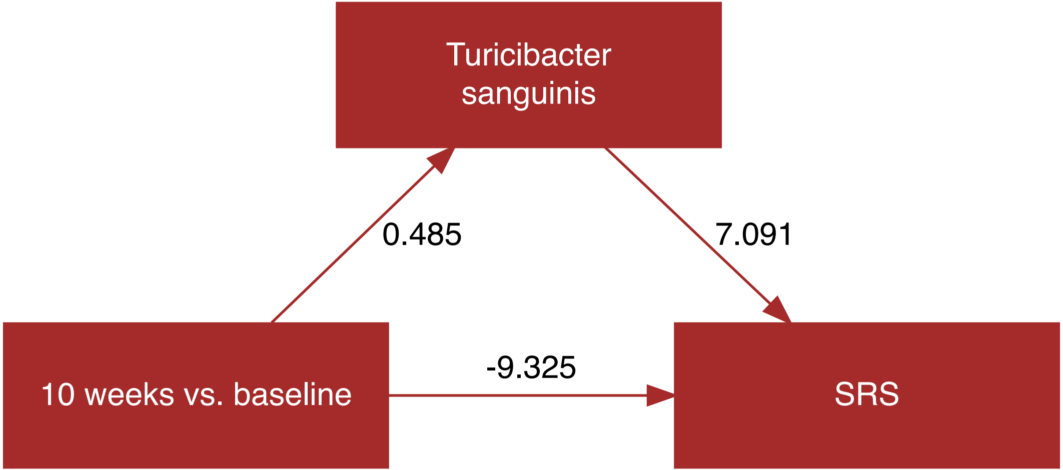
Turicibacter
sanguinis
0.485
7.091
-9.325
10 weeks vs. baseline
SRS

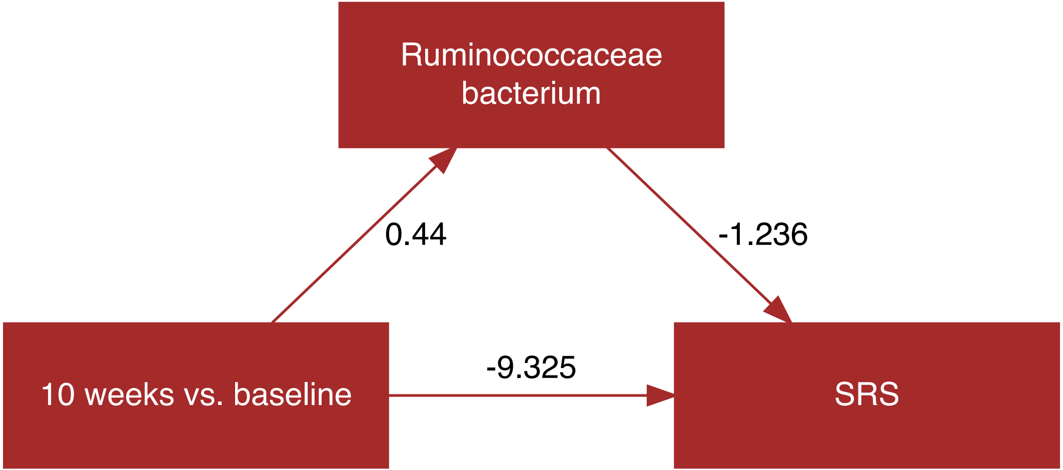
Ruminococcaceae bacterium
0.44
-1.236
-9.325
10 weeks vs. baseline
SRS

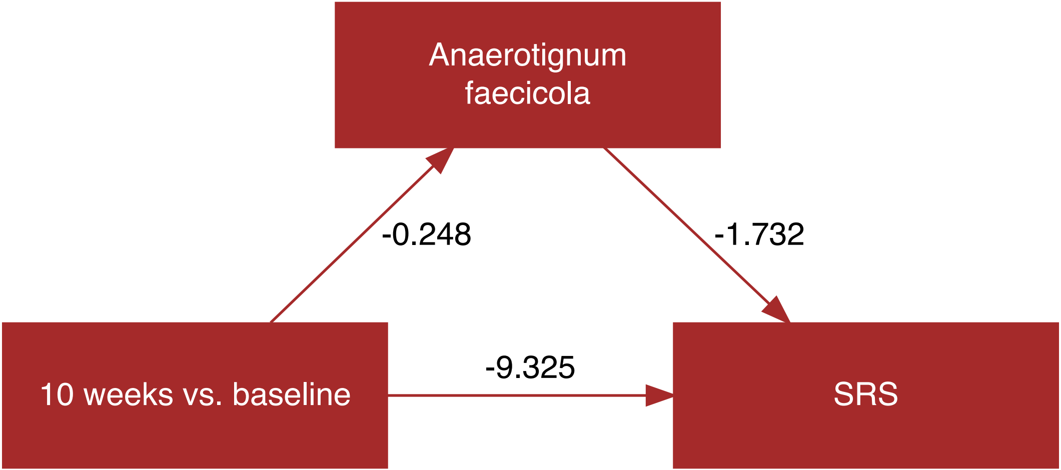
Anaerotignum faecicola
-0.248
-1.732
-9.325
10 weeks vs. baseline
SRS

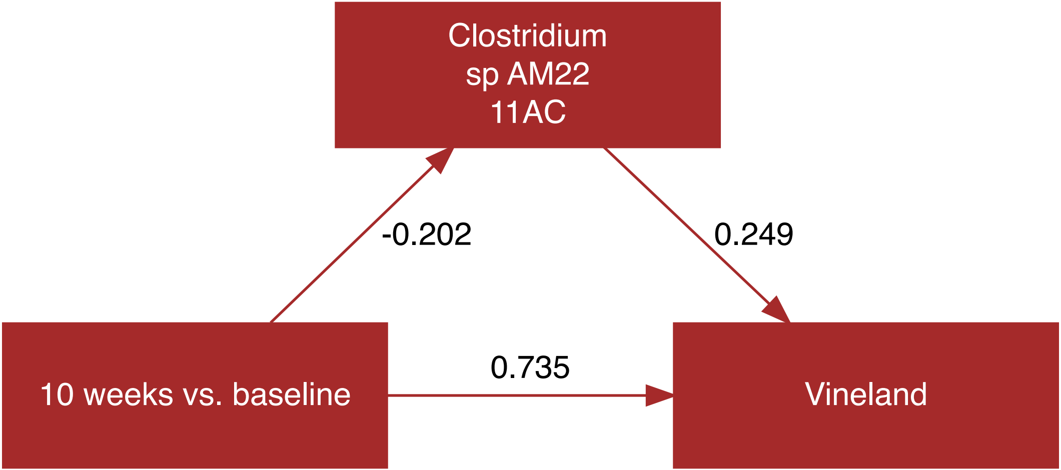
Clostridium
sp AM22
11AC
-0.202
0.249
0.735
10 weeks vs. baseline
Vineland

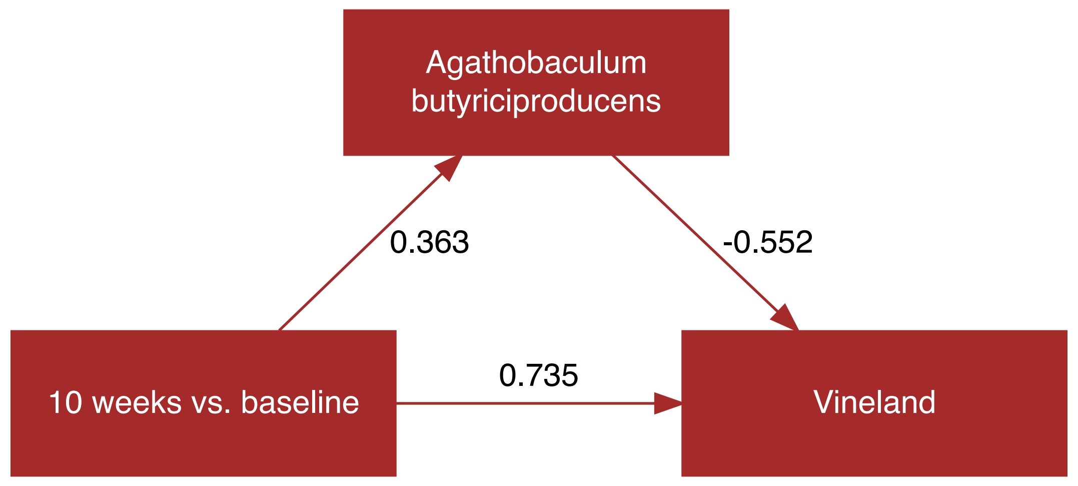
Agathobaculum butyriciproducens
0.363
-0.552
10 weeks vs. baseline
0.735
Vineland

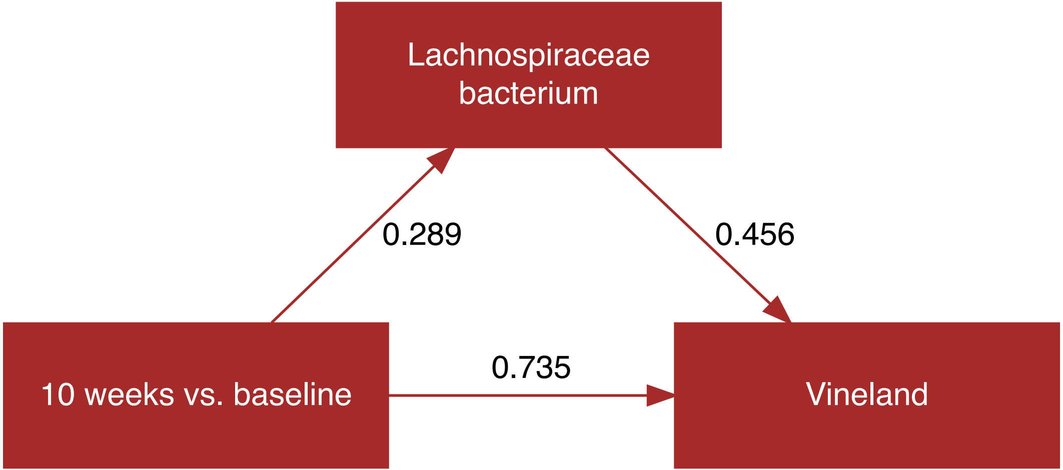
Lachnospiraceae
bacterium
0.289
0.456
10 weeks vs. baseline
0.735
Vineland

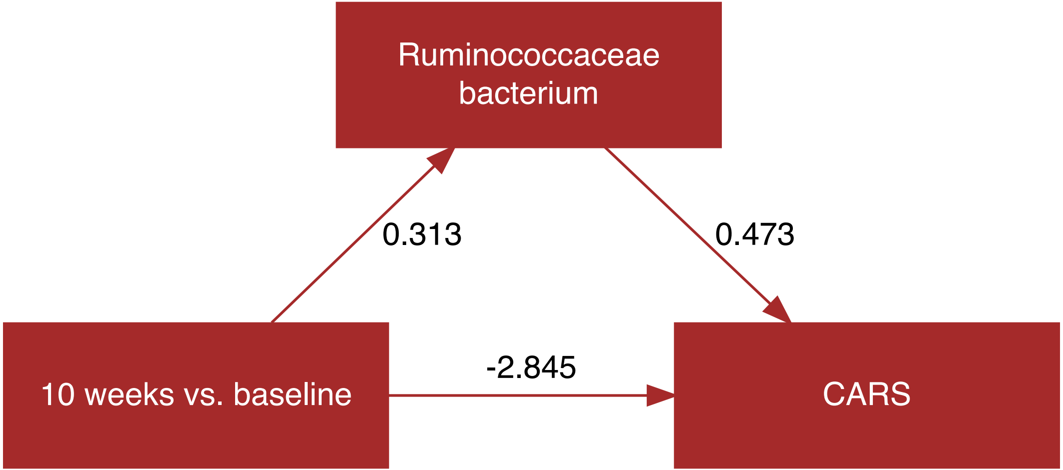
Ruminococcaceae bacterium
0.313
0.473
-2.845
10 weeks vs. baseline
CARS

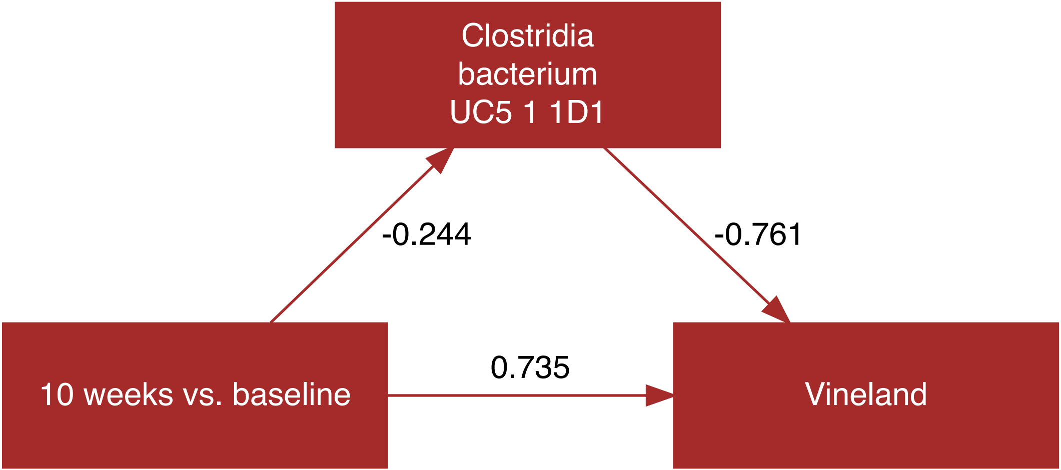
Clostridia
bacterium
UC5 1 1D1
-0.244
-0.761
0.735
10 weeks vs. baseline
Vineland

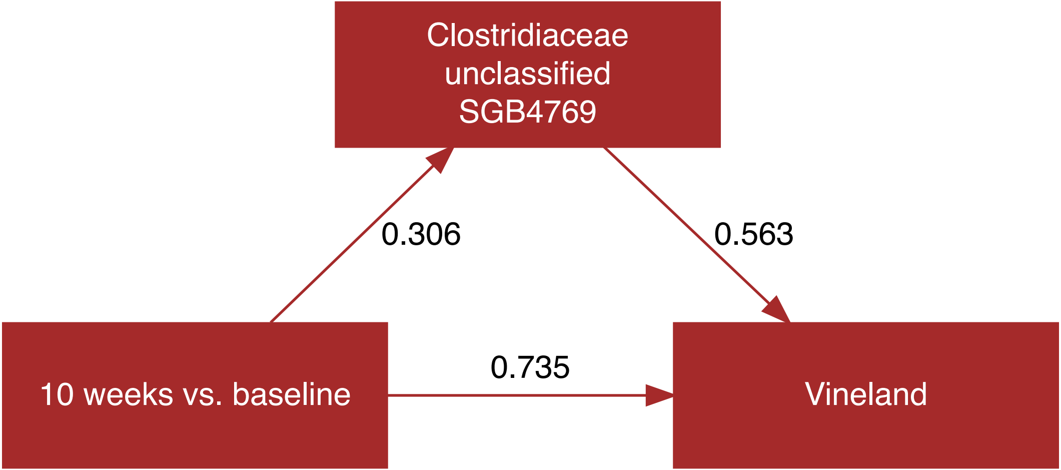

Clostridiaceae unclassified SGB4769
0.306
0.563
10 weeks vs. baseline
0.735
Vineland

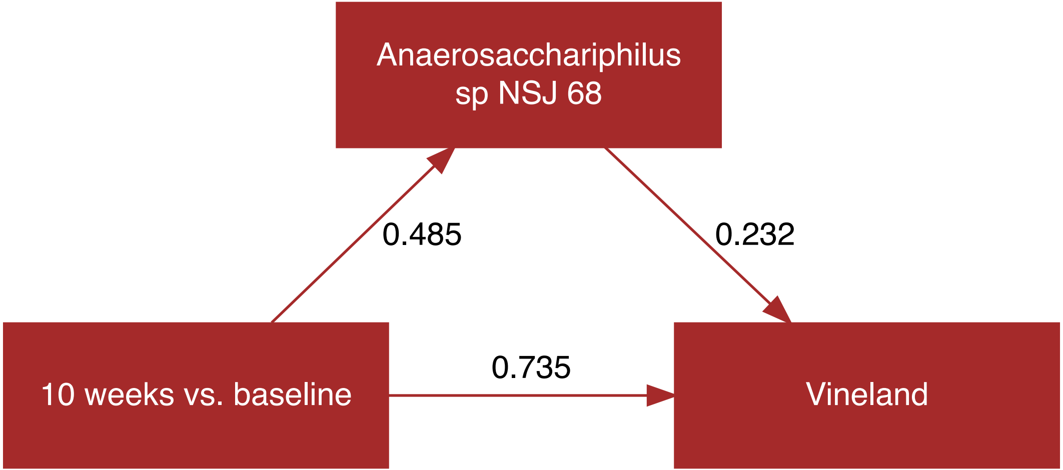
Anaerosacchariphilus
sp NSJ 68
0.485
0.232
10 weeks vs. baseline
0.735
Vineland

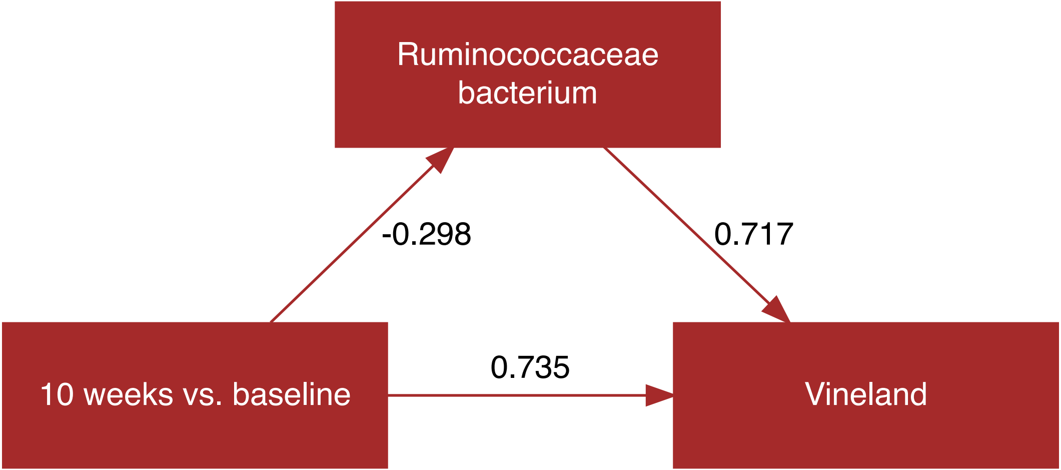
Ruminococcaceae bacterium
-0.298
0.717
10 weeks vs. baseline
0.735
Vineland

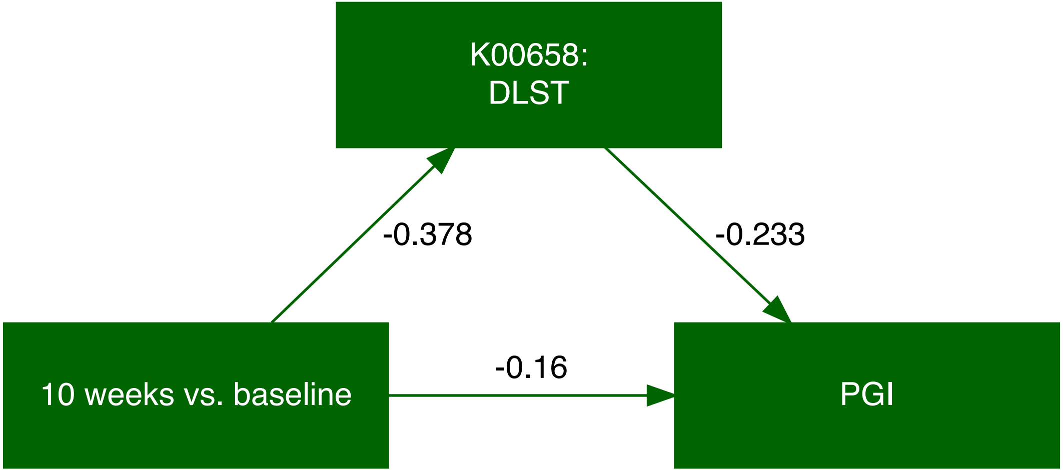
K00658:
DLST
-0.378
-0.233
10 weeks vs. baseline
-0.16
PGI

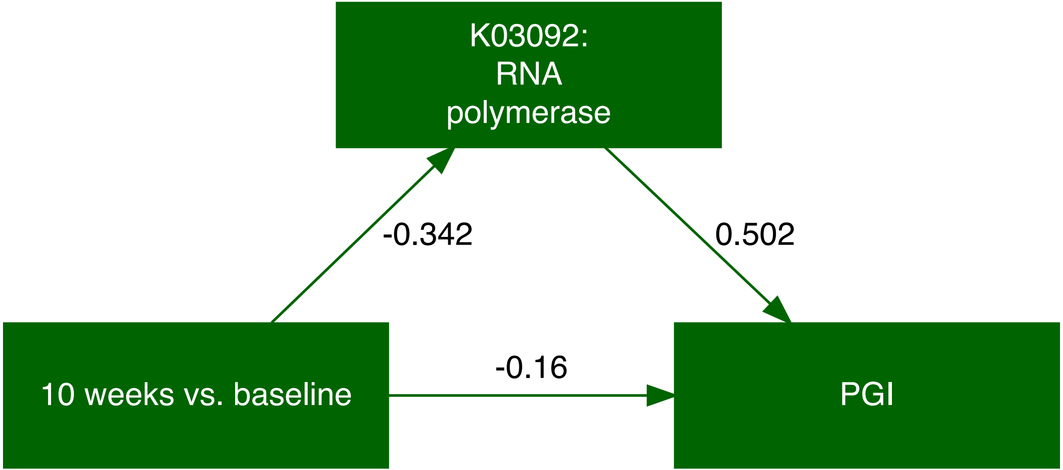
K03092:
RNA
polymerase
-0.342
0.502
-0.16
10 weeks vs. baseline
PGI

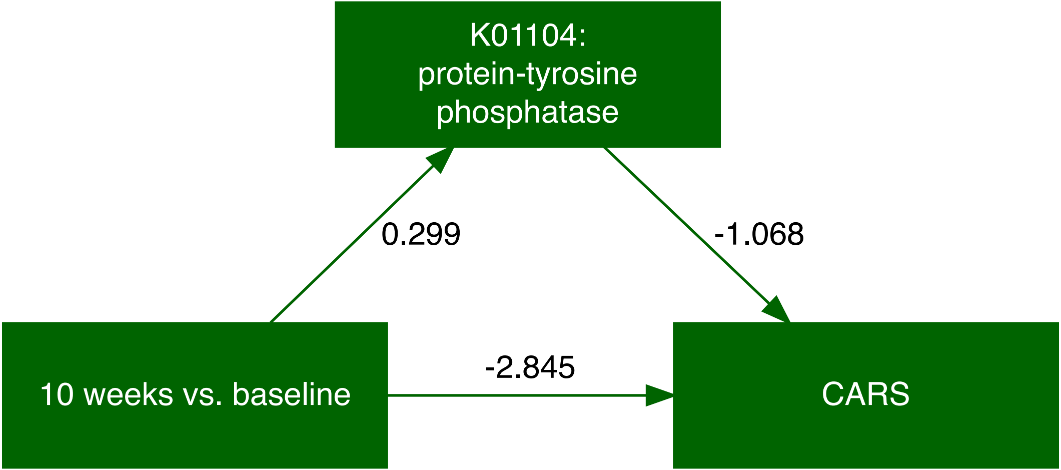
K01104:
protein-tyrosine
phosphatase
0.299
-1.068
10 weeks vs. baseline
-2.845
CARS

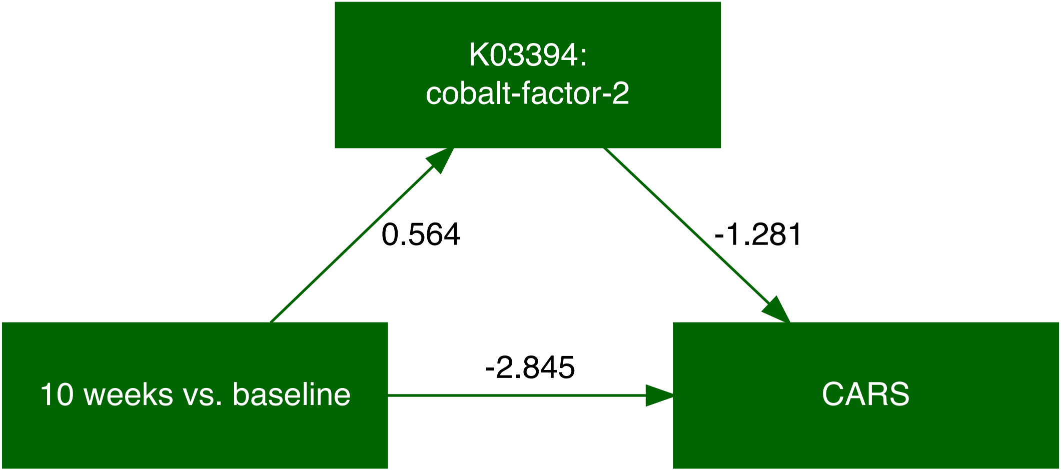
K03394:
cobalt-factor-2
0.564
-1.281
-2.845
10 weeks vs. baseline
CARS

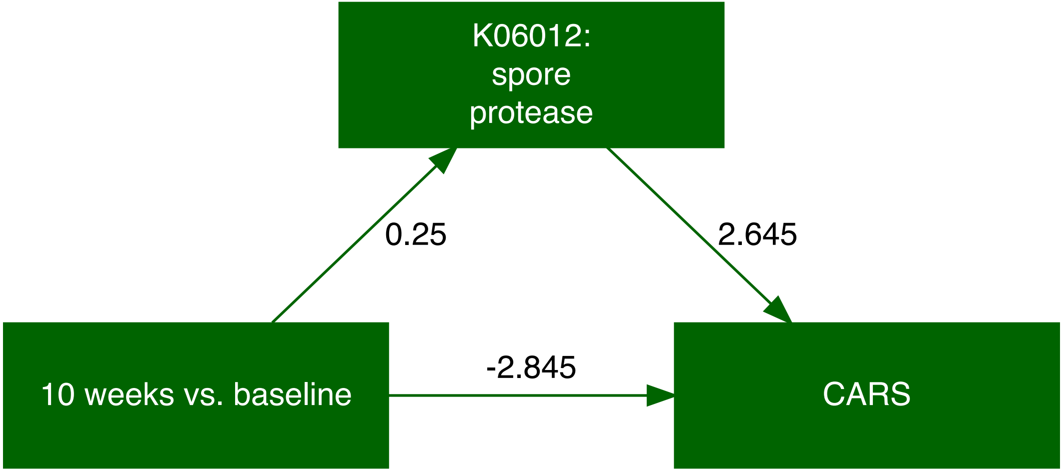
K06012:
spore
protease
0.25
2.645
-2.845
10 weeks vs. baseline
CARS

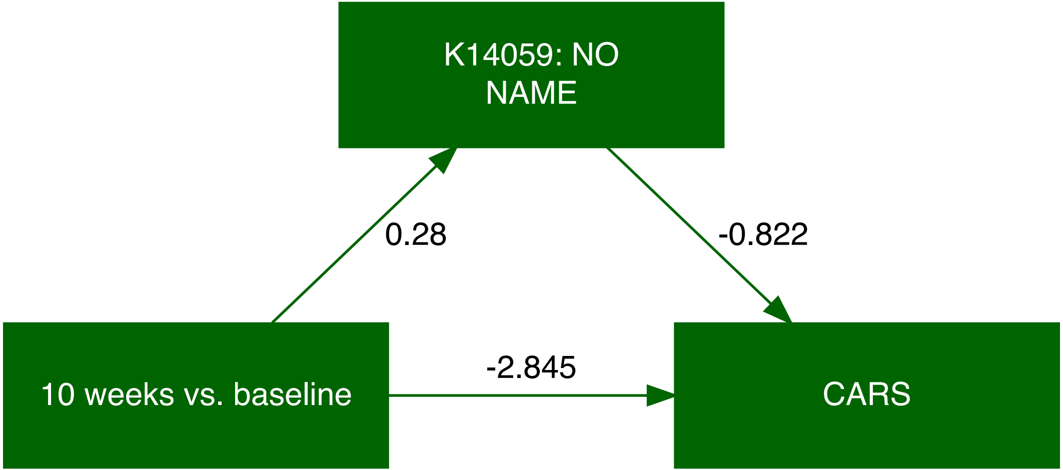
K14059: NO
NAME
0.28
-0.822
-2.845
10 weeks vs. baseline
CARS

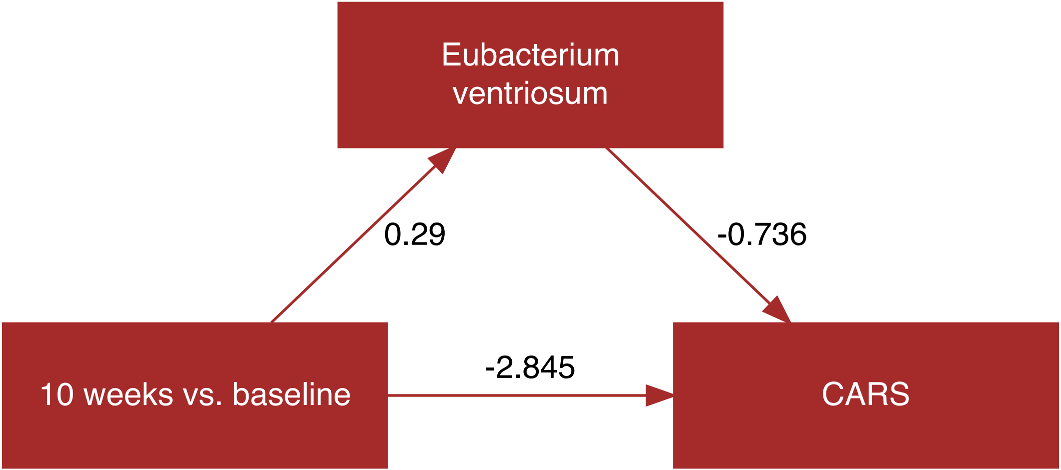
Eubacterium ventriosum
0.29
-0.736
10 weeks vs. baseline
-2.845
CARS

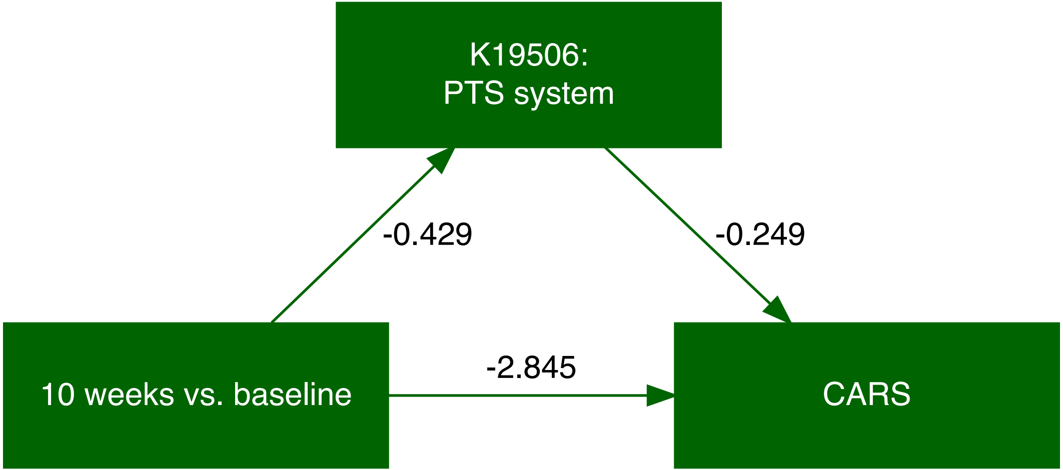
K19506:
PTS system
-0.429
-0.249
10 weeks vs. baseline
-2.845
CARS

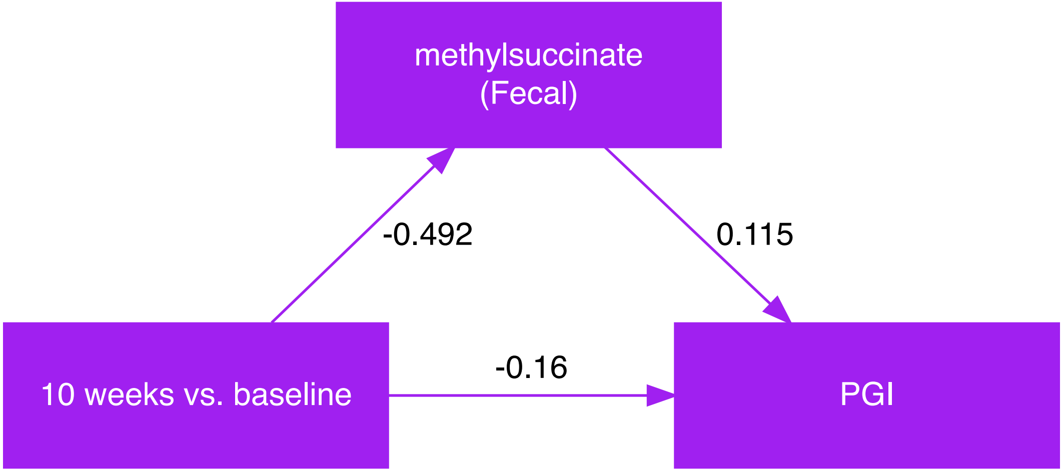
methylsuccinate
(Fecal)
-0.492
0.115
10 weeks vs. baseline
-0.16
PGI

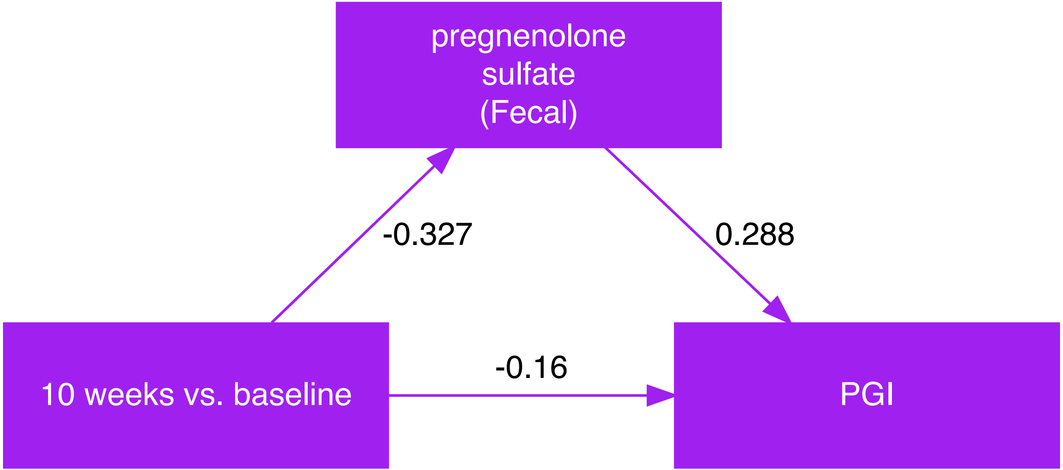
pregnenolone
sulfate
(Fecal)
-0.327
0.288
-0.16
10 weeks vs. baseline
PGI

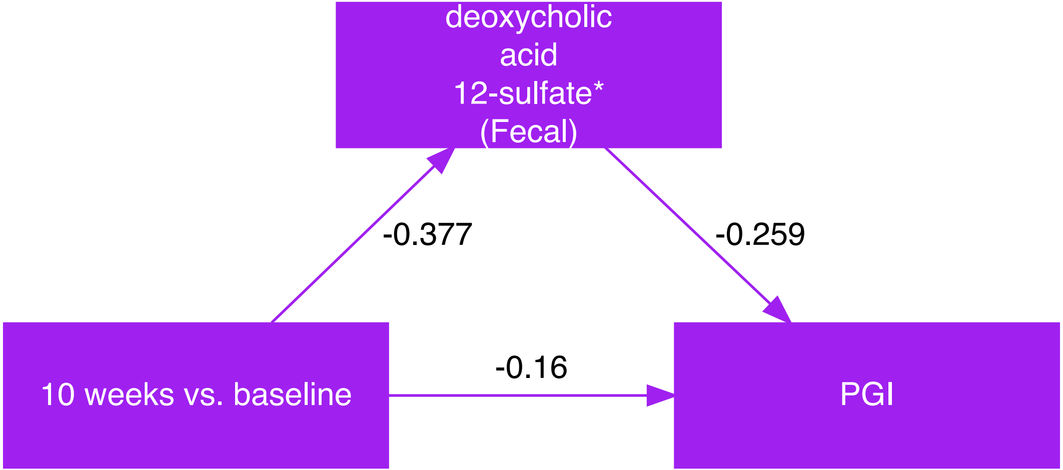
deoxycholic
acid
12-sulfate*
(Fecal)
-0.377
-0.259
-0.16
10 weeks vs. baseline
PGI

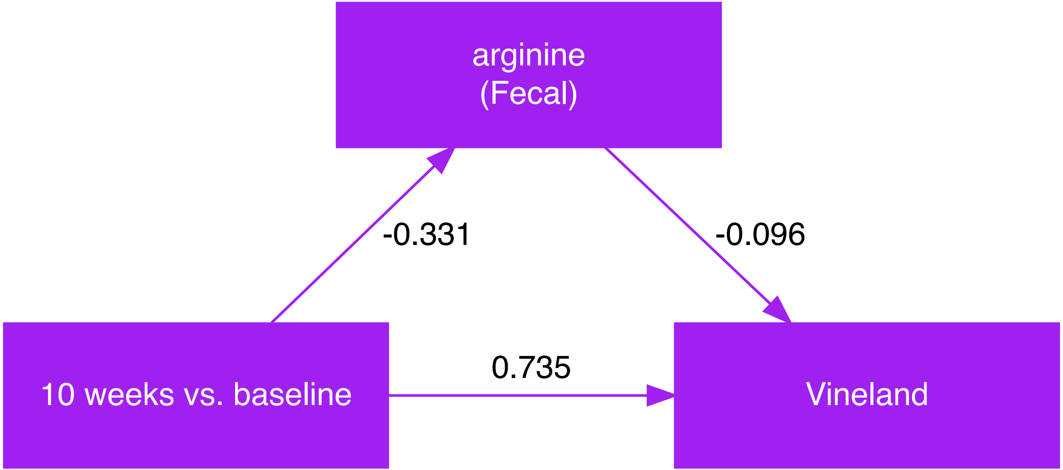
arginine
(Fecal)
-0.331
-0.096
0.735
10 weeks vs. baseline
Vineland

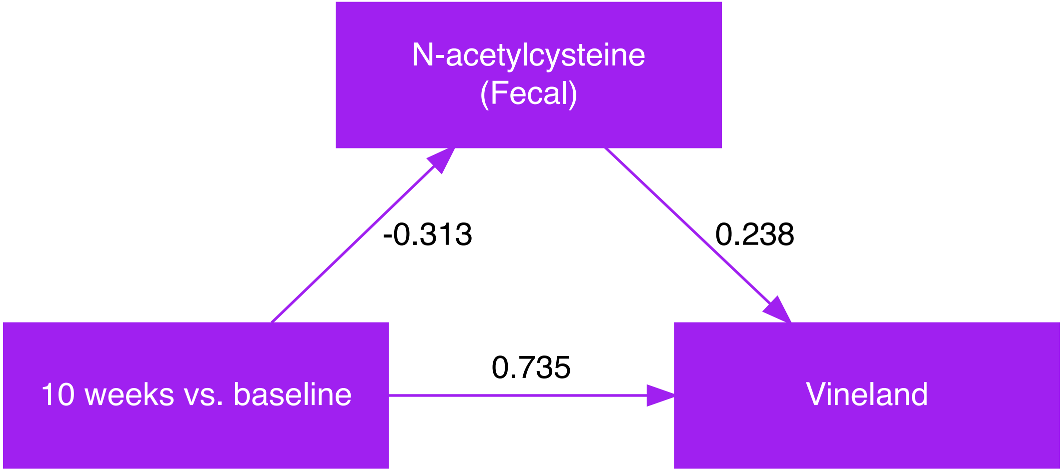
N-acetylcysteine (Fecal)
-0.313
0.238
0.735
10 weeks vs. baseline
Vineland

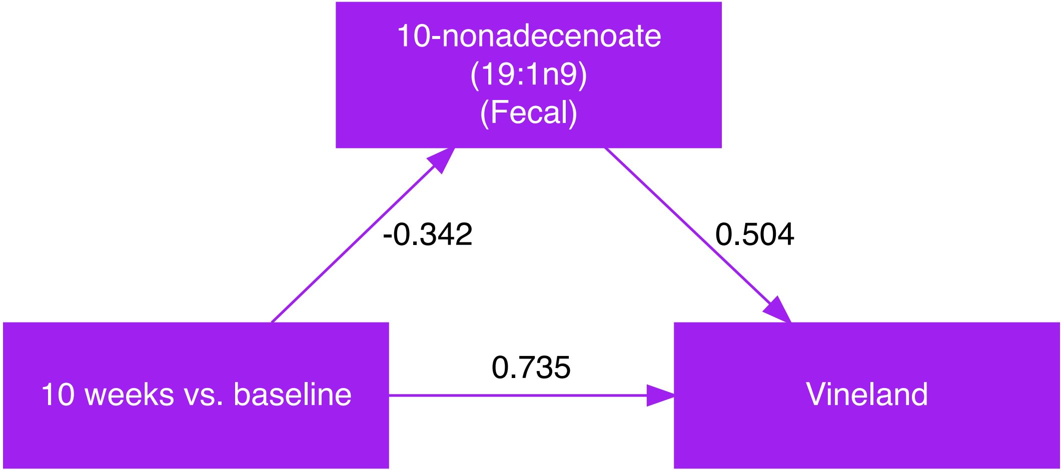
10-nonadecenoate
(19:1n9)
(Fecal)
-0.342
0.504
10 weeks vs. baseline
0.735
Vineland

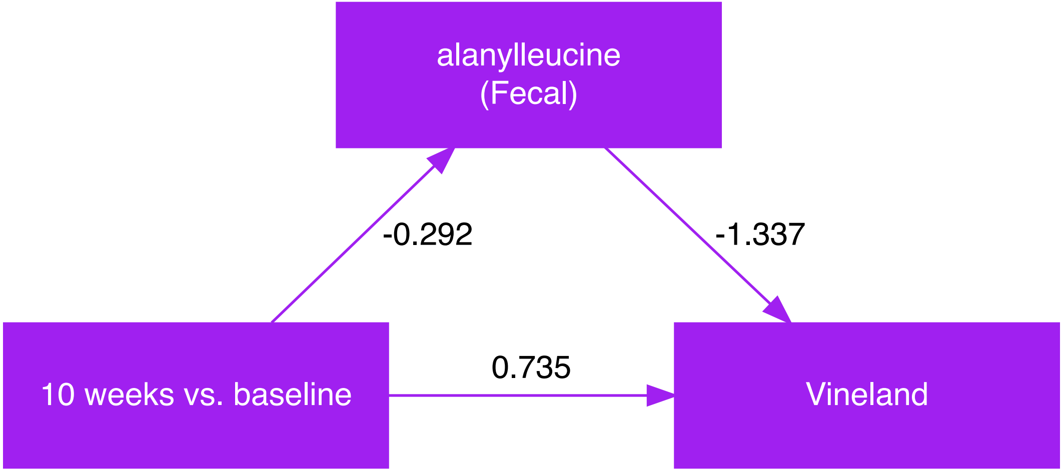
alanylleucine
(Fecal)
-0.292
-1.337
10 weeks vs. baseline
0.735
Vineland

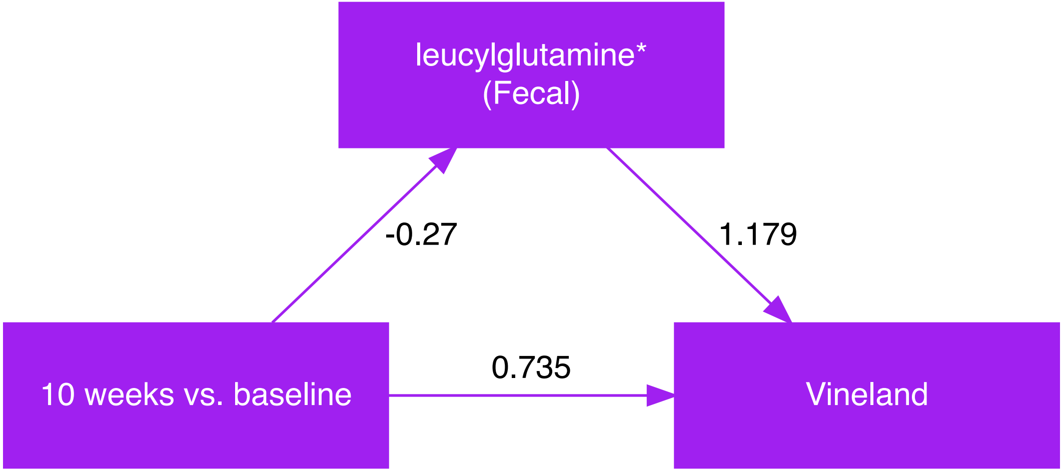
leucylglutamine*
(Fecal)
-0.27
1.179
0.735
10 weeks vs. baseline
Vineland

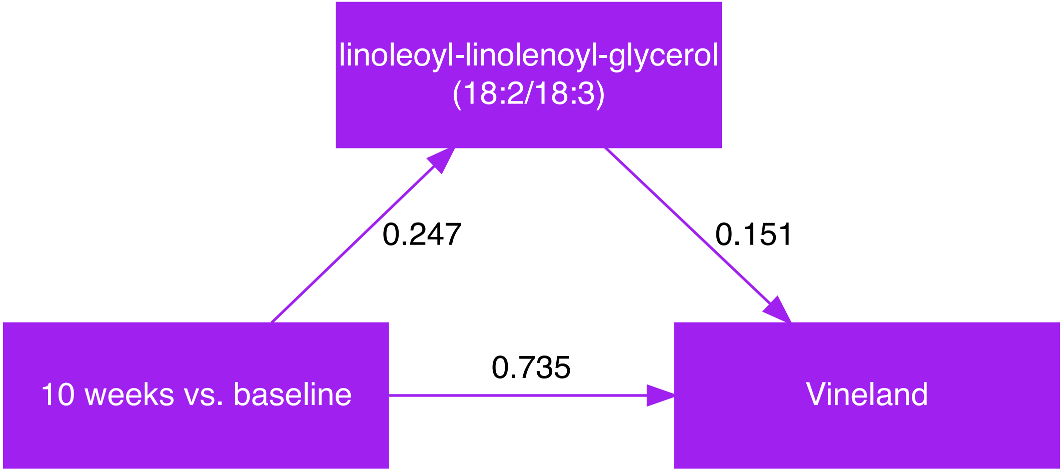
linoleoyl-linolenoyl-glycerol
(18:2/18:3)
0.247
0.151
0.735
10 weeks vs. baseline
Vineland

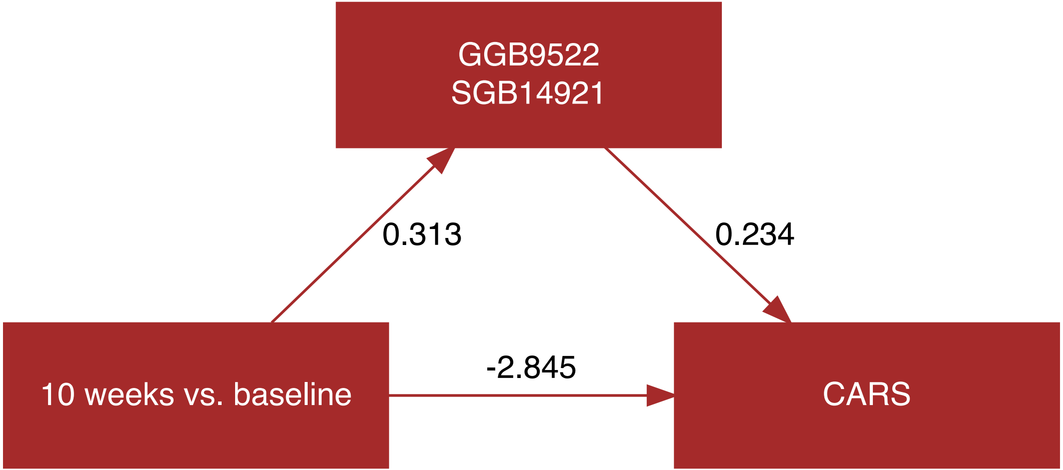
GGB9522
SGB14921
0.313
0.234
-2.845
10 weeks vs. baseline
CARS

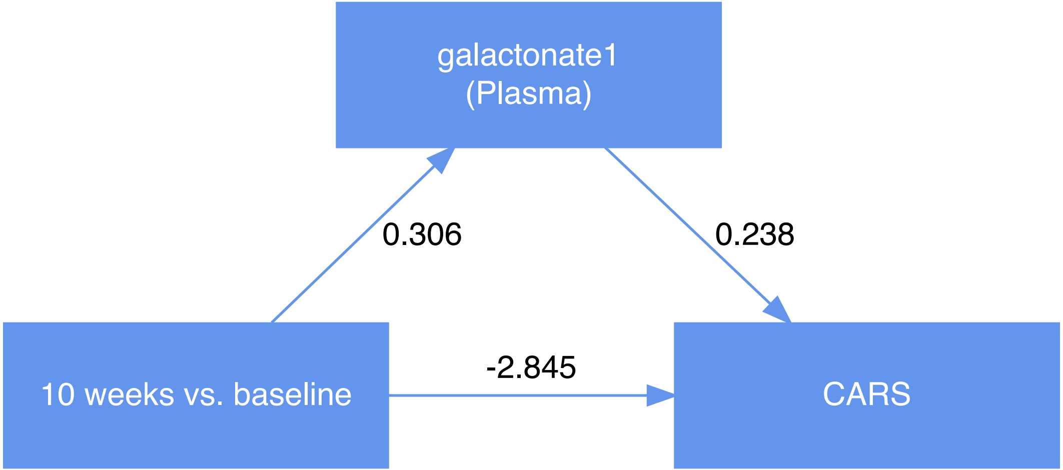
galactonate1
(Plasma)
0.306
0.238
10 weeks vs. baseline
-2.845
CARS

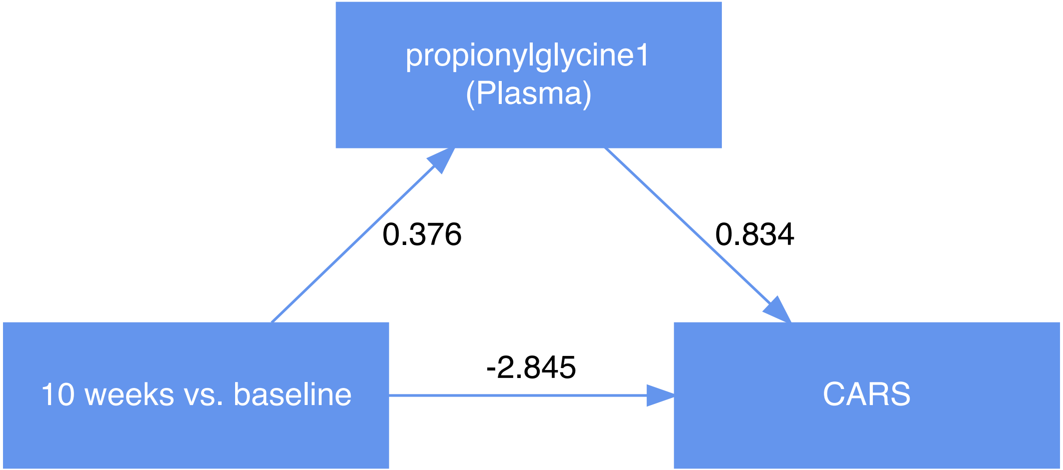
propionylglycine1
(Plasma)
0.376
0.834
-2.845
10 weeks vs. baseline
CARS

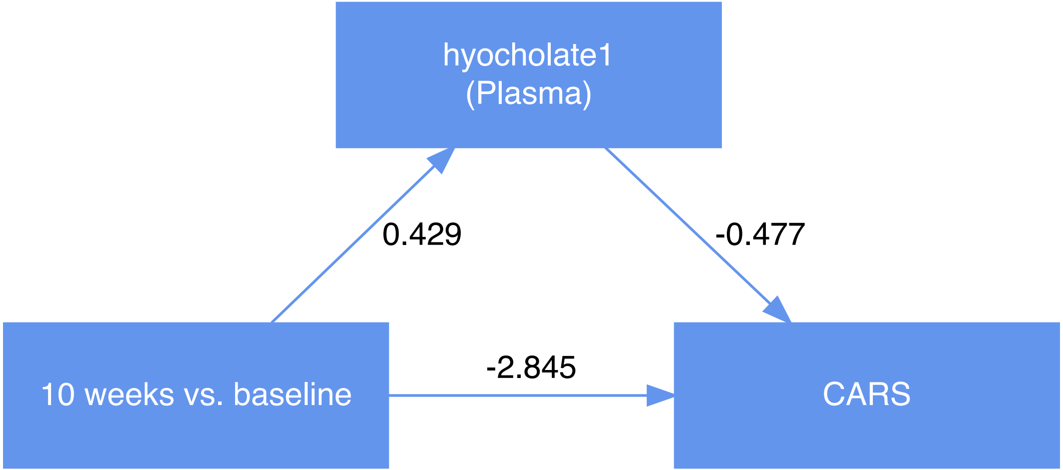
hyocholate1
(Plasma)
0.429
-0.477
10 weeks vs. baseline
-2.845
CARS

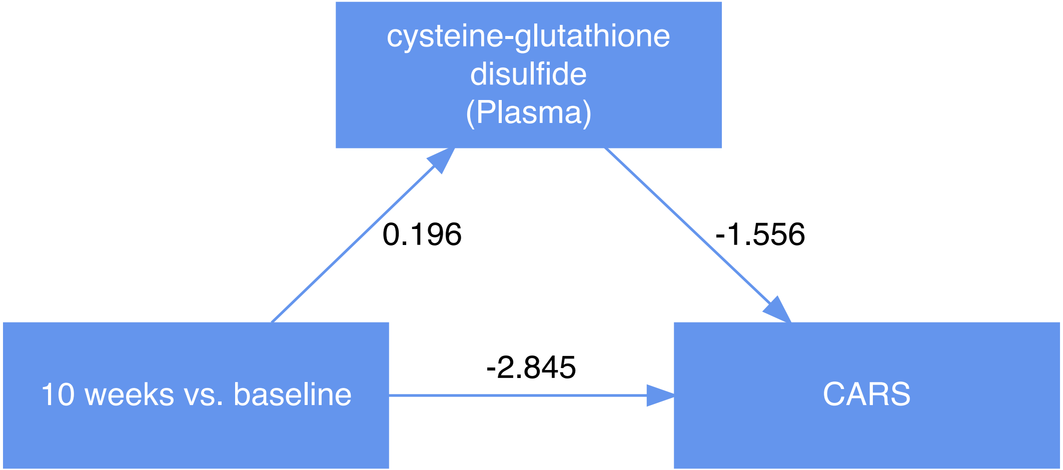
cysteine-glutathione disulfide (Plasma)
0.196
-1.556
-2.845
10 weeks vs. baseline
CARS

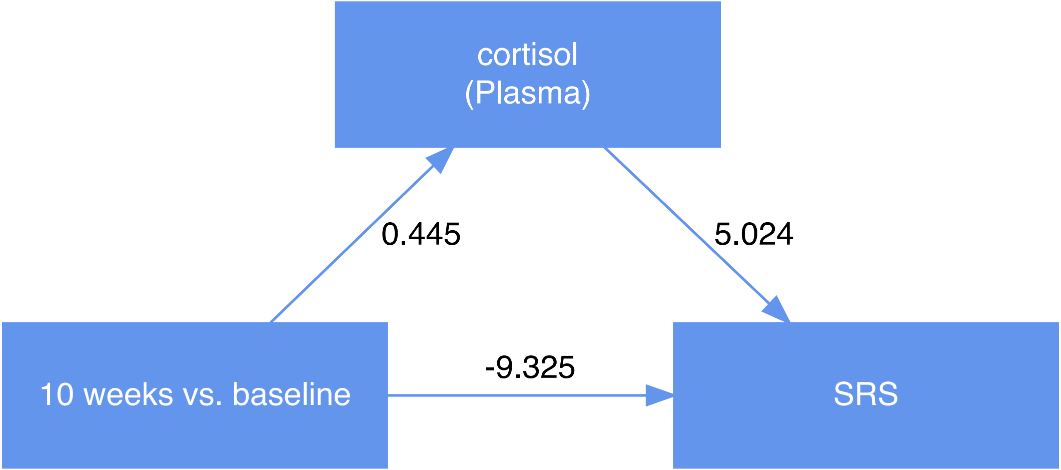
cortisol
(Plasma)
0.445
5.024
-9.325
10 weeks vs. baseline
SRS

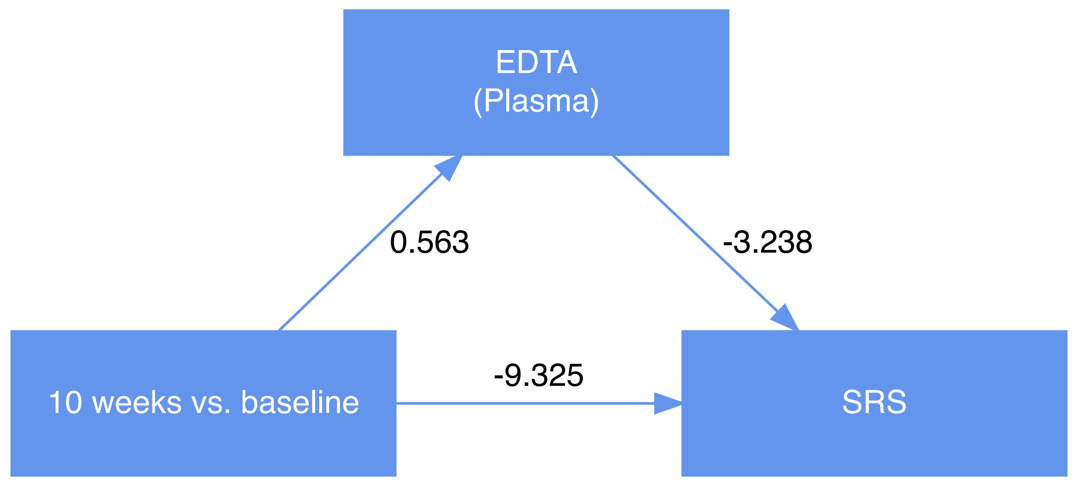
EDTA
(Plasma)
0.563
-3.238
-9.325
10 weeks vs. baseline
SRS

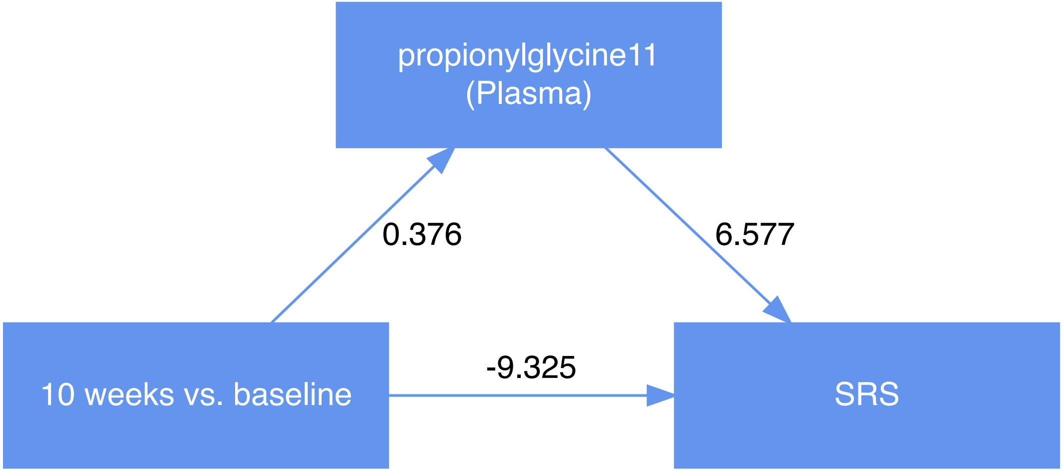
propionylglycine11
(Plasma)
0.376
6.577
-9.325
10 weeks vs. baseline
SRS

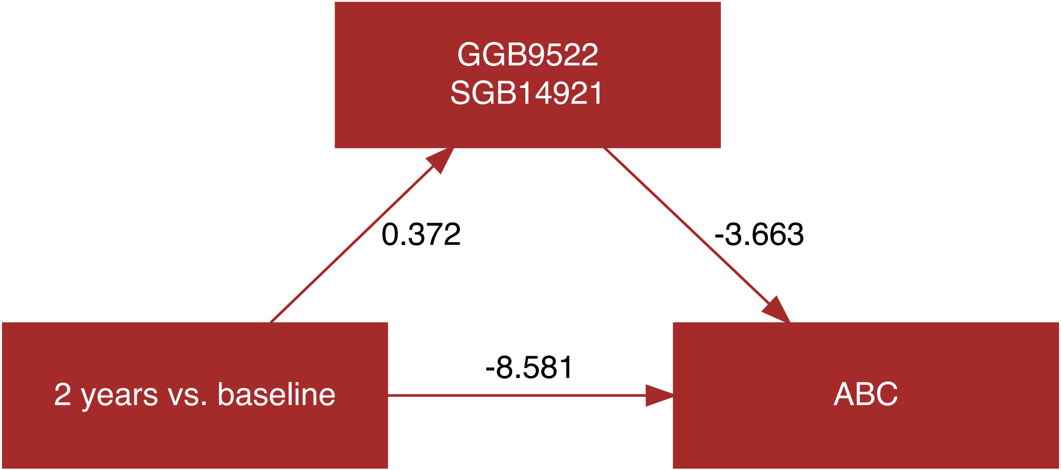
GGB9522
SGB14921
0.372
-3.663
-8.581
2 years vs. baseline
ABC

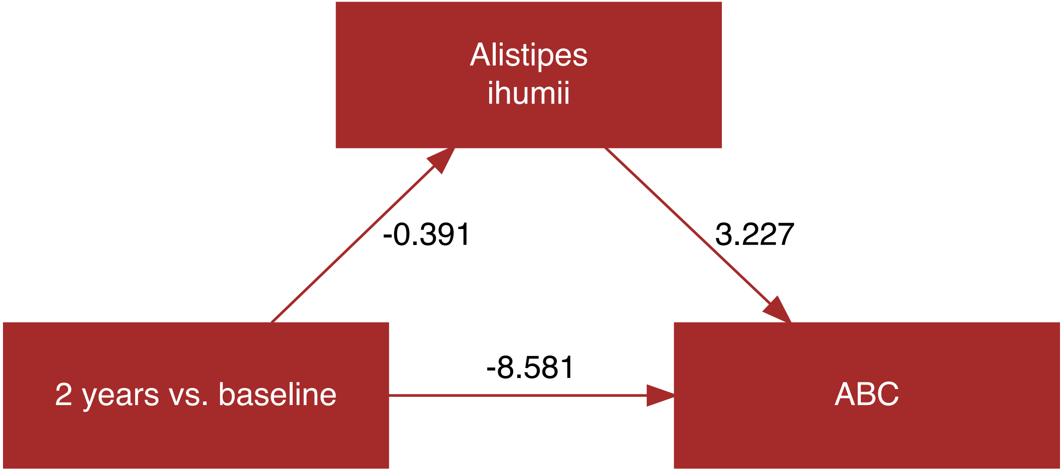
Alistipes
ihumii
-0.391
3.227
-8.581
2 years vs. baseline
ABC

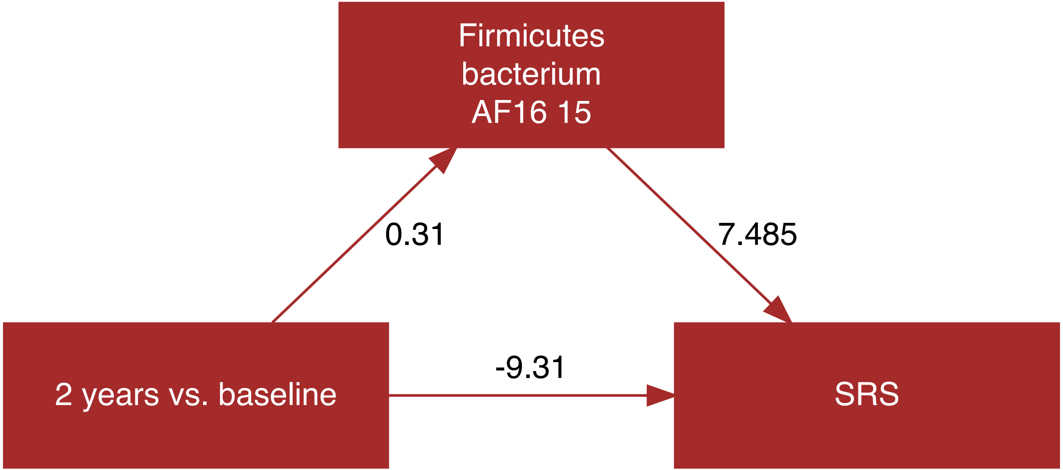
Firmicutes
bacterium
AF16 15
0.31
7.485
2 years vs. baseline
-9.31
SRS

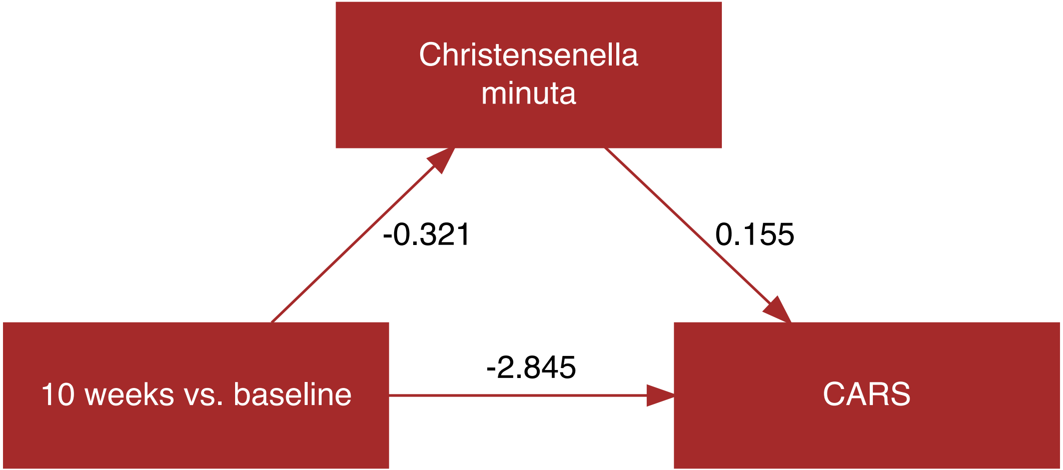
Christensenella minuta
-0.321
0.155
-2.845
10 weeks vs. baseline
CARS

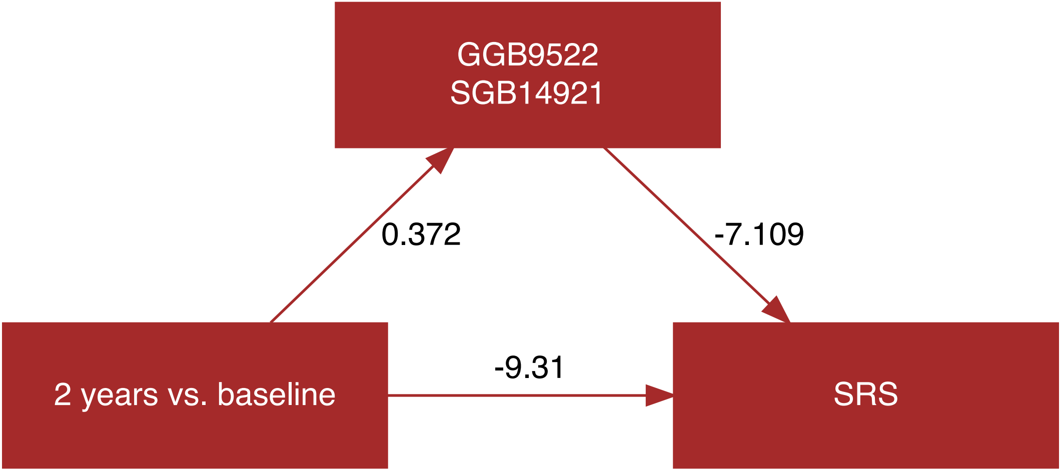
GGB9522
SGB14921
0.372
-7.109
-9.31
2 years vs. baseline
SRS

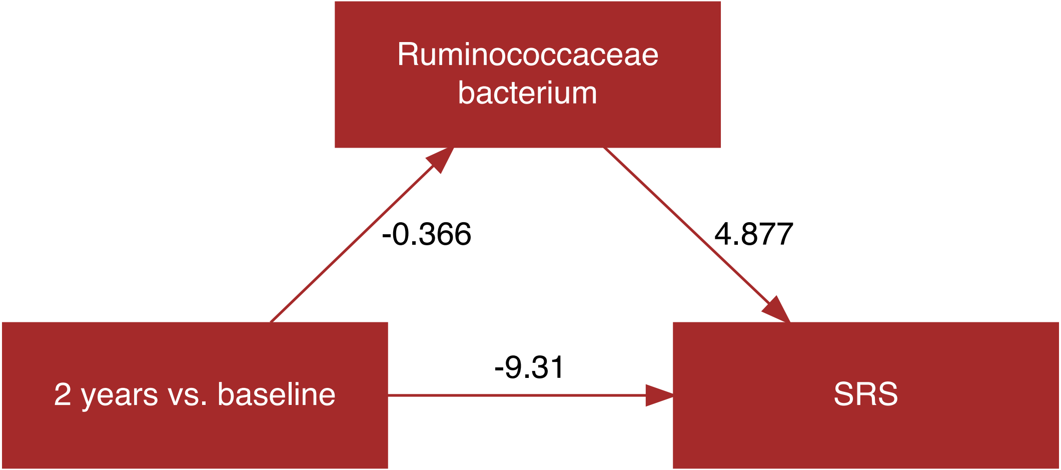

Ruminococcaceae
bacterium
-0.366
4.877
2 years vs. baseline
-9.31
SRS

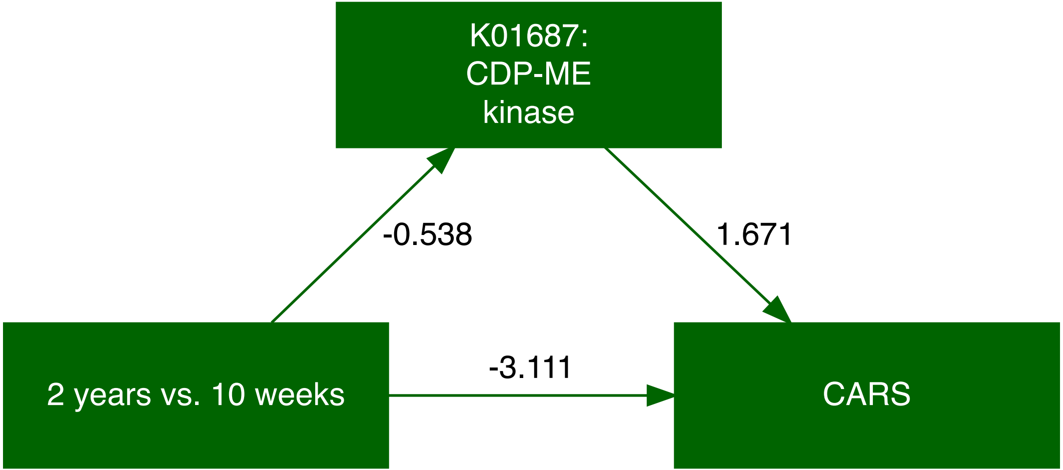
K01687:
CDP-ME
kinase
-0.538
1.671
2 years vs. 10 weeks
-3.111
CARS

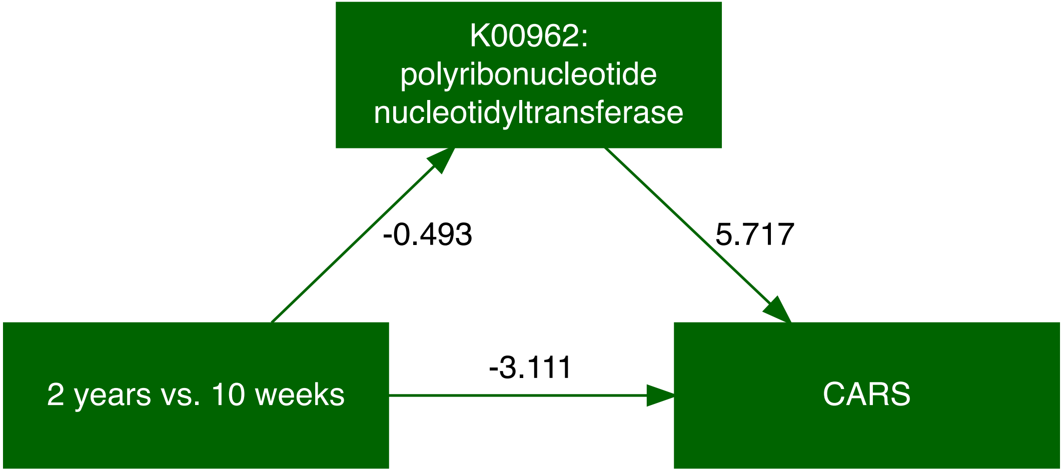
K00962:
polyribonucleotide
nucleotidyltransferase
-0.493
5.717
-3.111
2 years vs. 10 weeks
CARS

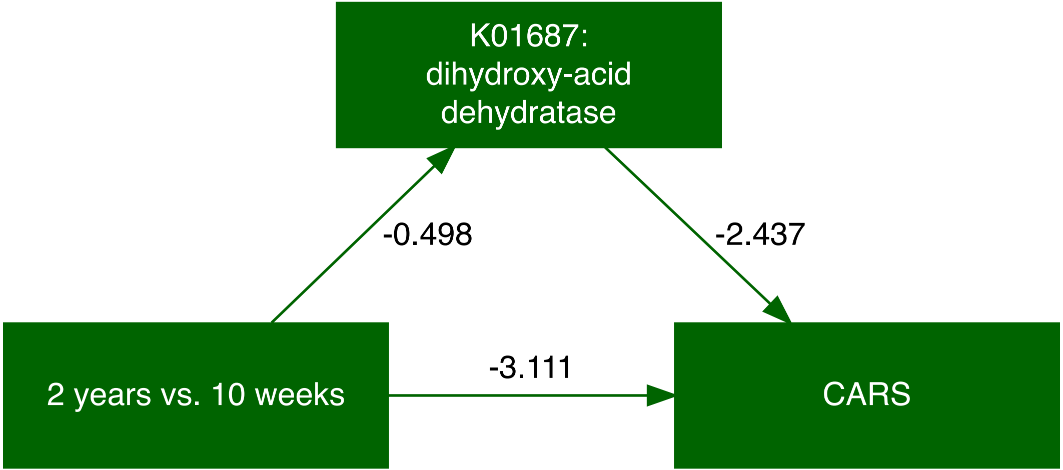
K01687:
dihydroxy-acid
dehydratase
-0.498
-2.437
-3.111
2 years vs. 10 weeks
CARS

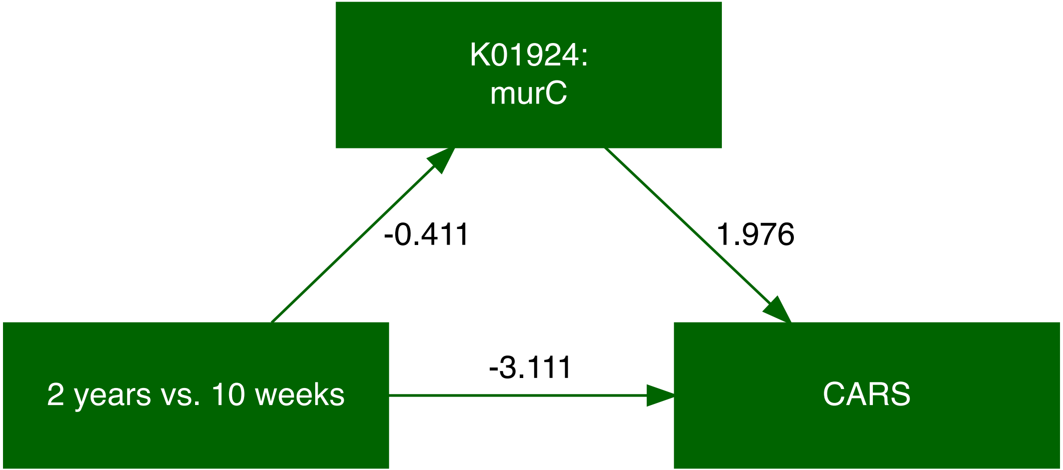
K01924:
murC
-0.411
1.976
-3.111
2 years vs. 10 weeks
CARS

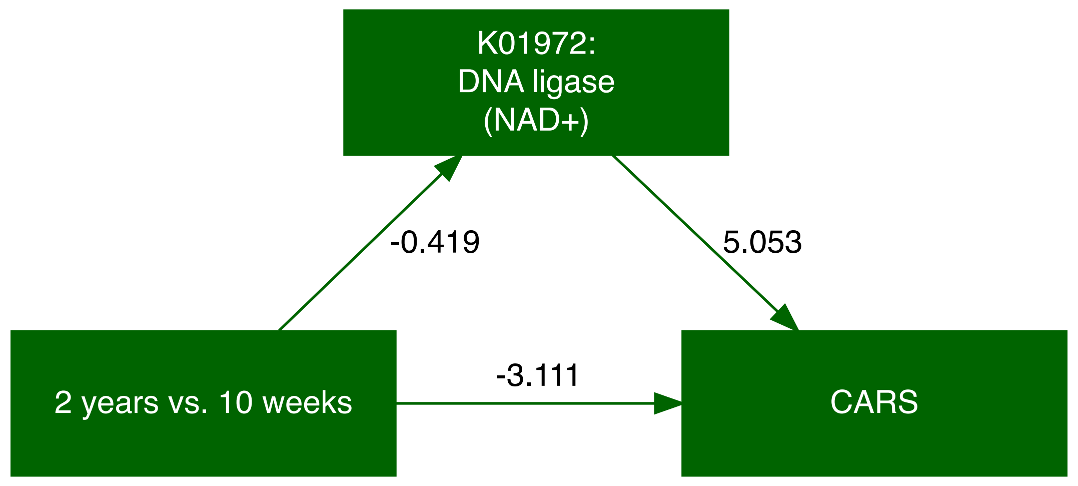
K01972:
DNA ligase
(NAD+)
-0.419
5.053
2 years vs. 10 weeks
-3.111
CARS

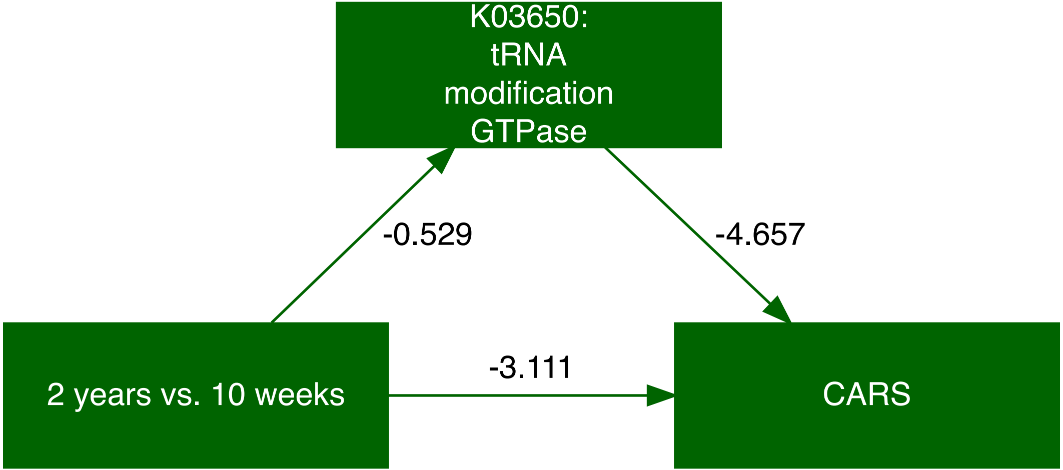
K03650:
tRNA
modification
GTPase
-0.529
-4.657
-3.111
2 years vs. 10 weeks
CARS

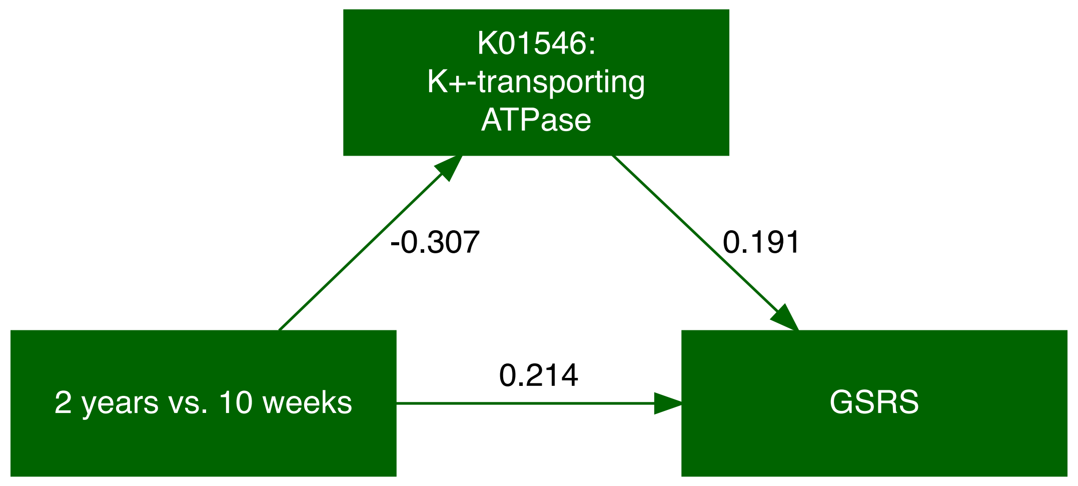
K01546:
K+-transporting
ATPase
-0.307
0.191
0.214
2 years vs. 10 weeks
GSRS

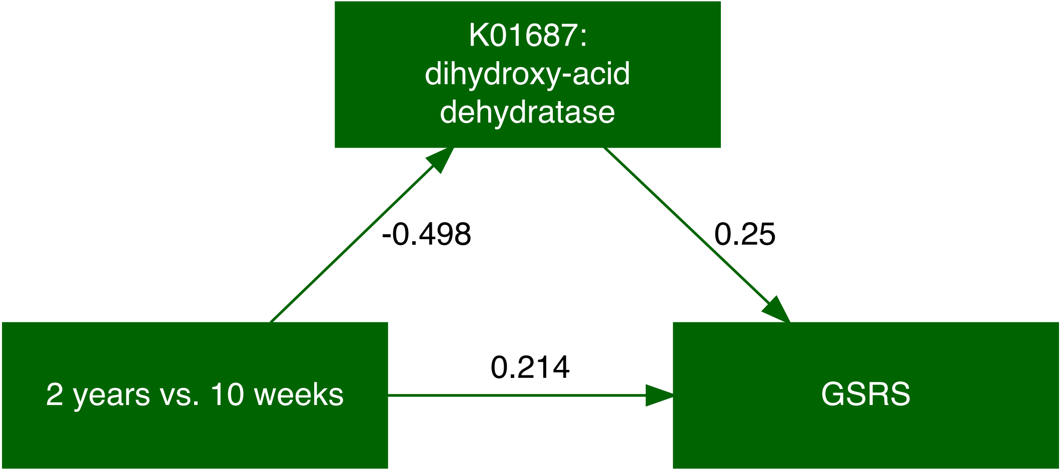
K01687:
dihydroxy-acid
dehydratase
-0.498
0.25
0.214
2 years vs. 10 weeks
GSRS

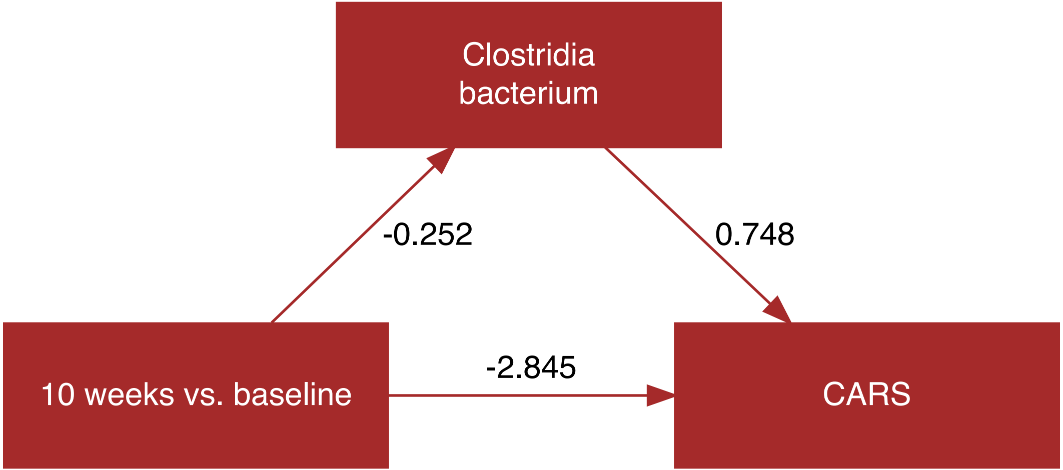
Clostridia
bacterium
-0.252
0.748
10 weeks vs. baseline
-2.845
CARS

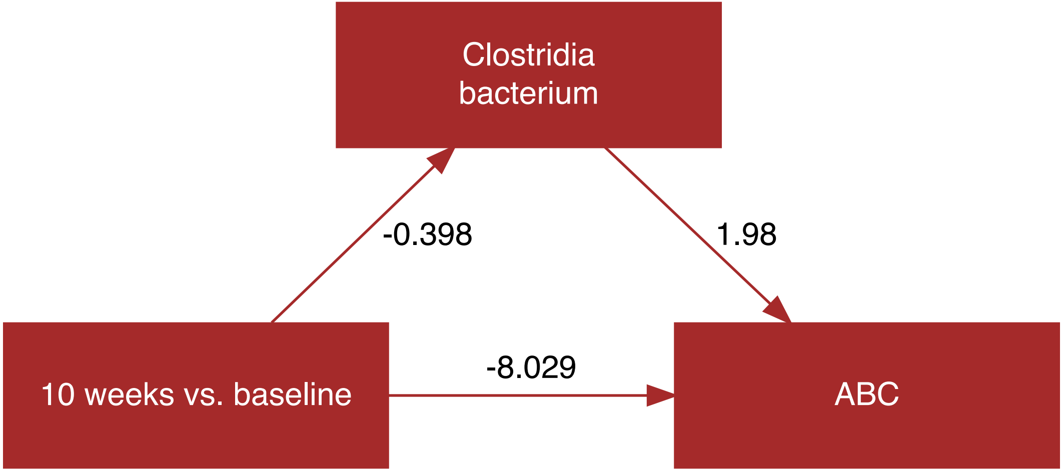
Clostridia
bacterium
-0.398
1.98
-8.029
10 weeks vs. baseline
ABC

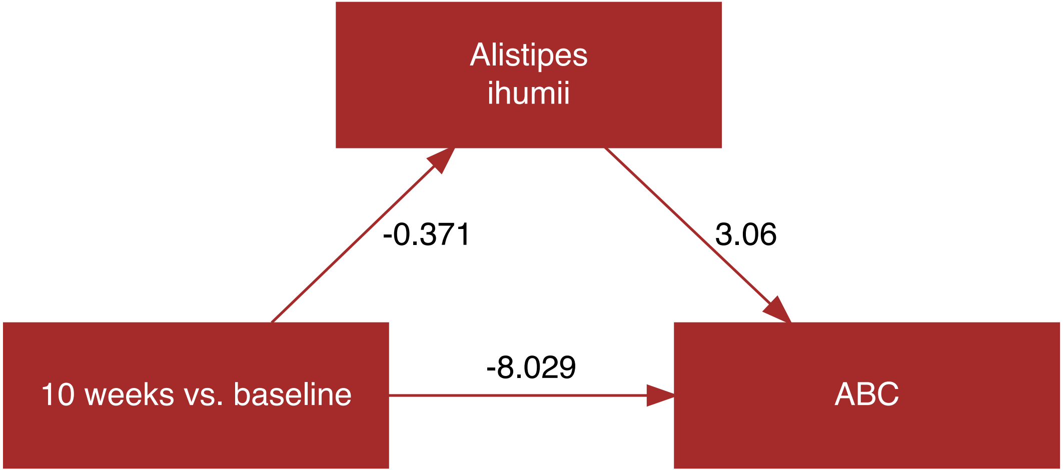
Alistipes ihumii
-0.371
3.06
-8.029
10 weeks vs. baseline
ABC

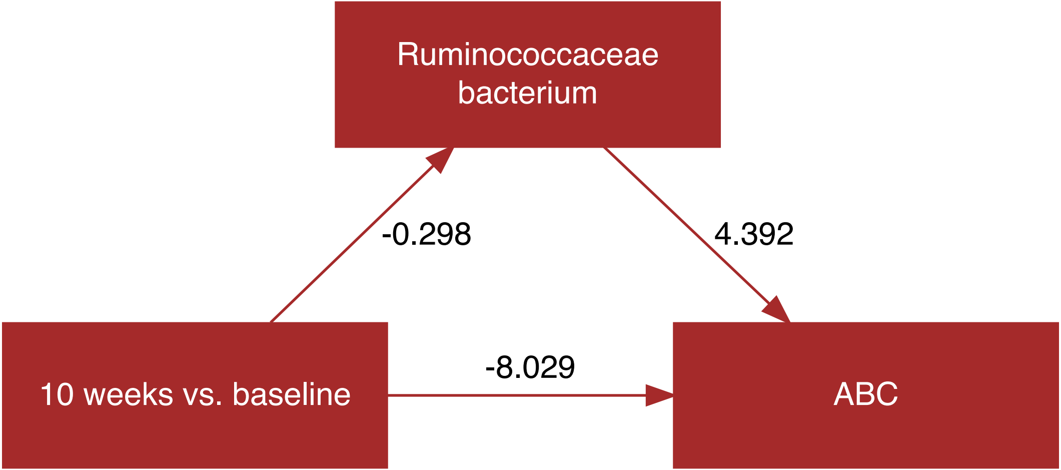
Ruminococcaceae bacterium
-0.298
4.392
10 weeks vs. baseline
-8.029
ABC
